# Impact of analytic decisions on test-retest reliability of individual and group estimates in functional magnetic resonance imaging: a multiverse analysis using the monetary incentive delay task

**DOI:** 10.1101/2024.03.19.585755

**Authors:** Michael I. Demidenko, Jeanette A. Mumford, Russell A. Poldrack

## Abstract

Empirical studies reporting low test-retest reliability of individual blood oxygen-level dependent (BOLD) signal estimates in functional magnetic resonance imaging (fMRI) data have resurrected interest among cognitive neuroscientists in methods that may improve reliability in fMRI. Over the last decade, several individual studies have reported that modeling decisions, such as smoothing, motion correction and contrast selection, may improve estimates of test-retest reliability of BOLD signal estimates. However, it remains an empirical question whether certain analytic decisions *consistently* improve individual and group level reliability estimates in an fMRI task across multiple large, independent samples. This study used three independent samples (*N*s: 60, 81, 119) that collected the same task (Monetary Incentive Delay task) across two runs and two sessions to evaluate the effects of analytic decisions on the individual (intraclass correlation coefficient [ICC(3,1)]) and group (Jaccard/Spearman *rho*) reliability estimates of BOLD activity of task fMRI data. The analytic decisions in this study vary across four categories: smoothing kernel (five options), motion correction (four options), task parameterizing (three options) and task contrasts (four options), totaling 240 different pipeline permutations. Across all 240 pipelines, the median ICC estimates are consistently low, with a maximum median ICC estimate of .43 - .55 across the three samples. The analytic decisions with the greatest impact on the median ICC and group similarity estimates are the *Implicit Baseline* contrast, Cue Model parameterization and a larger smoothing kernel. Using an *Implicit Baseline* in a contrast condition meaningfully increased group similarity and ICC estimates as compared to using the *Neutral* cue. This effect was largest for the Cue Model parameterization; however, improvements in reliability came at the cost of interpretability. This study illustrates that estimates of reliability in the MID task are consistently low and variable at small samples, and a higher test-retest reliability may not always improve interpretability of the estimated BOLD signal.

## Introduction

Reliability in functional magnetic resonance imaging (fMRI) is essential to individual differences research as well as for the development of clinical biomarkers. Unfortunately, numerous studies have demonstrated that reliability of individual estimates in fMRI is low (Elliott et al., 2020; Noble et al., 2019) and the reliability of group estimates in statistical maps is sensitive to varying analytical decisions made by researchers (Botvinik-Nezer et al., 2020)^1^. Poor reliability can hamper validity in cognitive neuroscience research, reducing the ability to uncover brain-behavior effects (Hedge et al., 2018; Nikolaidis et al., 2022) and the ability to detect differences in distinct brain states and individual traits (Gell et al., 2023; Kragel et al., 2021). It remains to be seen whether certain analytic decisions *consistently* reduce individual and/or group reliability estimates of blood oxygen-level dependent (BOLD) activity across measurement occasions in univariate task fMRI analyses.

FMRI analysis involves a range of analytic decisions (Caballero-Gaudes & Reynolds, 2017; Soares et al., 2016) that can result in a vast number of statistical brain maps across which BOLD activity can vary subtly or substantially (Bowring et al., 2022; Carp, 2012; Li et al., 2021). Simple decisions, such as using different MNI template brains, can greatly affect the agreement between parameter estimates between two preprocessing pipelines (Li et al., 2021). Furthermore, the approach used to model a task design can also alter interpretations (Botvinik-Nezer et al., 2020). As a result of numerous arbitrary choices, preprocessing and task modeling decisions can significantly impact the reliability of voxel/region of interest (ROI) estimates (Dubois & Adolphs, 2016).

Different metrics of reliability provide quantitative indices of the consistency (or similarity) of estimates of BOLD activity in specific brain regions (or voxels) during fMRI task activation across repeated measurement occasions (Bennett & Miller, 2013). Researchers can quantify the consistency of two repeated measures in terms of estimated effects (continuous) and/or the presence/absence of a significant effect (binary). In terms of the continuous effects, reliability is an estimate of the consistency of the numerical representation of a measure (e.g., BOLD activity in the supplementary motor area during a finger tapping task [Witt et al., 2008]) of a mental process (e.g., index finger movement) across repeated measurement occasions within an *individual* (e.g., task fMRI contrasts across two or more sessions, which can be hours, days or weeks). This form of reliability is usually calculated using an intraclass correlation (ICC) at the whole brain (i.e., voxel-wise) and/or ROI level. In terms of binary estimates of an effect, reliability is an estimate of an experimental task’s (e.g., finger tapping task [Witt et al., 2008]) ability to evoke statistically significant activation (above a pre-specified threshold) in the same regions for *groups* of subjects for a specific condition (e.g., finger movement versus rest) across measurement occasions (e.g., task fMRI contrasts across two or more scanning sessions). Binary estimates of reliability are often calculated using Dice (Rombouts et al., 1998) or Jaccard’s similarity coefficients (Maitra, 2010). Together, these two forms of reliability reflect the consistency (or agreement) in either the magnitude or the binary statistical significance of an experimental effect occurring during task fMRI.

Traditionally, empirical studies have referred to the “robustness” of above-threshold activation signals in group fMRI analyses as an implicit indicator of reliability of an fMRI task. While a useful heuristic, Fröhner et al. (2019) argued that robustness across measurement occasions only represents reliability of *group* (overall average) BOLD activity and does not accurately represent *individual* variability in BOLD activity. In addition, thresholding is a nonlinear operation that can result in substantial variability (Cohen & DuBois, 1999). When quantifying reliability of BOLD activity in the brain, researchers often report an ICC or a similarity coefficient for task fMRI (Bennett & Miller, 2013; Fröhner et al., 2019). The lack of standardization makes it challenging to precisely quantify reliability, relative to individual differences, and assess the impact of different fMRI analysis decisions on continuous and binary estimates of reliability.

To date, several studies have examined the impact of analytic decisions, such as spatial smoothing, motion correction and contrast modeling, on individual estimates of reliability of task fMRI. Caceres et al. (2009, *n* = 10) found that an optimal smoothing kernel size of 8-10 FWHM (full-width half-maximum) on a 1.5T scanner with 3.75mm voxels improved reliability. Results regarding the impact of motion correction on reliability are mixed, with Gorgolewski et al. (2013, *n* = 11) reporting a positive effect on reliability while Plichta et al. (2012, *n* = 25) reporting no effect during a reward task and a negative effect during a faces and N-back task on reliability. However, in a large, young sample, Kennedy et al. (2022, *n* = 5,979 - 6,593) reported that excluding high motion subjects modestly improved reliability. Finally, Han et al. (2022, *n* = 29 - 120) and Kennedy et al. (2022, *n* = 5,979 - 6,593) reported that using an implicit baseline for different tasks (e.g., rest phase during the task) rather than a neutral cue increased reliability across measurement occasions. Some, but not all, of these findings are consistent with a previous review of the fMRI reliability literature (Bennett & Miller, 2013), which suggests that motion, spatial smoothing and task signal likely impacts reliability in task fMRI. However, differences in modeling decisions across these studies leaves an important question unanswered: Are there certain analytic decisions that *consistently* improve reliability (e.g., ICC) of neural activity for an fMRI task across samples?

The ICC is a statistic adopted from behavioral research to estimate reliability of observed scores across measurement occasions (Bartko, 1966; Fisher, 1934; Shrout & Fleiss, 1979; Spearman, 1904). In the context of multi-session data, there are several ways to estimate an ICC, but for typical univariate fMRI studies, two specific types (ICC[2,1] and ICC[3,1]) are recommended (For a discussion, see Noble et al., 2021). As described elsewhere (Bennett & Miller, 2013; Fisher, 1934), the ICC is similar to the product moment correlation. Unlike the product moment correlation, which estimates separate means and variances between distinct classes (e.g., age and height), the ICC estimates the mean and variances within a single class (e.g., measure). For two or more variables from a single class, test-retest reliability estimates the consistency (or agreement) of the observed scores across the measurement occasions. Using the correlation coefficient as an example, if there are no differences in subjects’ scores across two measurement occasions, the correlation coefficient would be 1.0. However, if the measure is affected by systematic and/or unsystematic error across measurement occasions, this would impact the covariance between observed scores across subjects and decrease the linear association between measures across the two occasions. Unlike the product moment correlation, however, the ICC factors out measurement bias which reflects the reproducibility of observed scores across measurement occasions (Liu et al., 2016). While the correlation between two occasions (**A** = [1, 3, 6, 9, 12] & **B** = 3x**A** = [3, 9, 18, 27, 36]) may be perfect (*r*_AB_ = 1.0), the consistency in observed scores between the two measurement occasions would be lower (ICC[3,1] = .60). In fMRI, the reliability of the BOLD signal may be impacted by biological (e.g., differences in BOLD across brain region), analytic (e.g., task design and analytic decisions), and participant-level factors (e.g., practice effects, motion, habituation and/or development). These fluctuations, whether typical or atypical, may contribute to observed differences and the reduced consistency in scores across measurement occasions, leading to decreased estimates of reliability.

As discussed in prior work on fMRI reliability (Bennett & Miller, 2010, 2013; Caceres et al., 2009; Chen et al., 2017; Herting et al., 2017; Noble et al., 2021), the ICC decomposes the total variance of the data across all subjects and sessions into two key parts: *Between-subject* and *Within-subject* variance (for statistical formulas and discussion of ICC, see Liljequist et al., [2019] and flowchart in McGraw & Wong [1996, p. 40]). The ICC estimate can be altered by increasing the differences in BOLD activity between subjects (e.g., subjects differ more in BOLD activity in index finger movements) and/or ensure that BOLD activity within subjects is more similar across scans (e.g., BOLD activity in response to finger movements versus rest for Subject A is consistent across Session 1 and Session 2). Some have argued that the low *between-subject* variability may be a reason for low reliability of behavioral responses in experimental tasks that are commonly used in fMRI (Hedge et al., 2018). However, there is little empirical research on whether the culprit in the reportedly low reliability of fMRI signal across measurement occasions is a *decreased between-subject* and/or an *increased within-subject* variability. It also remains an open question whether certain analytic decisions differentially impact the between/within subject variance and consistently improve reliability across different samples with the same task. As it relates to prediction and global signal-to-noise ratio, evidence from Churchill et al. (2015; *n* = 25) suggest that there are likely to be optimal preprocessing pipelines; however, the degree to which these differ across datasets and individuals is currently unknown.

The current study uses a multiverse (Steegen et al., 2016) of analytic alternatives to simultaneously evaluate the effects of analytic decisions on the continuous and binary reliability estimates of neural activity in task fMRI in three samples. The three samples administered with the comparable Monetary Incentive Delay (MID) task during fMRI across two runs and two sessions. The purpose of multiple samples with the same task design is to evaluate the consistency in findings across studies that vary in their sample populations and task design as little evidence exists on the *consistency* of reliability estimates for the same task across independent samples. **Aim 1** evaluates the effects of analytic decisions including task model smoothing, motion correction, parameterization (i.e., modeling) and task contrasts on the impacts on reliability, calculated using ICC(3,1) for individual [continuous] beta estimates and Jaccard’s similarity coefficient using significance thresholded group [binary] estimates (*p* < .001, uncorrected) and Spearman correlation group [continuous] estimates. The decisions are noted in **Table 1**. **Aim 1 Hypothesis** is that the highest produced ICC and similarity coefficient/correlation is for the model decisions indicated by **blue** for A-D decisions in Table 1. This, in part, is because the analytic strategy includes 1) motion correction techniques that limit the number of noisy (high motion) subjects and reduce the number of degrees of freedom that are lost due to censoring, 2) an optimal smoothing for the size of voxels, and 3) the highest activation contrast from a task modeling phase that is relatively efficient. We hypothesize this to be more so the case for the older (e.g., AHRB/MLS) than younger samples (e.g., ABCD) due to changes occurring as a result of development (Herting et al., 2017; Noble et al., 2021). Due to the lack of information regarding how the between-subject variance (BS) and within-subject variance (WS) is impacted by analytic choices in task fMRI analyses, **Aim 2** evaluates the change in BS and WS components. Due to the poor reliability of individual estimates in task fMRI (Elliott et al., 2020), reported evidence of high between-subject variability in BOLD activity (Turner et al., 2018), and limited evidence on changes in BS and WS variance components in the MID task, we do not have a specific **Aim 2 Hypothesis**. Finally, seeing as the ICC is, in some ways, similar to a moment product correlation (Bennett & Miller, 2010) which stabilizes at larger sample sizes (Grady et al., 2020; Marek et al., 2022; Schönbrodt & Perugini, 2013), **Aim 3** evaluates at what sample the ICC stabilizes using the most optimal pipeline (e.g., highest median ICC) used in Aim 2. Stability of Jaccard coefficient group maps is not considered in Aim 3 as these estimates are sensitive to significance thresholding. Using the evidence from prior work on correlations (Grady et al., 2020; Schönbrodt & Perugini, 2013), the **Aim 3 Hypothesis** is that the ICC will stabilize a sample size between 150 to 500.

**Table 1.** Proposed Analytic Permutations: 360 Total Modeling Combinations for MID task

| First-level Pipeline Decisions | Options |
| --- | --- |
| **A. Smoothing (FWHM)** |  |
| 1. 1.5x voxel | ON / OFF |
| 2. 2x voxel | ON / OFF |
| 3. 2.5x voxel | ON / OFF |
| 4. 3x voxel | ON / OFF |
| 5. 3.5x voxel | ON / OFF |
| **B. Motion Correction** |  |
| 1. None | ON / OFF |
| 2. Regress: Translation/Rotation (x,y,z) + Derivative (x,y,z) | ON / OFF |
| 3. Regress: Regress: Translation/Rotation (x,y,z) + Derivative (x,y,z) + First 8 aCompCor Components | ON / OFF |
| 4. Regress: Translation/Rotation (x,y,z) + Derivative (x,y,z) + First 8 aCompCor Components + Censor High Motion Volumes (FD ≥ .9) | ON / OFF |
| #5. Regress: Translation/Rotation (x,y,z) + Derivative (x,y,z) + First 8 aCompCor Components, Exclude mean FD ≥ .9 | ON / OFF |
| #6. Regress: Translation/Rotation (x,y,z) + Derivative (x,y,z) + First 8 aCompCor Components + Censor High Motion Volumes, Exclude mean FD ≥ .9 | ON / OFF |
| **C. Task Modeling** |  |
| 1. MID: Cue Onset, Cue Duration only | ON / OFF |
| 2. MID: Cue Onset, Cue + Fixation Duration | ON / OFF |
| 3. MID: Fixation onset, Fixation Duration | ON / OFF |
| **D. Task Contrasts** |  |
| 1. MID: Big Win > Neutral | ON / OFF |
| 2. MID: Big Win > Implicit | ON / OFF |
| 3. MID: Small Win > Neutral | ON / OFF |
| 4. MID: Small Win > Implicit | ON / OFF |

Blue text: Model hypothesized to produce the highest test-retest reliability; aCompCor: Anatomical Component Based Noise Correction; MID: Monetary Incentive Delay task; FD: Framewise displacement.

## Methods

To answer the questions proposed in Aim 1 and Aim 2, this study will require multiple samples and tasks to obtain a comprehensive view of how analytic decisions impact group and individual reliability metrics (Aim 1) and how BS and WS is impacted (Aim 2) across multiple samples and similar MID task. We use three samples with subjects that have at least two repeated sessions of data. To answer the question about the sample at which ICC stabilizes (Aim 3), we use the repeated session data from a large consortium sample.

The studies were selected based on two criteria. First, the goal is to derive group and individual estimates of reliability using sample sizes that are larger than the reported median sample size in fMRI research. The median reported sample size in fMRI is <30 subjects (Poldrack et al., 2017; Szucs & Ioannidis, 2017). From the review of task fMRI reliability by Bennet and Miller (2010), the median sample for individual (continuous) reliability is 10 subjects (mean = 10.5 [range = 1 to 26]) and for group (binary) reliability is 9.5 subjects (mean = 11.2 [range = 4 to 45]). A recent review and analysis of task fMRI reliability suggests sample sizes are increasing but remain lower than the median sample size in task fMRI, whereby the median sample size for individual reliability in the meta-analysis are 18 subjects (mean = 26.4 [range = 5 to 467]) and the analyses are 45 & 20 subjects (Elliott et al., 2020). Second, the goal is to limit the interaction between reliability estimates and unknown features of the data, such as the mental processes, to get a sense of how the analytic pipeline impacts reliability estimates *consistently* across a similar task design. Thus, the three samples described below exceed N > 50 and use a nearly identical task that is known to evoke a strong BOLD response in specific brain regions to achieve these two goals.

### Participants^2^

#### Adolescent Brain Cognitive Development (ABCD) Study

The ABCD Study® is a longitudinal national study that was designed to study the change in behavioral and biological measurements across development (Volkow et al., 2018). The focus here is on the 4.0 brain imaging data that is released by the ABCD-BIDS Community Collection (ABCC; Feczko et al. [2021]). As of February 2024, the ABCC data contains year 1 (approximately 11,000, participants Aged 9-10) and year 2 (approximately 7,000 participants, Age 11-13) fMRI data. For Aims 1 and 2, we use a subsample of ABCD participants at the University of Michigan site (site = 13) with maximum clean data available as this would be sufficient to test the hypotheses and limit site and scanner effects. For Aim 3, we use a subsample of N = 2,000 of the maximum clean data available from the ABCC sample and use an adaptive design to answer at which *N* ICC stabilizes. To reduce the use of unnecessary computational resources, the analyses are first performed in N = 525. If the difference between average ICC estimate for interval N*_i_*& N*_i-1_*is > .15, the sample will be extended to *N* = 1000, adding *N* = 500, until the plotted estimates are stable. As described elsewhere (Casey et al., 2018), the study collected fMRI data during the Stopsignal, Emotional N-back and MID tasks.

Reliability of consortium-derived region of interest level data for year 1 and year 2 has been reported elsewhere (Kennedy et al., 2022). We expand on these findings by evaluating how consistent these results are across studies and which analytic decisions impact estimates of reliability. Here, we use the raw BOLD timeseries from the MID task as this is consistent with the two other studies described below.

#### Michigan Longitudinal Study (MLS)

The MLS is a longitudinal study focused on the change in behavioral and biological measurements across development. As described elsewhere (Martz et al., 2016; Zucker et al., 2000), the MLS includes the Neuropsychological Risk cohort. The MLS Neuropsychological Risk cohort contains year 1 (approximately 159 participants, Age 18-24) and year 2 (approximately 150 participants, Age 20-26) fMRI data. The study collected fMRI data during the affective word and MID tasks. Here, we use the raw BOLD data from the MID task as it is consistent with the ABCD study and Adolescent Risk Behavior Study (described below).

#### Adolescent Risk Behavior (AHRB) Study

The AHRB study is a longitudinal study focused on the change in behavioral and biological measurements across development. The AHRB study contains year 1 (approximately 108 participants, Age 17-20) and year 2 (approximately 66 participants, Age 19-22). The study collected fMRI data during the Emotional Faces and MID tasks. Here, we use the raw BOLD data from the MID task as it is consistent with the MLS and AHRB study.

### FMRI Task, Data, Preprocessing

#### FMRI Tasks

Across the ABCD, AHRB and MLS studies, reward processing was measured using comparable versions of the MID task. The MID task (Knutson et al., 2000) is used to model BOLD signatures of the anticipation and receipt of monetary gains or losses. The MID task and their nuanced differences across the ABCD, AHRB and MLS studies are described in supplemental **Section 1.2**. The focus of the present work is on the anticipatory phase of the task.

#### MRI Acquisition Details

The acquisition details for the AHRB, ABCD and MLS datasets are summarized in supplemental **Section 1.3 Table S2**.

#### Data Quality Control and Preprocessing

First, quantitative metrics reported from MRIQC version 23.1.0 (Esteban et al., 2023) for the structural and BOLD data are evaluated to assess data quality and potentially problematic subjects. Second, behavioral data were inspected to confirm that participants have the behavioral data for each run and that participants performed at the targeted probe hit rate (e.g., at or near 60% overall probe hit rate, see supplemental **Section 1.2**). Then, structural and functional MRI preprocessing is performed using fMRIPrep v23.1.4 (Esteban et al., 2022; RRID:SCR_016216), which is based on Nipype 1.8.3 (Esteban, Markiewicz, Burns, et al., 2022; RRID:SCR_002502) and the results are inspected to confirm no subjects’ preprocessing steps failed.

Preprocessing between the ABCD, AHRB and MLS are held constant except for two differences. First, the MLS datasets did not collect fieldmaps and the repetition time for MLS (2000ms) is slower than the repetition time (800ms) in ABCD/AHRB. Therefore, fMRIPrep’s fieldmap-less distortion correction (SyN-SDC) is used to estimate and correct for fieldmap distortions in MLS and slice-timing correction is applied *only* on the MLS data. For the ABCD and AHRB data, fieldmap-less distortion correction is used *only* when a subject does not have the necessary fieldmaps. Outside of these two exceptions, the preprocessing of the BIDS data were preprocessed using identical pipelines. The complete preprocessing details are included in supplemental **Section 1.4**

### Analyses

This project is focused on the effects of analytic decisions on estimates of reliability across (run/session) measurement occasions in task fMRI. As a reminder, reliability is the estimate of how similar two measures (in this case, voxels for a given contrast from a fMRI 3D volume) are in terms of estimated effects (continuous) and/or the presence/absence of a significant effect (binary). We distinguish individual and group estimates in **Figure 1** and describe the calculations below. For the continuous estimates of reliability described below, the analyses will be performed separately on task voxels that exceed and do not exceed an *a priori* specified threshold applied on the NeuroVault (Gorgolewski et al., 2015) meta-analysis collection that comprises the anticipatory win phase across 15 whole brain maps for the MID task (Wilson et al., 2018; Collection: 4258, Image ID: 68843). The *suprathreshold* task-positive voxels are those that exceed the threshold (*z* > 3.1) and the *subthreshold* task voxels are those that do not exceed the threshold (*z* < 3.1) in the map. We acknowledge that the threshold of *z* = 3.1 is arbitrary (uncorrected, *p*-value = .001) and that the voxels that fall below and above this threshold may not be significantly different (Gelman & Stern, 2006). However, to constrain the problem space this is a researcher’s decision that is made in these analyses (Gelman & Loken, 2014; Simmons et al., 2011).

**Figure 1.**
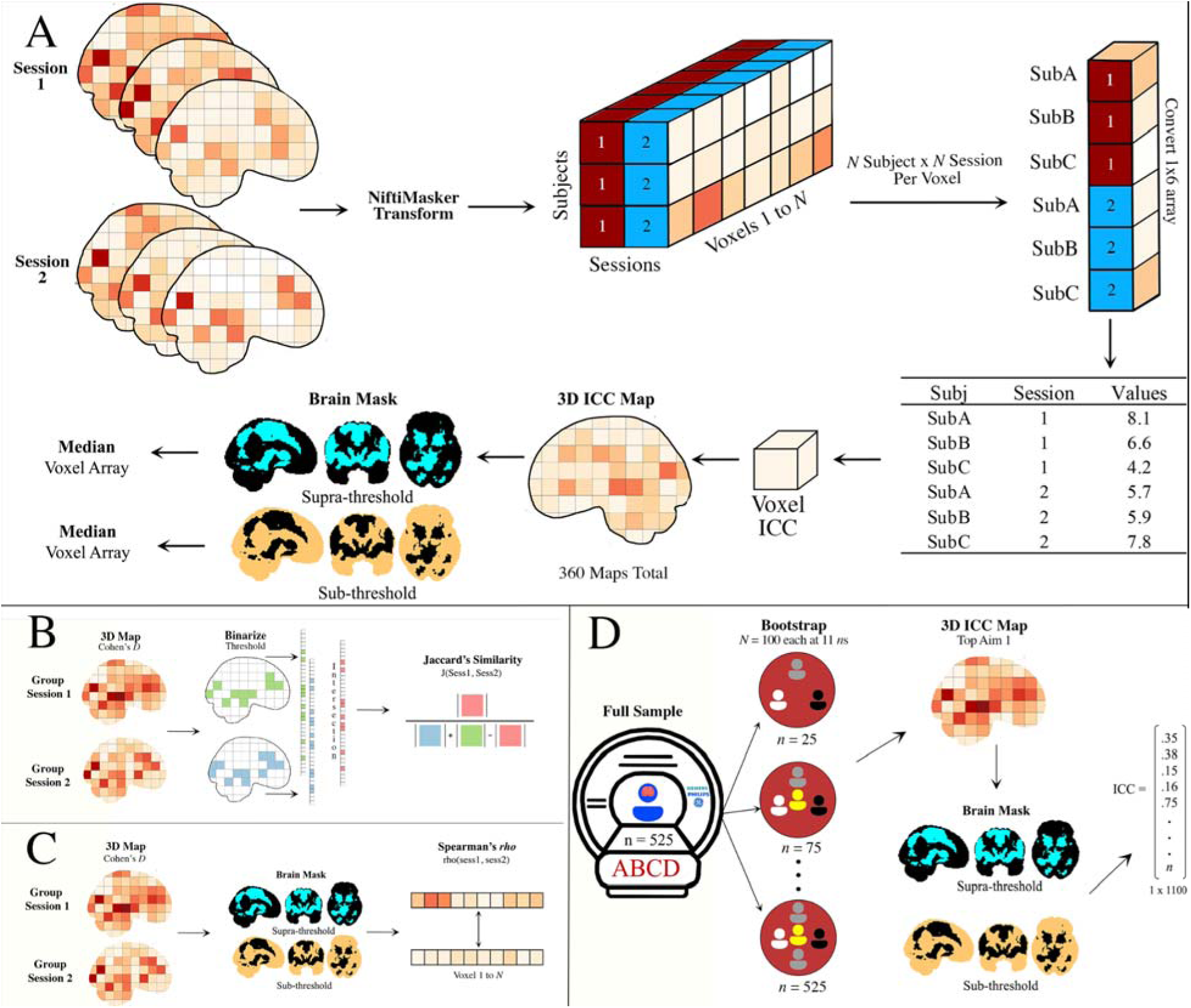
Diagram of (**A**) Continuous (individual), (**B**/**C**) binary/continuous (group) and (**D**) random subsampling of Estimates of Reliability across Measurement Occasions in 3D volumes of fMRI data. Group = group average of activation; Sub = Subject; ICC = Intraclass Correlation; Supra- and Sub-threshold mask is > 3.1 of NeuroVault Vault Image ID #68843 (Collection #4258)

#### Descriptive Statistics

The mean, standard deviation, count and frequencies are reported for demographic variables from the ABCD, AHRB and MLS datasets. For ABCD, AHRB and MLS, participants self-reported on Age, Sex and Race/Ethnicity. ABCD: Sex is reported as sex at birth (Male, Female, Other, or Not Reported); Race/Ethnicity is reported on a 5-item scale: White, Black, Hispanic, Asian, Other. AHRB: Sex is reported as sex at birth (Male or Female); Race/Ethnicity is available on a 4-item scale: White, Non-Hispanic, Black, Non-Hispanic, Hispanic/Latinx, Other. MLS: Sex is reported as Sex at Birth; Race is available on an 8-item scale: Caucasian, African American, Native American, Asian American, Filipino or Pacific Islander, Bi-Racial, Hispanic-Caucasian, and Other.

Behavioral data from the MID task, such as the mean and distribution of probe hit rate and mean response times (RT) across subjects, will be reported as supplemental information. The task design is programmed to achieve a probe hit rate of approximately 60% for each subject. It should be noted that the RT for the probe is not consistently collected across the ABCD, AHRB, and MLS datasets.

#### Impact of Analytic Decisions on Reliability in fMRI Data

First-, second- and group-level analyses are performed using Python 3.9.7 and Nilearn 0.9.2 (Abraham et al., 2014). Details about these three analytic steps are described below and the code is provided on Github (Demidenko, Mumford & Poldrack, 2024b). As listed in **Table 1** and described next, the analytic decisions will be limited to the first-level analysis.

##### Analytic Decisions

For reasons described in the introduction, the focus of analytic decisions in this paper will be on **four** categories: Smoothing, Motion Correction, Task Contrast and Task Parametrization. As reported in empirical studies and meta-analyses of task fMRI reliability (Bennett & Miller, 2010; Caceres et al., 2009), one way to improve reliability of fMRI data is by increasing the signal-to-noise ratio in the BOLD data through different smoothing kernels (Caceres et al., 2009), reducing motion effects in the fMRI data (Gorgolewski et al., 2013; Kennedy et al., 2022) and using task designs/contrasts that evoke increased neural activity (Han et al., 2022; Kennedy et al., 2022). These analytic decisions are described in greater detail in supplemental **Section 1.1**.

##### Within-run Analysis

A general linear model (GLM) is fit using Nilearn (e.g., *FirstLevelModel*) to estimate the response to task-relevant conditions in the BOLD timeseries for each participant/voxel. The BOLD timeseries are masked and spatially smoothed using specified full-width half-maximum (FWHM) Gaussian kernel options (see ‘Smoothing’ in **Table 1**) and the timeseries are prewhitened using an ‘ar1’ noise model. A GLM is fit (using *FirstLevelModel*) for a design matrix that includes the 15 task-relevant regressors (see task details in supplemental **Section 1.2**) and a set of nuisance regressors. Depending on the decision criteria (see ‘Motion Correction’ in **Table 1)**, nuisance regressors may include, for example, **A**) estimated translation and rotation (+ derivatives) of head motion or **A** + first eight aCompCor noise components and the corresponding cosine regressors for high pass filtering (with a cutoff of 128 seconds) that are calculated by fMRIPrep (see preprocessing of functional data). Task regressors are convolved with the SPM hemodynamic response function (HRF). The resulting beta estimates from the GLM, for each individual subject and run, are used to compute four contrasts for the MID task (see ‘Task Contrasts’ in **Table 1**).

##### Within-session Analysis

Per subject, each study collected two runs for each of two sessions. For each of the four contrast types, the beta and variances estimates from the two MID runs for each subject are averaged using Nilearn’s precision-weighted fixed effects model (i.e., *compute_fixed_effects*).

##### Group-level Analysis (within-session)

The MID task weighted fixed effects contrast files are used in a group-level mixed effect model (i.e., Nilearn’s *SecondLevelModel*) to average the within-subject estimates across subjects. These group maps are used as measures of the average activation patterns during the MID task in each of the studies across each of the four contrast types within each session.

The resulting individual and group maps from the four contrasts are used in calculating two different estimates of reliability (described in detail below). First, the resulting *within-run analysis* maps (i.e., for each run) are used for the continuous estimate of reliability *within* each session (i.e., reliability across runs). Then, the resulting *within-session analysis* maps, computed from the weighted fixed effects model, are used in the continuous estimate of reliability *between* the two sessions. Due to the temporal difference within and between sessions, the reliability within sessions would be hypothesized to be greater than between sessions. The resulting group-level analysis maps are used in the binary estimate of reliability *between* sessions.

#### Estimate of Reliability for Continuous Outcomes: Intraclass Correlation

Reliability for continuous outcomes at the individual level is estimated using ICC. The ICC is an estimate of between-subject and within-subject variance that summarizes how similar the signal intensities are for a given voxel from a 3D volume across sessions. As described in Liljequist et al. (2019), there are several versions of the ICC, which vary in whether the subjects and sessions are considered to be fixed (e.g., ICC[1]), subjects are considered to be random and sessions are considered to be fixed (e.g., consistency, estimated via ICC[3,1]) or the subjects and sessions are considered to be random (e.g., agreement, estimated via ICC[2,1]). In the case of these analyses, we assume that subjects are random but do not assume that sessions are random for two reasons. First, in the case of reliability of runs within a session, the runs are administered in a fixed manner and the state of the participant cannot be assumed to be random for each.

Second, in the case of reliability across sessions, during the follow-up session subjects have experienced the MRI environment and the task design in the scanner. In this case, again, it is difficult to assume that sessions are in fact random as the practice and session effects may be present. Thus, we estimate the consistency (ICC[3,1]) of the signal intensity for a given voxel across measurement occasions.

Several packages exist to calculate ICC and Jaccard/Dice coefficients. For example, *<u>ICC_rep_anova</u>* & *<u>Similarity</u>* in Python (Gorgolewski et al., 2011), *<u>fmreli</u>* in MATLAB (Fröhner et al., 2019) and *<u>3dICC</u>* in AFNI (Chen et al., 2017). However, these packages are either a) limited to a specific ICC calculation (e.g., ICC[3,1]), b) not easy to integrate into reproducible python code (e.g., *fmreli*), c) do not include similarity calculations (e.g., *3dICC*), or do not return information about between-subject, within-subject and between-measure variance components. Thus, to have the flexibility to estimate ICC(1), ICC(2,1) and ICC(3,1), Dice and Jaccard similarity coefficients and Spearman correlations simultaneously, we wrote and released an open-source Python package with reliability and similarity functions that works on 3D NifTi fMRI images.

The *PyReliMRI* v2.1.0 (Demidenko, Mumford & Poldrack, 2024a) Python package is used to calculate continuous estimates of reliability. *PyReliMRI* implements a voxel-wise ICC calculation (e.g., *voxelwise_icc*) for 3D NIfTI images between runs and/or between sessions (see the ICC example in study flowchart, Figure 1A). The function takes in a list of lists (e.g., list of session 1 and list of session 2) of ordered paths to the preprocessed data [in MNI space] for session 1 (or run 1) and session 2 (or run 2) subjects, and a binary [MNI space] brain mask. The package is flexible to take in more than 2 sessions (or runs). An ICC type option (e.g., ‘icc_1’, ‘icc_2’ or ‘icc_3’) indicates the type of ICC estimate that is calculated across the voxels within the masked 3D volume. The function returns a dictionary with five separate 3D volumes containing the voxel-wise (1) ICC estimate, (2) lower bound ICC, (3) upper bound ICC, (4) Between-subject variance (BS) and (5) Within-subject variance (WS) and, in case of ICC(2,1), (5) Between-measure variance, or the measurement additive bias. Like the ICC & 95% confidence calculation in the *pingouin* package (Vallat, 2018), the ICC confidence interval in *PyReliMRI* is calculated using the *f-*statistic (Bonett, 2002) to reduce the computation time compared to using bootstrapped estimates.

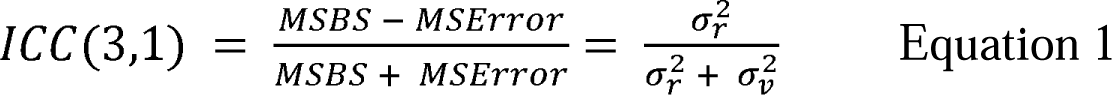

*Aim 1a*: evaluated the effect of analytic decisions (see **Table 1**; Figure 1A) on the ICC(3,1) (equation 1 for two measurement occasions) for individual [continuous] estimates of voxel activity across the ABCD, AHRB and MLS studies. The parameters in Equation 1 are: *MSBS* is the Mean Squared Between-subject Error and *MSError* is the Mean Squared Error. As described in Liljequist et al. (2019), the differences in the numerator is the between-subject variance (0^2^) and the denominator is the sum of the between-subject variance (0^2^) and the within-subject variance (or noise, [0^2^]). For each study, *voxelwise_icc* within the *brain_icc.py* script is used to estimate the voxel-wise ICC(3,1) for between run and between session reliability across the 360 model permutations. First, voxel-wise average and standard deviation from the resulting ICCs for the 360 model permutations are reported in two 3D volumes. Second, the range and distribution of median ICCs across each study (three) and analytic decision category (four) are plotted across suprathreshold task-positive and subthreshold ICCs using Rainclouds (Allen et al., 2019) and the median and standard deviation are reported in a table. Third, to visualize the ordered median ICCs across the 360 model permutations for suprathreshold task-positive and subthreshold ICCs, specification curve analyses are used (Simonsohn et al., 2020). Specifically, results across the 360 model permutations are reported using a specification curve to represent the range of estimated effects across the variable permutations. This consists of two panels: Panel A represents the *ordered* median ICC coefficients and the associated 95% confidence interval (across samples) colored based on no significance (gray), negative (red) or positive (blue) significance from the Null (Null here is 0) and Panel B represents the analytic decisions from each of the four categories (see **Table 1)** that produced the median ICC estimates. The median ICC estimates from the 360 models are reported separately for suprathreshold task-positive and subthreshold activation (the specification curve for all ICC estimates for suprathreshold task-positive and subthreshold activation are provided as supplemental information). Finally, to evaluate the effect of the analytic decisions on the median ICC, hierarchical linear modeling (HLM) is performed as implemented in the *lmer()* function from the *lme4* R package (Bates et al., 2020). HLM is used to regress the median ICC on the [four] analytic decisions as fixed effects with a random intercept model is fit (Matuschek et al., 2017)for samples across the suprathreshold task-positive and subthreshold maps. Multiple comparisons corrections are applied using the Tukey adjustment as implemented in the *emmeans* package (Lenth et al., 2023). For these HLM models, the interpretation focuses on the significant, non-zero effect of an independent variable (e.g., smoothing) on the dependent variable (e.g., median ICC) while the remaining independent variables are assumed to be zero.

*Aim 2:* evaluated the change in between- and within-subject variance across the analytic model permutations. Similar to Aim 1 (Figure 1A), *voxelwise_icc* within the *brain_icc.py* script is used to estimate the BS and WS across the 360 model permutations. The range and distribution of median BS and WS across each study and analytic decision category are plotted across suprathreshold task-positive and subthreshold BS/WS using Rainclouds. Then, two separate specification curve analyses report the *ordered* median BS and WS coefficients in one panel and the analytic decisions that produced the BS and WS estimates in a second panel separately for suprathreshold task-positive and subthreshold activation. Finally, like Aim 1, two HLMs are used to regress the median BS and median WS on the [four] analytic decisions as fixed effects with a random intercept only for sample across the suprathreshold task-positive and subthreshold maps. Multiple comparisons corrections are applied using the Tukey adjustment.

Like Aim 1, the interpretation focuses on the significant, non-zero effect of an independent variable (e.g., smoothing) on the dependent variable (e.g., median BS or median WS) while the remaining independent variables are assumed to be zero.

*Aim 3*: evaluated the sample size at which the ICC stabilizes (Figure 1D). The chosen pipeline is based on the highest median ICC across the studies for the suprathreshold task-positive mask from Aim 1a and is rerun for the ABCD sample. Based on this pipeline, the first-level analysis steps are repeated for N = 525 from the N = 2000 subsample for only the ABCD data. Then, *voxelwise_icc* within the *brain_icc.py* script is used to derive estimates of the median ICC, BS and WS for the between runs (e.g., measurement occasions) reliability across randomly sampled subjects for 25 to 525 subjects in intervals of 50. Similar to the methods in Liu et al. (2023), 100 iterations are performed at each N (with replacement) and the median ICC, the associated BS and WS estimates are retained from *voxelwise_icc*. The average and 95% confidence interval for the estimates across the 100 iterations is plotted for each interval of *N* with the y-axis representing the median ICC and x-axis representing *N*. The plotted values will be used to infer change and stability in the estimated median ICCs and variance components across the sample size. If stability is not achieved by *N* = 500, the sample is extended to *N* = 1,000 and the analyses are repeated.

#### Estimate of Reliability: Jaccard Coefficient for Binary & Spearman Correlation for Continuous Outcomes

The estimate of reliability for group analyses is estimated using the Jaccard Similarity for binary and Spearman correlation for continuous outcomes. The estimates are used to evaluate how the MID task evokes BOLD activation above a pre-specified threshold (*p* < .001) in the same voxels for *groups* of subjects across measurement occasions (run/session) in the ABCD, AHRB and MLS studies.

The *PyReliMRI* package is used. *PyReliMRI* calculates the similarity between two 3D volumes using a Jaccard’s coefficient which, in short, is the intersection divided by the union between two binary images (see Figure 1B) or the Spearman correlation, which is ranked correlation between two continuous variables (see Figure 1C). The Jaccard coefficient ranges from 0 to 1, whereby higher values reflect greater similarity between two images. Like the product-moment correlation, the Spearman correlation ranges from -1 to 1, whereby values >0 indicate a positive association between images and values <0 indicate a negative association between images. The function (i.e., *image_similarity*) takes in the paths for MNI *image file1* and *image file2*, a specified MNI mask and integer (i.e., z-stat/t-stat) at which to threshold the image. The images are masked (if a mask is provided), thresholded at the specified integer (if a threshold is provided) and the resulting images are binarized per user’s input (i.e., if threshold = 0, the resulting similarity = 1). Based on the specified similarity metric, the resulting estimates are similarity (e.g., Dice/Jaccard) or correlation coefficient (e.g., Spearman) between the two 3D NIfTI images. For similarity between 2+ NIfTI images, *pairwise_similarity* is used. Similar to *image_similarity*, *pairwise_similaity* takes in paths for an MNI mask, a threshold integer for the 3D volumes and the similarity type. Unlike *image_similarity*, *pairwise_similarity* allows for a list (2+) of paths pointing to 3D volumes and creates pairwise-combinations across the image paths between which to estimate similarity. The function returns the similarity coefficient in a dataframe with the resulting similarity (or correlation coefficient) and the image label (e.g., basename of the provided path for given volume).

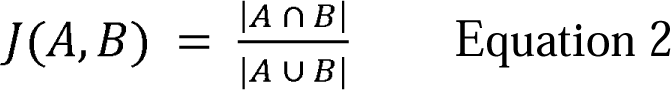

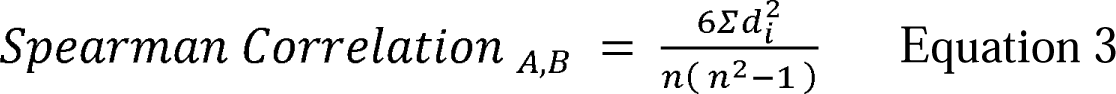

*Aim 1b*: evaluated the effect of analytic decisions (see Table 1) in the Jaccard’s similarity coefficient (Equation 2; Figure 1B) and Spearman correlation (Equation 3; Figure 1C) using the group binary & continuous estimates. In Equation 2, J(A, B) is the similarity coefficient between A (session 1) and B (session 2). This is derived from intersection, |A ∩ B|, which represents the elements that are common to both A and B divided by the union, |A ∪ B|, or the elements that are both in A and/or B. In Equation 3, the Spearman Rank Coefficient, as implemented in Scipy stats using *spearmanr* (Virtanen et al., 2020), is ranked correlation between unthresholded images A and B, whereby Σd² is the sum of squared differences between ranked values in session A and B, normalized by (n * (n² - 1)).

Since the Jaccard similarity coefficient is sensitive to thresholding and sample size (Bennett & Miller, 2010), in Aim 1b an equal sample size (e.g., N ∼ 60^3^) is chosen for each study to compare how the similarity between sessions varies across studies. For all 360 pipelines, a group-level (average) activation map is estimated for each session. In the case of the Jaccard coefficient, the group maps are thresholded at *p* < .001. In the case of the Spearman coefficient, the group maps are masked using a suprathreshold task-positive map from NeuroVault (https://identifiers.org/neurovault.collection:4258; Image ID: 68843). Then, the paths for the pipelines and sessions are called using the *pairwise_similarity* within the *similarity.py* script. The resulting coefficients report the similarity between analytic pipelines and sessions for each study. For each study, the coefficients are plotted to reflect the distribution and range of coefficients.

Both Jaccard’s and Spearman correlation are reported separately. Like Aim 1a & Aim 2, two HLMs are used to regress the Jaccard coefficients and Spearman correlation on the [four] analytic decisions nested within study. Multiple comparisons corrections are applied using the Tukey adjustment.

## Results

Given the breadth of the analyses (see **Table 2**), the results in the main text focus on the Session 1 between-run individual- and group-level reliability estimates for the supra-threshold mask. Differences are briefly noted for between-session reliability estimates and sub-threshold models and are reported in detail in the supplemental materials.

As permitted, aggregate and individual subjects’ data are made publicly available on NeuroVault (Gorgolewski et al., 2015) and/or OpenNeuro (Markiewicz et al., 2021). The complete set of group-level and ICC maps are publicly available on Neurovault for ABCD (6180 images; https://identifiers.org/neurovault.collection:17171), AHRB (2400 images; https://identifiers.org/neurovault.collection:16605) and MLS (2400 images; https://identifiers.org/neurovault.collection:16606). For each run and session, the BIDS input data and derivations for MRIQC v23.1.0 and fMRIPrep v23.1.4 are available on OpenNeuro for AHRB (Demidenko, Huntley, et al., 2024) and MLS (Demidenko, Klaus, et al., 2024). Since the ABCD data are governed by a strict data use agreement (March 2024), the processed data will be made publicly available via the NDA at a later date as part of the ABCC release. The final code for all analyses is publicly available on Github (Demidenko, Mumford & Poldrack, 2024b).

In the supplemental information of the Stage 1 submission, we stated that we would adjust the smoothing weight for the MLS as its voxel size, 4 mm anisotropic, would result in greater inherent smoothness of the data than ABCD/AHRB samples (2.4 mm isotropic voxel). A weight of .50 was applied to the smoothing kernels of the MLS data. This resulted in 3.6, 4.8, 6.0, 7.2 and 8.4 mm smoothing kernels for the AHRB/ABCD data and 3.0, 4.0, 5.0, 6.0 and 7.0mm smoothing kernels for the MLS data (**Figure S4**). In the results, the MLS ordinal values are relabeled to map onto the values used for AHRB/ABCD for reporting purposes.

*Note:* Estimates in supplemental **Table S5**

### Deviations from Stage 1 Registered Report

There are one moderate and two minor deviations from the Stage 1 Registered Report (https://doi.org/10.17605/OSF.IO/NQGEH). First, fieldmap-less distortion correction is not applied on the MLS data because the data were collected using spiral acquisition. The ABCC data selects a single fieldmap within a session to apply on *all* of the functional runs, so subjects without a fieldmap folder are excluded and fieldmap-less distortion correction is not used on the ABCD data. In AHRB, fieldmap-less distortion correction was used for only *one* subject.

Second, in Aim 1b we proposed to use thresholded images (e.g., *p* < . 001, approx. *t* > 3.2) to estimate the Jaccard/Spearman similarity between the model permutations for the estimated group maps. However, this statistic is arbitrarily sensitive to differences in the number of model permutations when subjects are excluded in cases of failed preprocessing features, such aCompCor mask errors. To improve the interpretability of the similarity estimates across analyses with different numbers of included observations (see supplemental **Figure S3**), we converted all *t*-statistic group maps to Cohen’s *d* effect size maps using the formula: 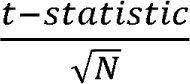.

Cohen’s *d* = .40 is used as the alternative threshold for Aim 1b as for pre-registered N ∼ 60 a conversion of *t-statistic* = 3.2 would be near this threshold. Third, the analyses proposed to evaluate 360 analytic decisions across the three samples. However, no subjects in the final AHRB and MLS samples exceeded mean FD = .9 so it was not possible to perform Motion option 5 (Motion option 3 + exclude mean FD ≥ .9) or Motion option 6 (Motion option 4 + exclude mean FD ≥ .9). As a result, the model permutations are restricted to 240 permutations (5 = FWHM, → 4 = Motion; 3 = Model Parameterization; 4 = Contrasts) with relevant data across the three samples and are the focus of the below analyses.

### Descriptive Statistics

The final sample for Aim 1 and Aim 2 for ABCD, AHRB and MLS samples (mean FD < .90) from the University of Michigan site that had two runs for at least two sessions, had behavioral data, and passed QC are *N*s 119, 60 and 81, respectively. For *N* = 15 subjects in the ABCD sample aCompCor ROIs failed, but otherwise the data passed QC and so these subjects were not excluded in Motion option3 and option4 models that include the top-8 aCompCor components as regressors. The final random subsample from the Baseline ABCD data for Aim 3 is *N* = 525.

Demographic information across the three samples for Aim 1 and Aim 2 (ABCD = 119; AHRB = 60; MLS = 81) are reported in supplemental **Table S4**. The average number of days between sessions is largest for the MLS sample (1090 days), followed by ABCD (747 days) and AHRB (419 days; **Figure S5**). On average, mean FD was higher in the ABCD sample versus the AHRB and MLS samples (**Figure S6**; **Table S5**). The samples also differed on average response probe accuracy (%), whereby on average MLS participants had a higher and faster probe response accuracy than ABCD and AHRB samples.

The estimated model efficiency, defined as 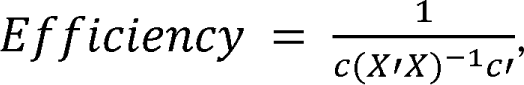, varied as a function of Model Parameterization and Contrast types across the three samples (see **Figure S7)**. The Anticipation Model (i.e., onset times locked to Cue onset and duration the combined duration of Cue and Fixation cross) was consistently estimated to be the most efficient model across the three samples for the *Large Gain* versus *Neutral* and *Small Gain* versus *Neutral* contrasts.

### Aim 1a: Effect of analytic decisions on median ICC estimates for individual continuous maps

Aim 1a proposed to evaluate the estimated individual map similarity between measurement occasions (runs/sessions) using the ICC(3,1) across 240 pipeline permutations. In **Table S5** (Figure 2), the median between-run Session 1 ICCs are slightly lower than the between-session ICCs (<u>between-run</u>: ABCD = .11 [range: -.04 - .43]; AHRB = .18 [range: .00 - .52]; MLS = .18 [range: .04 - .55]; <u>between-session</u>: ABCD = .15 [range: .03 - .34]; AHRB = .21 [range: .04 - .53]; MLS = .21 [range: .06 - .47]). The mean and standard deviation of the 3D volumes across the 240 analytic decisions are reported in supplemental **Figure S8**. Across the three samples, a consistent pattern is observed, whereby the regions with the highest ICCs, on average, are within the visual and motor regions. Notably, the lowest ICCs, on average, are within the ventricles and white matter. The supra-threshold distribution of the median estimates across the four model options and three samples are reported in Figure 3 and the specification curve of the median ICC estimates are reported in Figure 4. Note, the sub-threshold reported in supplemental **Figure S10**.

**Figure 2.**
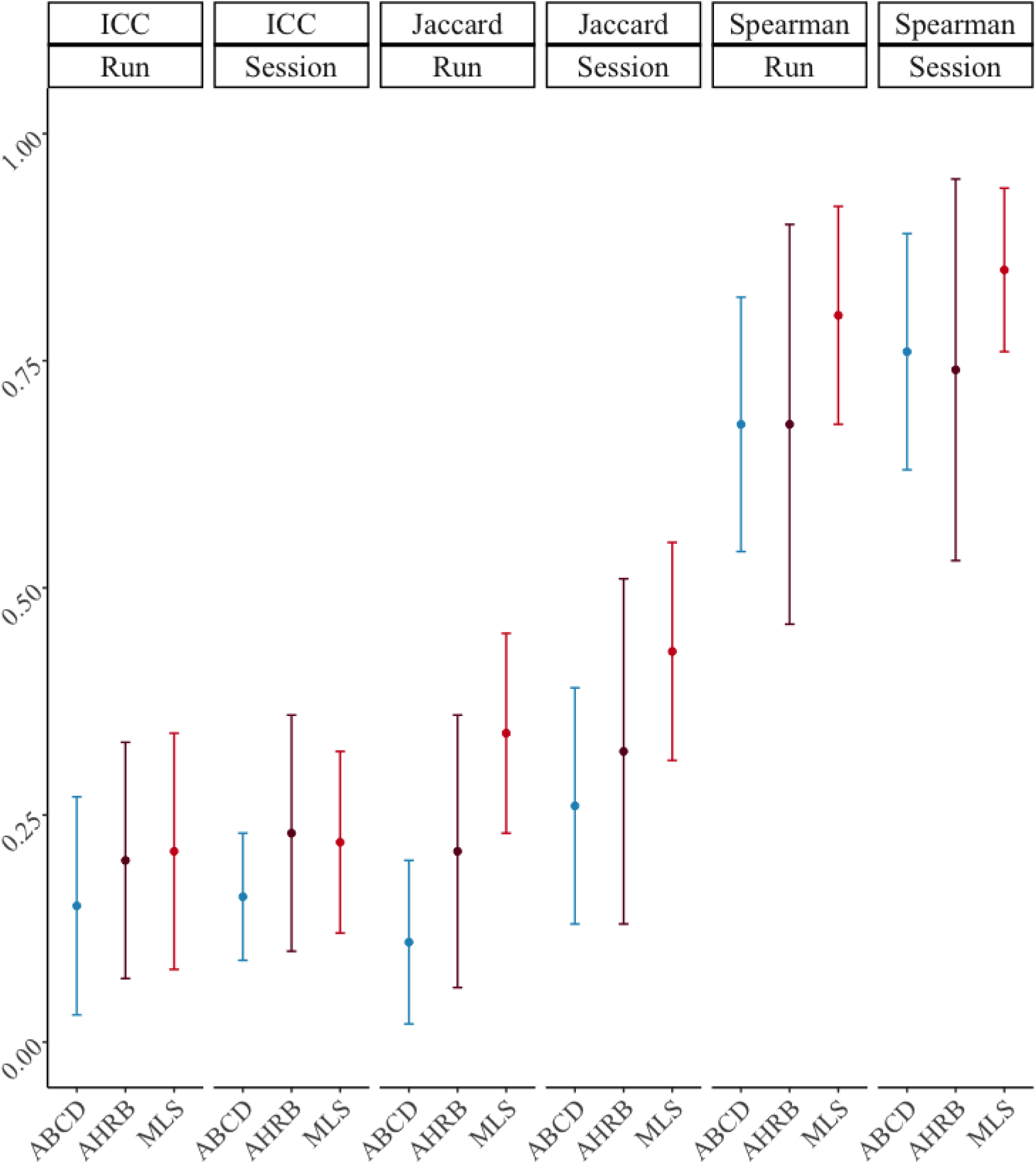
Session 1 Between-runs and Between-sessions: Mean +/- 1 Standard Deviation (SD) of Supra-threshold median Intraclass Correlation Coefficient (ICC), Jaccard and Spearman Similarity Coefficients from 240 analytic models across ABCD, AHRB and MLS Samples.

**Figure 3.**
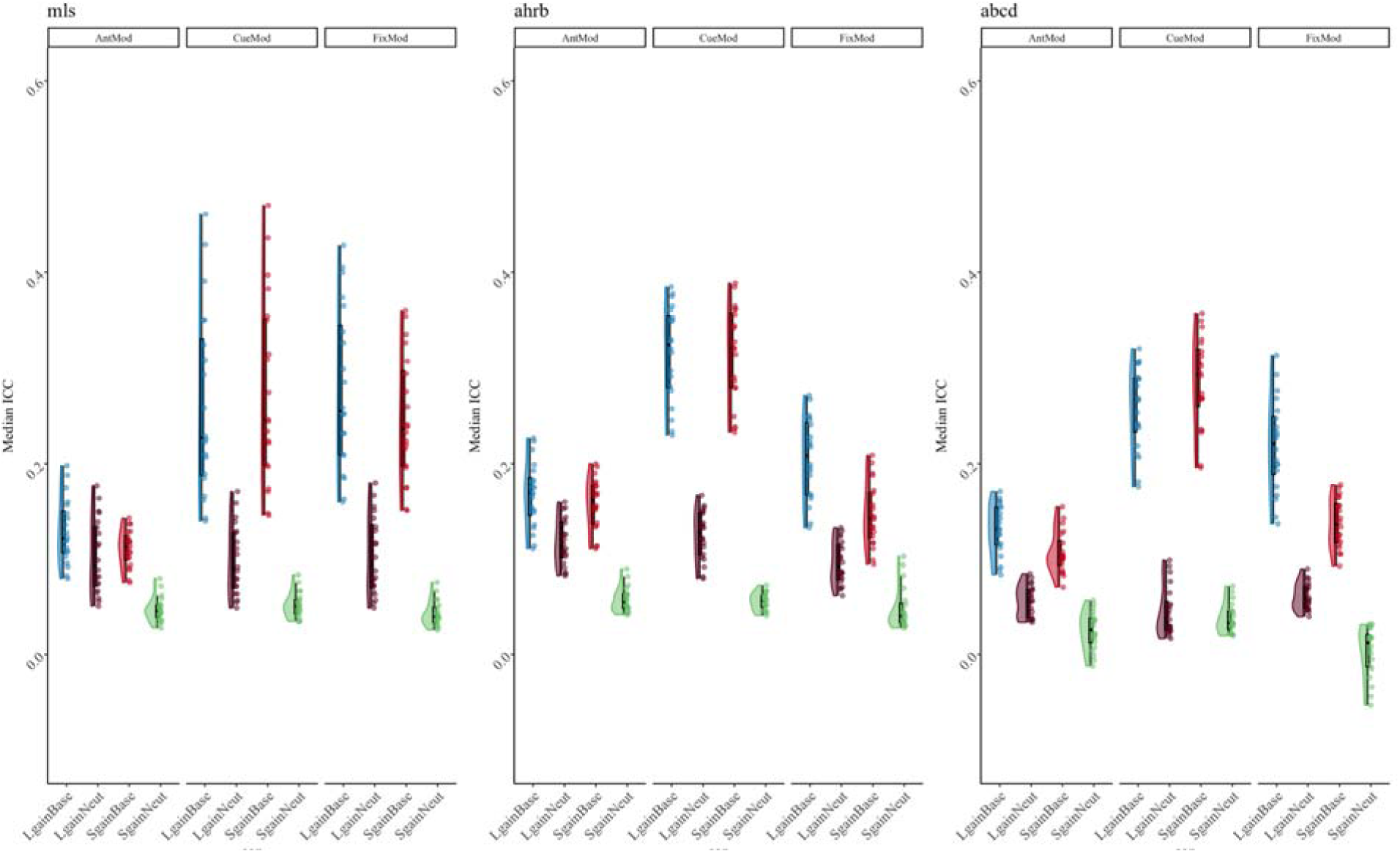
Supra-threshold Median ICC Session 1 between-run reliability estimates for Contrast (con) and Model Parameterization analytic options across the ABCD, AHRB and MLS samples. Complete distribution across four analytic options in supplemental **Figure S9**.

**Figure 4.**
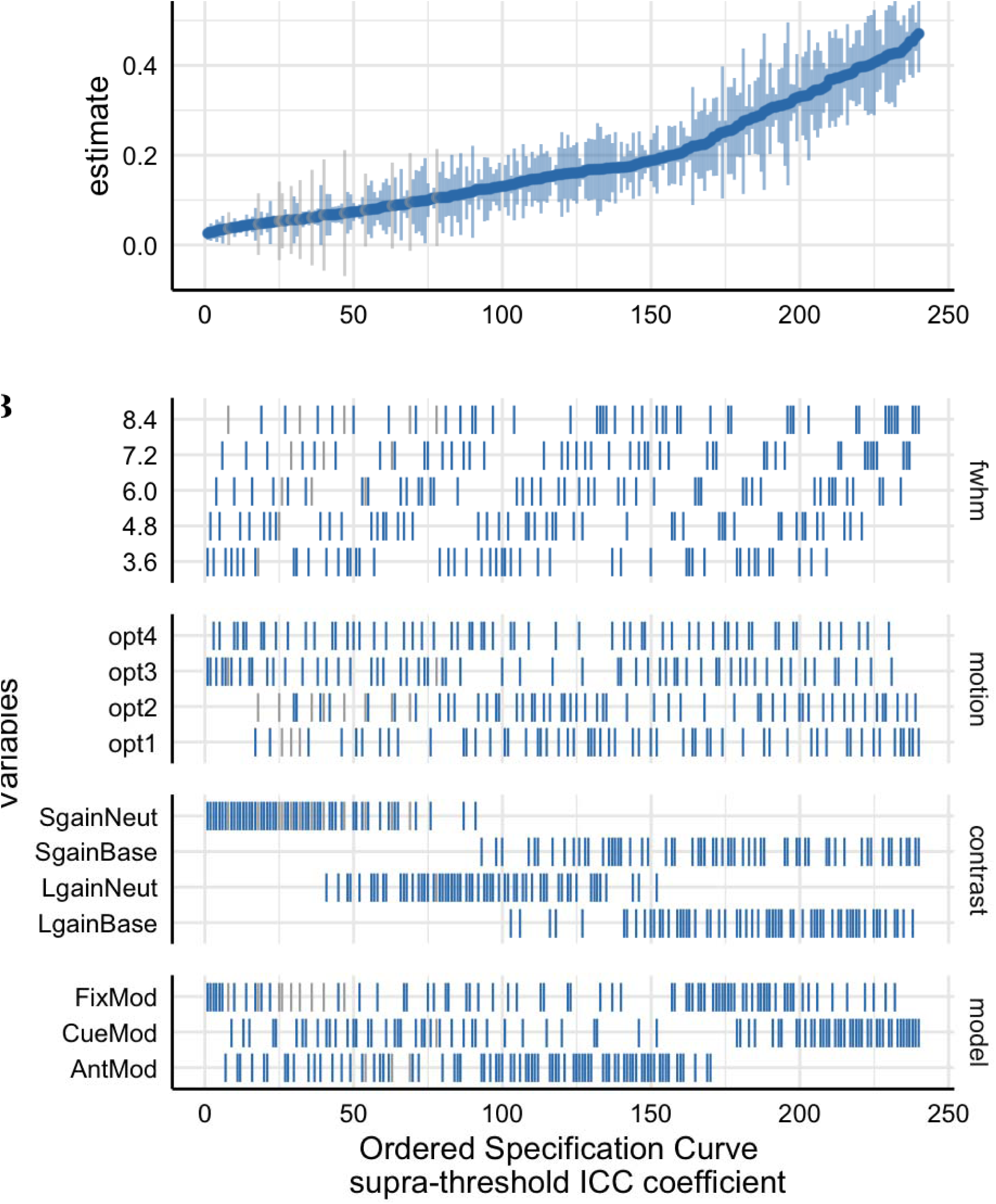
The supra-threshold Specification Curve of the Session 1 Between-run Median ICC estimates across 240 pipeline permutations for the ABCD, AHRB and MLS samples. Full length of estimates reported in **Figure S11.** A. The distribution of the point estimate (average) and distribution (error bars) across the three samples. B. The model options (four) associated with each estimate.

The effects reported in Figure 3 and Figure 4 illustrate that the largest differences in the median ICC estimate is associated with model parameterization and the contrast type. Even though the Anticipation Model (‘AntModel’) has the highest estimated contrast efficiency within each sample, contrary to our hypothesis the highest median ICC is associated with the Cue Model (‘CueMod’) in which the onset and duration are locked to the cue stimulus. However, using an interaction to probe the distributions in Figure 3, *post hoc* analyses suggest the Cue Model finding is largely driven by the *Implicit Baseline* contrasts (see Aim 1b) and the plot of the Model Parameterization-by-Contrast in supplemental **Figure S12** suggests negligible differences between Model Parameterization for the contrast of the *Neutral* contrasts.

Independent of model parameterization and consistent with our hypothesis and previous reports in the task fMRI literature (Han et al., 2022; Kennedy et al., 2022), the highest median ICC is consistently observed for the *Large Gain* versus *Implicit Baseline* contrast. In line with the reported estimates in Figure 3 and Figure 4, the HLM model for the supra-threshold mask shows a significant association between different FWHM, Motion, Model Parameterization and Contrasts model options compared to their respective reference values (**Table 3**). Specifically, the median ICC estimates increased with larger smoothing kernels and decreased with more stringent motion correction. Additionally, primarily driven by the *Implicit Baseline* conditions, median ICC for the ‘CueMod’ and ‘FixMod’ increased in comparison to the ‘AntMod’ (see interaction plot in **Figure S12**). Last, median ICC decreased in comparison to the *Large Gain* versus *Implicit Baseline* contrast. For example, the contrast *Large Gain* versus *Neutral* has an median ICC that is .17 lower, on average, compared to the *Implicit Baseline* contrast when holding other decisions constant (see marginal means comparisons in supplemental **Table S6)**.

**Table 3.**
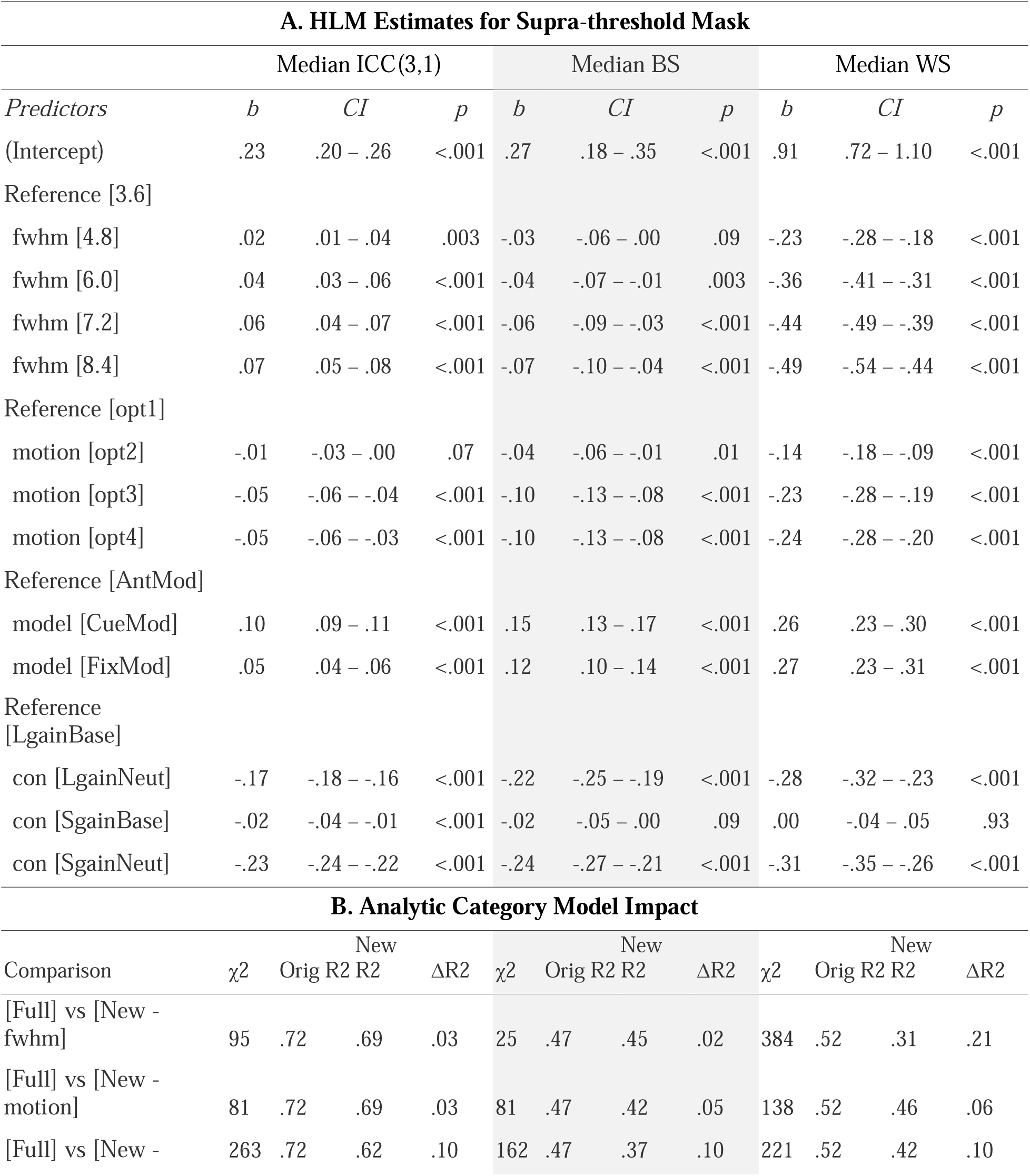
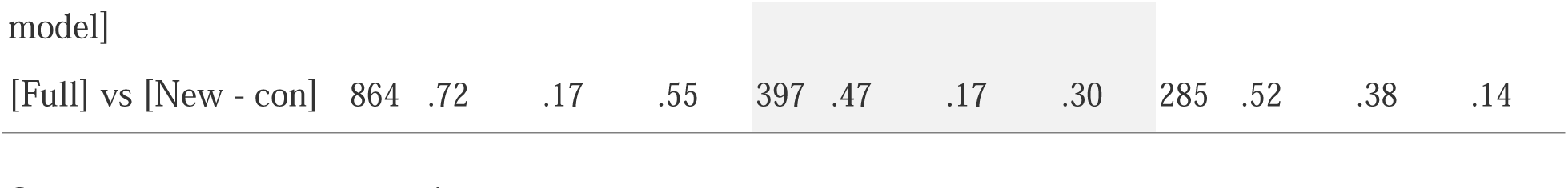
Hierarchical Linear Model: (A) Linear associations between the analytic decisions and the Session 1 between-run median Intraclass Correlation Coefficient (ICC[3,1]), Between-subject (BS) and Within-subject variance (WS) from supra-threshold mask and (B) the impact of the analytic category on the marginal R^2^.

While most parameters are significant in **Table 3**, the effects vary in their relative importance in the model. The variability in the median ICC estimate across 240 pipelines and three samples is best explained by contrast (marginal ΔR2: .55) and model parameterization (marginal ΔR2: .10). FWHM and motion had a smaller impact on ΔR2, .03 and .03 respectively. In fact, including aCompCor components (Motion option 3) and aCompCor components + censoring high motion volumes (Motion option 4) is associated with a slight decrease in the median ICC estimate as compared to no motion correction (Motion option 1), b = -.05 and b = -.05, respectively. A similar finding is observed for the sub-threshold mask, whereby the contrast (ΔR2: .56) and model parameterization (ΔR2: .10) decision had a larger impact on ΔR2 than the FWHM (ΔR2:.04) or motion (ΔR2: .02) decisions (see **Figure S14**; **Table S7).** In general, the voxelwise distribution of ICC estimates tends to be higher for the supra-threshold mask than the sub-threshold masks (see supplemental Figure S14). Interpretations are generally consistent for between-session median ICC estimates across the 240 pipeline permutations (see **Table S9** and **Figure S18**, **S19**).

We had hypothesized that the ICC estimates in the older samples (AHRB/MLS) would meaningfully differ from the younger sample (ABCD). Overall, ICC estimates were higher in the older than younger sample for *between-run*, *t*(497.2) = 5.53, p < .001, *d* = .43, and *between-session*, *t*(669.9) = 9.57, p < .001, *d* = .66.

### Summary of Findings for Aim 1a

Overall, between-run ICCs are slightly lower than between-session ICCs. Across the three samples, the highest ICCs, on average, are within visual and motor areas and the lowest ICCs are within the ventricles and white matter. In Table 1, it was hypothesized that the optimal analytic decisions would be: FWHM Smoothing 2.5x the voxel size, Motion correction that includes translation/rotation, their derivatives, the first 8 aCompCor components and exclusion of > .90 mFD subjects, the anticipation Model Parameterization, and Contrast *Large Gain* > *Implicit Baseline*. Contrary to registered hypotheses: (1) smoothing had a small but linear effect on ICC estimates, whereby the largest median ICC was for the largest FWHM smoothing kernel (3.5x voxel size); (2) Motion correction had minimal and negative impact on median ICCs in case of more rigorous corrections; and (3) the Cue and Fixation Models had higher estimated median ICCs than the Anticipation model. *Post hoc* analyses illustrated Model Parameterization is largely driven by the Implicit Baseline contrast, as Model Parameterization has a negligible impact on between condition contrasts. Consistent with registered hypotheses, the *Large Gain* versus *Implicit Baseline* had the highest estimated median ICC. Contrary to registered hypotheses, there was little evidence to suggest that analytic decisions differentially impacted estimated median ICCs between developmental samples (e.g., oldest MLS/AHRB versus younger ABCD data). Finally, the older samples (AHRB/MLS) had higher between- and between-session estimated ICCs than the younger sample (ABCD).

### Aim 1b: Effect of analytic decisions on Jaccard (binary) and Spearman (continuous) similarity estimates of group maps

Aim 1b proposed to evaluate the estimated group map similarity between measurement occasions (runs/sessions) using a Jaccard similarity for thresholded binary maps and a Spearman similarity for continuous measures across the 240 pipeline permutations. The distribution of the estimates across the four model options and three samples are reported in Figure 5 for Jaccard and supra-threshold Spearman similarity. The specification curve of the Session 1 between-run estimates are reported in Figure 6 for Spearman similarity (see **Figure S21** for Jaccard). Based on the group-level Cohen’s *d* maps, there is a high similarity between the *Small Gain* and *Large Gain* versus *Implicit Baseline* (and *Large Gain*) contrasts that appears to be driven by the *Implicit Baseline* condition and high similarity between Cue and Fixation models (see **Figure S22**).

**Figure 5.**
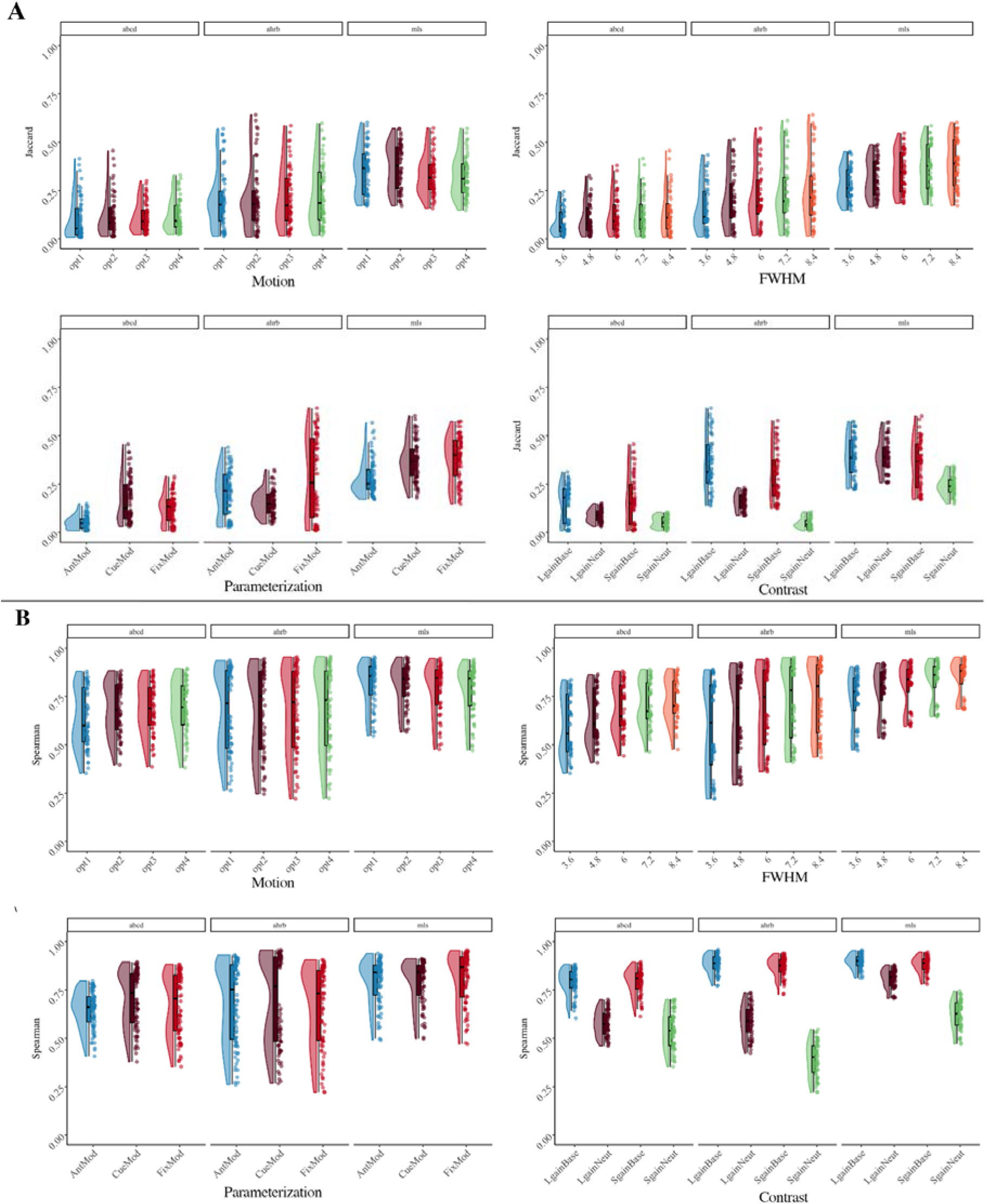
(**A**) *Jaccard* and (**B**) *supra-threshold Spearman Session 1 Between-run* similarity estimates across [Four] analytic options for between-run reliability across the ABCD, AHRB and MLS samples.

**Figure 6.**
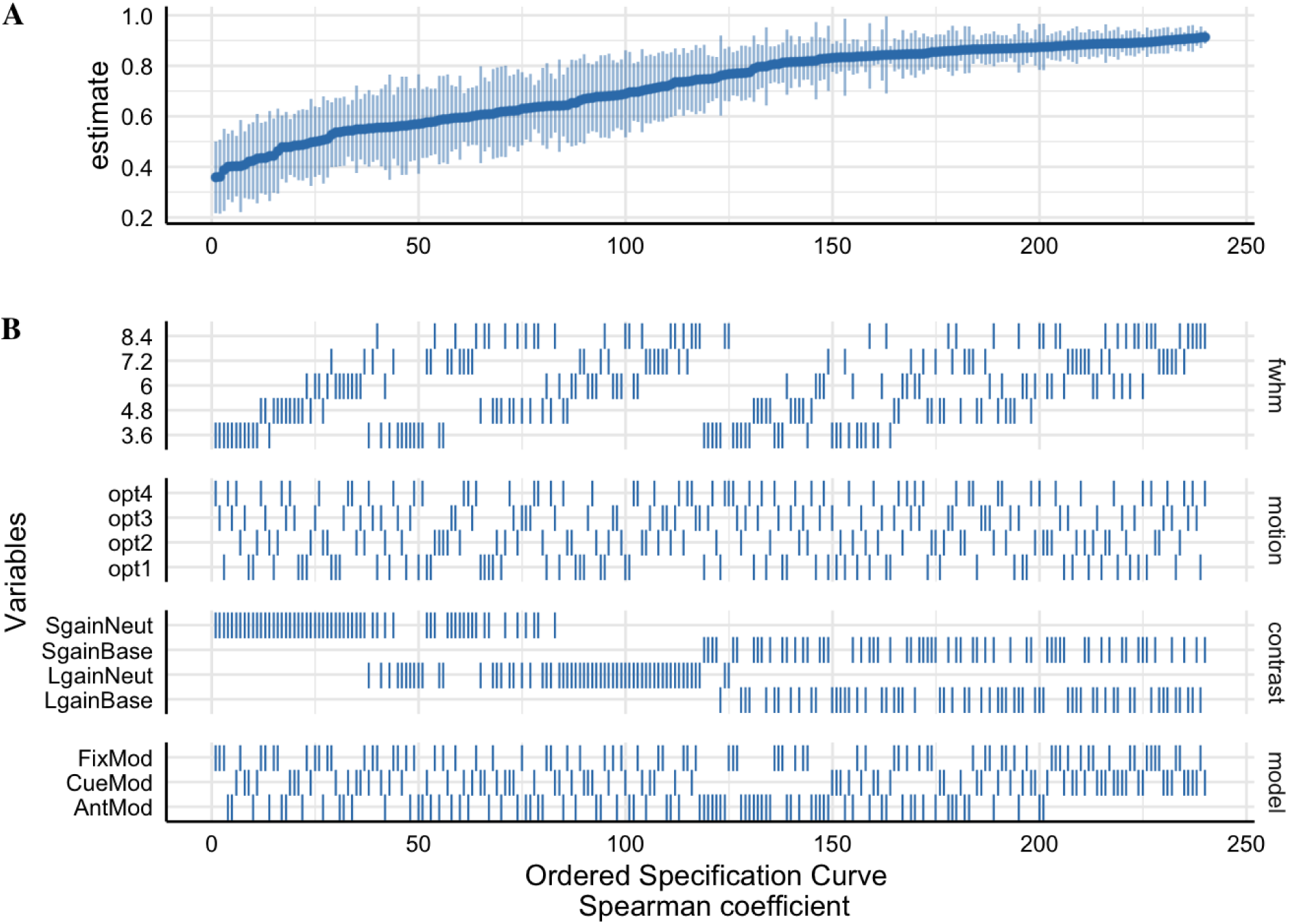
The supra-threshold Specification Curve of the *Session 1 Between-run Spearman similarity* estimates across 240 pipeline permutations for the ABCD, AHRB and MLS samples. A. The distribution of the point estimate (average) and distribution (error bars) across the three samples. B. The model options (four) associated with each estimate.

Similar to Aim 1a (**Table S5**; Figure 2), on average the Session 1 between-run supra-threshold Spearman similarity is slightly lower than the supra-threshold between-session Spearman similarity (between-run: ABCD = .68 [range: .35 - .89]; AHRB = .73 [range: .22 - .96]; MLS = .84 [range: .47 - .96]; between-session: ABCD = .80 [range: .40 - .94]; AHRB = .82 [range: .32 - .97]; MLS = .87 [range: .59 - .97]). A similar trend is observed for the Jaccard Similarity coefficient. The effects reported in Figure 5 illustrate that the analytic categories have unique impacts on the estimated Jaccard and supra-threshold Spearman coefficients. While the Jaccard coefficient varies most across contrast and model parameterization options (Figure 5A), the Spearman similarity varies most across FWHM and contrast type (Figure 5B). The specification curve for the Spearman similarity coefficients illustrate a near ceiling similarity for estimates at the upper tail of the estimates and little variability across the three samples (Figure 6). The HLM estimates indicate that a change from 3.6 to 8.4 FWHM results in a *b* = .08 increase in Jaccard similarity and a *b* = .13 increase in Spearman similarity. Furthermore, the change from the contrast *Large Gain* versus *Implicit Baseline* to *Large Gain* versus *Neutral* results in a *b* = -.09 decrease in Jaccard Similarity and a *b* = -.20 decrease in Spearman similarity. While most parameters are significant in **Table 4**, the effects vary in relative importance in the model. The variability in the estimated coefficients across 240 pipelines and three samples is best explained by Contrast (marginal ΔR^2^: .21) and model parameterization (marginal ΔR^2^: .05) for Jaccard similarity coefficient, and Contrast (marginal ΔR^2^: .66) and FWHM (marginal ΔR^2^: .08) for supra-threshold Spearman similarity coefficient. Surprisingly, the motion regressor options had a near-zero impact on the variability on both Jaccard and Spearman similarity coefficients. Similar to Aim 1a, *post hoc* analyses illustrate an interaction between Contrasts and Model Parameterization (**Figure S23**), whereby the largest driver of Model Parameterization differences in the Spearman *rho* similarity is as a function of the contrasts included the *Implicit Baseline*.

**Table 4.** Hierarchical Linear Model: (A) Linear associations between the analytic decisions and the *Jaccard and Spearman supra-threshold* mask Session 1 between-run similarity and (B) the impact of the analytic category on the marginal R^2^.

| A. HLM Group-map Estimates |  |  |  |  |  |  |
| --- | --- | --- | --- | --- | --- | --- |
| Predictors | Jaccard |  |  | Spearman |  |  |
| | $b$ | $CI$ | $p$ | $b$ | $CI$ | $p$ |
| (Intercept) | .20 | .09 – .31 | <.001 | .76 | .69 – .83 | <.001 |
| Reference [3.6] |  |  |  |  |  |  |
| fwhm [4.8] | .03 | .01 – .05 | .004 | .05 | .04 – .07 | <.001 |
| fwhm [6.0] | .05 | .03 – .07 | <.001 | .09 | .07 – .10 | <.001 |
| fwhm [7.2] | .07 | .05 – .09 | <.001 | .11 | .10 – .13 | <.001 |
| fwhm [8.4] | .08 | .06 – .10 | <.001 | .13 | .12 – .15 | <.001 |
| Reference [opt1] |  |  |  |  |  |  |
| motion [opt2] | .01 | -.00 – .03 | .13 | .01 | -.00 – .03 | .05 |
| motion [opt3] | .00 | -.02 – .02 | .85 | .01 | -.00 – .02 | .20 |
| motion [opt4] | .00 | -.01 – .02 | .69 | .01 | -.00 – .03 | .08 |
| Reference [AntMod] |  |  |  |  |  |  |
| model [CueMod] | .05 | .04 – .07 | <.001 | .02 | .01 – .03 | <.001 |
| model [FixMod] | .08 | .07 – .10 | <.001 | .01 | -.00 – .02 | .18 |
| Reference [LgainBase] |  |  |  |  |  |  |
| con [LgainNeut] | -.09 | -.10 – -.07 | <.001 | -.20 | -.21 – -.18 | <.001 |
| con [SgainBase] | -.03 | -.05 – -.01 | .001 | -.01 | -.02 – .00 | .17 |
| con [SgainNeut] | -.18 | -.20 – -.16 | <.001 | -.34 | -.35 – -.32 | <.001 |

B. Analytic Category Model Impact
| Comparison | $\chi^2$ | Orig R2 | New R2 | $\Delta R2$ | $\chi^2$ | Orig R2 | New R2 | $\Delta R2$ |
| --- | --- | --- | --- | --- | --- | --- | --- | --- |
| [Full] vs [New - fwhm] | 78 | .30 | .26 | .04 | 292 | .74 | .66 | .08 |
| [Full] vs [New - motion] | 3 | .30 | .30 | .00 | 5 | .74 | .74 | .00 |
| [Full] vs [New - model] | 104 | .30 | .25 | .05 | 14 | .74 | .73 | .01 |
| [Full] vs [New - con] | 348 | .30 | .09 | .21 | 1205 | .74 | .08 | .66 |

The group-level maps indicate a notable difference in contrasts using the *Neutral* and *Implicit Baseline* conditions (NeuroVault ABCD: https://identifiers.org/neurovault.collection:17171 AHRB: https://identifiers.org/neurovault.collection:16605; MLS: https://identifiers.org/neurovault.collection:16606). As **Figure S22** shows, the *Large Gain* versus *Neutral* contrast reflects a qualitatively comparable activation map across Cue, Fixation and Anticipation Models. On the other hand, the *Large Gain* versus *Implicit Baseline* contrast differs across models, where the most notable pattern is that the Cue model is negative of the Fixation model across the samples. Specifically, in ABCD, AHRB and MLS there is increased negative activity in the insular, visual, motor and visual areas, in the Cue Model, and this pattern is mostly opposite of the Fixation Model. Meanwhile, in the Anticipation model there is high positive activity in the dorsal striatal, SMA and Insular regions. This reflects the variable meanings of *Implicit Baseline* across the models. The relative symmetry between the Cue and Fixation models is consistent with the fact that each serves as the B_o_ in the models, e.g., 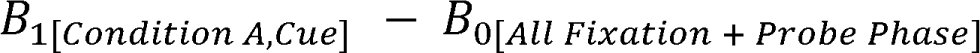 and 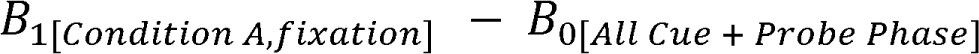. The Anticipation model is more variable as it is contrasted with a more narrow phase of the task, e.g., *B*_1[*condition A,Cue + Fixation*]_ – *B*_0[*Probe Phase*]_.

### Summary of Findings for Aim 1b

Similar to Aim 1a, on average, the supra-threshold Session 1 between-run Spearman and Jaccard similarity is slightly lower between-session similarity. Spearman similarity meaningfully differed across Contrast, Model Parametrization and Smoothing, and it is near the ceiling for the upper tail of the Spearman similarity estimates. Like Aim 1a, Model Parametrization is driven by the Implicit Baseline. Finally, mean-based group activity maps illustrate that the Cue and Fixation models are opposite of each other when the contrast is a between condition and implicit baseline comparison.

### Aim 2: Effect of analytic decisions on median BS/WS estimates from individual continuous maps

Aim 2 proposed to evaluate the changes in the Between-subject variance (BS) and Within-subject variance (WS) components that differentially relate to the ICC(3,1) across the 240 workflow permutations. The supra- and sub-threshold distributions across the four model options and three samples are reported in supplemental **Figure S24 & S25** and specification curves for BS in supplemental **Figure S28** and WS in supplemental **Figure S29**. The HLM estimates (**Table 3**) suggest that the Implicit Baseline contrasts increase BS variance and more stringent motion correction decrease BS variance, and Implicit Baseline contrasts and larger smoothing kernels reduce WS variance. The variability in the estimated BS coefficients across 240 pipelines and three samples is best explained by Contrast (ΔR^2^: .30), model parameterization (ΔR^2^: .10) and then motion (ΔR^2^: .04). The variability in the estimated WS coefficients across 240 pipelines and three samples is best explained by FWHM (ΔR^2^: .21), Contrast (ΔR^2^: .14) and then model parameterization (ΔR^2^: .10). A comparable trend is observed in the between-session estimates (**Table S9**), with the exception of Contrast selection explaining more variability (ΔR^2^: .26) than FWHM (ΔR^2^: .16). We avoid interpreting the sub-threshold mask as it includes regions that are high-noise (e.g., white matter and ventricles) and drop-out areas (e.g. cerebellar and medial orbital frontal cortex) which exaggerates the BS and WS components.

### Aim 3: Stability of the ICC, BS and WS Components across Sample Size

As expected, based on sampling theory which demonstrates that variability decreases as a function of the square root of *N*, the variability in estimates decreased as *N* increased.

Specifically, the bootstrapped estimates for the median ICC, BS and WS change slowly at higher intervals of *N* (Figure 7). In *post hoc* comparisons of whole brain voxelwise ICC maps, the largest variability occurs below N = 275. As reported in supplemental **Figure S36**, at *N* = 25 the minimum and maximum median whole brain ICC maps have a wider voxelwise distribution of ICC values which are notably different (Cohen’s *d* = 1.9). With increasing *N,* Cohen’s *d* of the whole brain voxelwise distributions between the minimum and maximum 3D ICC maps narrows, *d* = 1.4 at *N* = 225 and *d* = 1.0 at *N* = 525, respectively.

**Figure 7.**
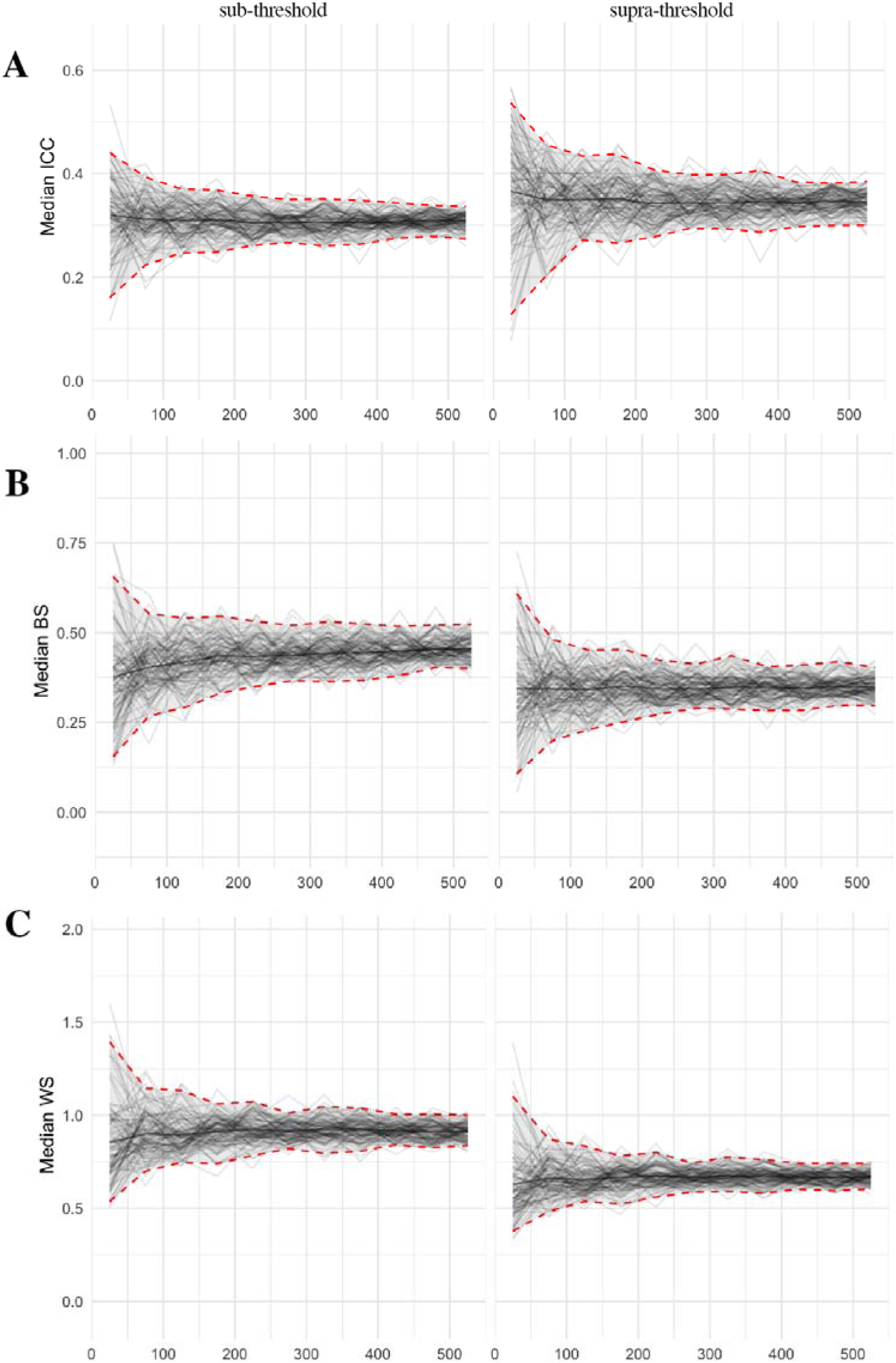
Changes in the Supra-& Supra-threshold Median Intraclass Correlation (ICC), Between-subject variance (BS) and Within-subject variance (WS) estimate in the ABCD sample for *N* 25 to 525 with 100 bootstraps at each *N* Note: Based on the top model from Figure 2: *Small Gain* vs *Implicit Baseline* Contrast, ‘CueMod’ Model, Motion option 1 and FWHM 8.4.

### Post Hoc Analyses

An exploratory set of analyses were performed to evaluate 1) the effect of analytic decisions on ICC for the Left and Right Nucleus Accumbens and 2) the association between voxelwise Cohen’s *d* estimates at the group-level and the voxelwise ICC maps. These are reported in supplemental **section 2.6**.

## Discussion

Understanding the analytic decisions that may consistently increase individual- and/or group-level reliability estimates has implications for the study of individual differences using fMRI. The current study expands on previous work by simultaneously evaluating the effects of smoothing, motion correction, task parameterization and contrast selection on the continuous and binary reliability estimates of BOLD activity during the MID task for run- and session-level data across three independent samples. The five major findings are: (1) The ICC(3,1) test-retest reliability estimates in the MID task are consistently low; (2) Group-level estimates of reliability are higher than individual [ICC] estimates; (3) Contrast selection and Model Parameterization have the largest impact on median ICC estimates, and Smoothing and Contrast selection has the largest impact on similarity estimates; however, gains in reliability across different contrasts comes at the cost of interpretability and may differ; (4) Motion correction strategies in these analyses did not meaningfully improve individual or group similarity estimates and, in some cases, *reduced* estimates of reliability; and (5) the median ICC estimate varied across sample size but the variability decreased with increased sample size. Excluding some differences, the results are relatively consistent across the three samples, runs and sessions, providing a comprehensive overview of how analytic decisions at the GLM impact reliability of estimated BOLD in commonly used versions of the MID task.

The findings from these multiverse analyses confirm previous reports that ICC estimates are relatively low in univariate task-fMRI and in the current state are inadequate measures for use in individual differences research (Elliott et al., 2020; Kennedy et al., 2022). Consistent with Elliott et al (2020), reliability estimates in the sub-threshold (or non-target mask) are lower than the supra-threshold of the MID task (target mask). The range of median ICCs varied across analytic decisions. Using commonly employed cut-offs (Cicchetti & Sparrow, 1981; Elliott et al., 2020; Noble et al., 2019), ICC estimates for *Large Gain* versus *Neutral* contrast are in the ‘Poor’ range and the *Large Gain* versus *Implicit Baseline* contrast ranged between ‘Poor’ and ‘Fair’ across the three samples. Test-retest reliability for the *Large Gain* (*Small Gain*) versus *Implicit Baseline* contrast are modulated by Model Parameterization, whereby the Cue Model had a meaningfully higher reliability than the Anticipation Model. However, this may come at the cost of validity, which is discussed below. Nevertheless, based on voxelwise distributions from the top performing model (Model: Cue Model, Contrast: *Small Gain* versus *Implicit Baseline*, Motion Correction: None, Smoothing: 8.4 mm kernel), visual and motor regions had the highest ICCs, in the ‘Fair’ to ‘Good’ range. *Post hoc* analyses of the bilateral NAc illustrate that, on average, ICC estimates in this region of interest are in the ‘Poor’ range. Notably, ICCs in this *post hoc* region were not meaningfully impacted by Model Parameterization but were impacted by Contrast and Motion correction, suggesting that test-retest reliability may be uniquely impacted by analytic strategy depending on the voxels under consideration. These findings illustrate that the test-retest reliability of the MID task is relatively low, even in the most common ROI such as the Left and Right NAc. While Kennedy et al. (2022, p. 13) speculated that low reliabilities in the ABCD sample may be attributed to the participants’ young age, our results demonstrate that median ICC estimates are *higher* in older than younger samples but reliability estimates in the MID task remain consistently low across early adolescents and late adolescents/young adults. To understand how analytic strategies differentially impact ICCs in different brain regions, we encourage future researchers to use the publicly available estimated maps to probe this question further.

Consistent with Fröhner et al. (2019), the group-level maps are not always representative of the individual-level maps across analytic decisions. On average, the Spearman *rho*, Jaccard coefficients and median ICC estimates are higher for the between-session than between-run estimates. Consistently, Spearman *rho* estimates are meaningfully higher for supra-threshold group maps than supra-threshold median ICC estimates derived from individual maps. This suggests that across each of the three samples, the MID task is relatively effective at eliciting a group-level activation map; however, the individual estimates are lower and more variable. In the context of the MID task, the between-run and between-session effects may be the result of within-session effects *decreasing* across runs (Demidenko, Mumford, et al., 2024). Notably, the higher between-session than between-run reliabilities is inconsistent with values reported in previous work (Fröhner et al., 2019), this is likely the result of those between-run estimates being based on randomly split-half (within runs) which are inflated as a result of dependencies in the model estimates within runs (Mumford et al., 2014). Nevertheless, the results here emphasize that group-level maps and group similarity are not a good indicator of individual-level reliabilities. This is unsurprising, considering that the MID task design was optimized to elicit activity in anatomical regions at a group-level and for averaged time-courses within an anatomical region (Knutson et al., 2003).

A major question of these analyses was: Are there decisions that *consistently* result in higher individual- (continuous) and/or group-level reliability estimates (continuous/binary)? The results across the analytic choices illustrate that reliability estimates are impacted most by contrast, model parameterization and smoothing decisions. Across the three samples, for between-run and between-session estimates, the contrast type had the largest influence of individual and group reliability estimates. Consistent with previous reports (Baranger et al., 2021; Han et al., 2022; Kennedy et al., 2022; Vetter et al., 2015, 2017), the contrast *Large Gain* (and *Small Gain*) versus *Implicit Baseline* had meaningfully higher estimated ICC, Jaccard and Spearman *rho* similarity estimates than the *Large Gain* versus *Neutral* contrast. The estimated ICC and Spearman *rho* coefficients for contrasts are modulated by the model parameterization, whereby the conditions including the *Implicit Baseline* are highest for the Cue Model parameterization. Conversely, ICC and similarity estimates are relatively stable across the three model parameterizations when comparisons are against the *Neutral* condition. Whether using contrasts or percent signal changes, estimates of BOLD activity suffer from decreases in reliability due to difference scores (Hedge et al., 2018). Where gains are observed from the less reliable *Large Gain* versus *Neutral* to the more reliable *Large Gain* versus *Implicit Baseline* contrast, it comes at the cost of interpretability and face validity that is expected in the estimated BOLD activity. Finally, higher FWHM smoothing kernels positively impacted between-run and between-session median ICC estimates and Spearman *rho* similarity estimates whereas motion correction strategies had a smaller but negative impact on these estimates (i.e., more stringent motion correction reduced reliability estimates). Decisions to smooth in the MID task are especially important given that larger smoothing kernels have been reported to spatially bias reward-related activity in the MID task (Sacchet & Knutson, 2013). In general, variability in reliability estimates decreased with large sample sizes.

Improvements in estimated reliability as a function of contrast selection may come at the cost of interpretability. For example, in the context of the *Large Gain* versus *Neutral* contrast, despite differences in the estimated efficiencies the ICC estimates are relatively stable across the model parameterizations in each of the three samples and the activation patterns are interpretable at the group-level. In the context of the *Large Gain* versus *Implicit Baseline* contrast, there are meaningful differences in the ICC estimates across model parameterizations, whereby the Cue and Fixation models demonstrate a substantial improvement over the Anticipation model parameterization, but the group-level activity patterns are less interpretable. As a researcher looking for BOLD estimates that are consistent from run-to-run or session-to-session for individual participants, the *Implicit Baseline* suggests a considerable and valuable improvement on the reliability of estimated values. However, the difference of means for the *Implicit Baseline* is complicated by the intercept in the GLM at the first level. For example, in the Cue Model parameterization, the intercept takes on the average for the unmodeled phase of the task which includes the fixation cross (between cue and probe phase) and the probe response phase. In this instance, isolating the difference of [Cue *Large Gain*] - [Fixation + Probe phase] to a specific cognitive function becomes especially challenging (Poldrack & Yarkoni, 2016; Price & Friston, 1997). It is well recognized that different definitions of “baseline”, whether rest, passive or task-related, in task-fMRI will result in different activation patterns (Newman et al., 2001). The use of “neutral” or “fixation” is a cause for caution as it impacts interpretability in various fMRI task designs (Balodis & Potenza, 2015; Filkowski & Haas, 2017). Here, we illustrated how contrasts with the unmodeled phases of a task (*Implicit Baseline*) may improve reliability estimates but may be heavily biased by the activity patterns throughout the task and diminish the validity of the measure. It is reasonable to suspect that subtle modeling deviations between similar and different task designs would further complicate comparisons between studies when using an *Implicit Baseline* condition.

In the context of test-retest reliability of estimated BOLD activity, it is important to consider alternative methods to improve reliability, estimation procedures and considerations of what a ‘reliable’ BOLD estimate implies. In general, the evidence here illustrates that the test-retest reliability for the modified version of the MID task is consistently low using the intraclass correlation (ICC[3,1]), even at its maximum. The analytic decisions at the GLM modeling phase demonstrated improvements in reliability from between-run to between-session. Higher between-session reliability may be related to decreasing activity from early to later runs (Demidenko, Mumford, et al., 2024) or based on the sessions being an average of two runs/increased trials (Han et al., 2022; Ooi et al., 2024). In the current analyses, we focused on univariate maps and the parametric, voxelwise ICC estimation procedures (ICC[3,1]). Parametric and non-parametric multivariate methods are reported to improve reliability estimates over univariate estimates using multi-dimensional BOLD data (Gell et al., 2023; Noble et al., 2021). For example, I2C2 is a parametric method that pools variance across images to estimate a global estimate of reliability using a comparable ratio as ICC (Shou et al., 2013) and the discriminability statistic is a non-parametric statistic that is a global index of reliability testing whether the between-subject distance between voxels is greater than the within-subject voxels (Bridgeford et al., 2021). Each of these metrics uniquely summarizes the within- and between-subject variability of the estimated BOLD data and so a consensus and definition of reliability in task-fMRI remains a challenge (Bennett & Miller, 2010). In our analyses we used the ICC as it estimated the reliability for each voxel in an easy-to-interpret coefficient that is useful in common brain-behavior studies. Cut-offs from the self-report literature (Cicchetti & Sparrow, 1981) are often leveraged in fMRI research (Elliott et al., 2020; Noble et al., 2019); however, these cut-offs should depend on the optimal level of precision necessary for the question and reasonable for the methods (Bennett & Miller, 2010; Lance et al., 2006). Some recommendations have been made to use bias-corrections in developmental samples to adjust for suboptimal levels of reliability (Herting et al., 2017), but these corrections should be used cautiously as they do not account for the underlying problems of the measure or the complexities in the data that prevent accurate measurement of the latent process (Nunnally, 1978).

## Study Considerations

The analytic decisions in the current analyses focused primarily on a subset of decisions at the First Level GLM model and its impact on estimates and supra/sub-threshold masks. As a result, other decisions were not considered that may arise at the preprocessing (Li et al., 2021), assumed hemodynamic response function (Kao et al., 2013; Lindquist et al., 2009), cardiac and respiratory correction (Allen et al., 2022; Birn et al., 2006), and the effects of different methods of signal distortion correction (Montez et al., 2023). Furthermore, we focused on voxelwise estimates of reliability which are typically noisier than *a priori* anatomical regions. It is unclear how much interpretation would change if ICC estimates were compared across variable parcellations. Nevertheless, we shared all aggregate maps for the three samples and the preprocessed data for the MLS/AHRB samples to facilitate reanalysis.

The results provide a comprehensive overview of individual and group reliability estimates for the modified version of the MID task, but it is challenging to infer how reflective these results are of alternate MID designs and different reward tasks. Based on prior reports of low test-retest reliabilities in task fMR, if a sufficient sample size is used, we suspect that results may be comparable to other MID and reward task designs. Future research should consider how reliability estimates change as a function of modeling decisions in different task paradigms.

## Conclusion

With the increasing interest in test-retest reliability in task fMRI and methods for improving reliability estimates of BOLD, the current study evaluated which decisions at the GLM model improved group and individual reliability estimates of reliability. In general, the findings illustrate that the MID task group activation maps are more reliable than individual maps across testing occasions and independent samples. Across group and individual models, between-session estimates are consistently higher than between-run estimates of reliability.

Furthermore, estimates of reliability were more variable at the median fMRI sample size and stabilized with *N*. While individual estimates of reliability are low (ICC[3,1]), contrasts and model parameterization meaningfully improved test-retest reliability. However, the improvement in reliability came at the cost of interpretability and may be region specific in the current version of the MID task. This underscores the importance of evaluating reliability in larger samples sizes and ensuring improved estimates reflect the neural processes of interest. While Model Parameterization and Contrast selection had the largest impact on voxelwise ICCs, further work is needed to expand on these findings by evaluating alternative brain regions and analytic decisions that may result in improved test-retest reliability that may be meaningful in individual differences research.

## Data & Code Availability Statement

*Adolescent Brain Cognitive Development* (ABCD) data: The ABCD BIDS data, MRIQC v23.1.0 and fMRIPrep v23.1.4 derivatives can be accessed through the ABCD-BIDS Community Collection (ABCC) with an established Data Use Agreement (see https://abcdstudy.org/). The data used in these analyses will be available at a future release onto the National Institute of Mental Health Data Archive. The complete set of group-level and ICC maps are publicly available on Neurovault for ABCD (6180 images; https://identifiers.org/neurovault.collection:17171).

*Michigan Longitudinal Study* (MLS) and *Adolescent Health Risk Behavior* (AHRB) data: The BIDS inputs, fMRIPrep v23.1.4 and MRIQC v23.1.0 derivates are available on OpenNeuro.org (MLS: https://doi.org/10.18112/openneuro.ds005027.v1.0.1 AHRB: https://doi.org/10.18112/openneuro.ds005012.v1.0.1). The complete set of group-level and ICC maps are publicly available on Neurovault for MLS (2400 images; https://identifiers.org/neurovault.collection:16606) and AHRB (2400 images; https://identifiers.org/neurovault.collection:16605)

*R and Python code*: The *.html* and *.rmd* file containing the code to be run on extracted estimates from reliability maps are available on Github with the associated output files containing the estimates across the models and samples. Likewise, all of the code for first level, fixed effect, group and ICC models are available online at https://github.com/demidenm/Multiverse_Reliability and DOI: 10.5281/zenodo.12701228x.

## Acknowledgements

MID is funded by the Ruth L. Kirschstein Postdoctoral Individual National Research Service Award through the National Institute on Drug Abuse (F32 DA055334-01A1). RAP is supported by the National Institute of Mental Health (R01MH117772 and R01MH130898). Thanks to Dr. Daniel Keating for agreeing to share the Adolescent Health Risk Behavior (AHRB; R01HD075806) study data and to Dr. Mary Heitzeg for agreeing to share the Michigan Longitudinal Study (MLS; R01HD075806) data for this project. The authors would also like to thank the research participants and staff involved in data collection of the Adolescent Brain Cognitive Development (ABCD) Study data. The ABCD Study is a multisite, longitudinal study designed to recruit more than 10,000 children ages 9 and 10 and follow them over 10 years into early adulthood. The ABCD Study is supported by the National Institutes of Health (NIH) and additional federal partners under award numbers U01DA041048, U01DA050989, U01DA051016, U01DA041022, U01DA051018, U01DA051037, U01DA050987, U01DA041174, U01DA041106, U01DA041117, U01DA041028, U01DA041134, U01DA050988, U01DA051039, U01DA041156, U01DA041025, U01DA041120, U01DA051038, U01DA041148, U01DA041093, U01DA041089, U24DA041123, and U24DA041147. The list of supporters is available at https://abcdstudy.org/federal-partners.html. The list of participating sites and study investigators is available at https://abcdstudy.org/study-sites/.

Thanks to members of the Cognitive Development and Neuroimaging Lab (CDNI) at the University of Minnesota, specifically Eric Feczko, rae McCollum and Audrey Houghton, for assisting and providing access to the ABCD-BIDS Community Collection (ABCC) data. The analyses here are based on data available from CDNI as of February 2024. The ABCC data repository grows and changes over time (https://collection3165.readthedocs.io/). Thanks to Krisanne Litinas at the University of Michigan for providing expert advice and scripts to convert the AHRB data into BIDS format. Thanks to Mary Soules and Ryan Klaus (with assistance from Krisanne Litinas) at the University of Michigan for working to convert the MLS data to BIDS format.

## Author’s Contribution

MID obtained data sharing agreements. MID conceptualized the study with critical input from RAP. MID defined the methodology with critical input from RAP and JAM. MID curated the analytic code and performed the formal analysis and interpretation with input from RAP and JAM. MID wrote the original draft and curated the visualizations. RAP and JAM reviewed, edited, and provided critical feedback on the draft and all revisions.

## Footnotes

1 Reliability of parameter estimates at the individual level and thresholded activation maps at the group level have previously been distinguished as “reliability” and “reproducibility” of BOLD activity, respectively (Bennett & Miller, 2013; Plichta et al., 2012; Zuo et al., 2014). We elect to refer to individual and group estimates as distinct forms of reliability and use ‘reproducibility’ to refer to a broader set of concepts describing various aspects of the ability to reproduce or generalize a research finding (e.g. Goodman et al. [2016]).

2 For the Stage 1 submission, the data for the different studies was not fully accessed, inspected, preprocessed or analyzed. Thus, the sample size approximations. The final *N* for each sample is expected to deviate from the approximated values because of complete data availability and quality control exclusions.

3 At Stage 1 the sample was based on an approximation. During Stage 2, we realized it would be more effective to take advantage of the complete available data by using standardized effect Cohen’s *d* maps.

