## Supplemental Materials for "Impact of analytic decisions on test-retest reliability of individual and group estimates in functional magnetic resonance imaging: a multiverse analysis using the monetary incentive delay task"

### Section 1 – Analytic Decisions, FMRI Task, Data & Preprocessing

#### 1.1 Description of analytic decisions

The effect of smoothing is evaluated by selecting smoothing kernels that range from 1.5x - 3.5x the voxel size (in half point increments). This range is used in place of specific sized smoothing kernels (e.g., 4 mm) because the MLS and ABCD/AHRB differ in their voxel size, 4mm & 2.4mm, respectively. To avoid inflating the smoothing kernel in the MLS dataset, we scale the magnitude (e.g., voxel 4 mm x 2) by a magnitude of .60 (e.g., 2.4 mm/4 mm voxel size) for MLS data. The approximate smoothness between the two datasets is evaluated using Nipype's (Gorgolewski et al., 2018) interface of FSL's *SmoothEstimate()* applied to the model residuals to ensure the resulting smoothing in the BOLD data is comparable between the ABCD/AHRB and MLS samples. A range of liberal (e.g., no motion correction) to conservative strategies (e.g., censoring high motion volumes, excluding high motion subjects, and regressing estimated motion, their derivatives and eight anatomically derived noise components) are used to reduce the effects of motion and other artifacts that are historically acknowledged to increase variance in signal (Tomarken, 1995). Finally, over the years there have been several different modeling techniques for the MID task. For example, the cue phase (Demidenko et al., 2021; Srirangarajan et al., 2021) or fixation phase (Bjork et al., 2004; Sacchet & Knutson, 2013) may be modeled as the 'anticipation'. Below, **Figure S2**, suggests that these modeling decisions impact the efficiency of the design which may alter the variance structure across contrasts with lower and higher BOLD activity.

For demonstration purposes, the MID task events data from the AHRB study are used to generate the regressors for efficiency using the *neuRosim* package (Welvaert et al., 2011). Events information from 101 subjects (for this demonstration, some do not have the necessary outcome events which prevent the use of data in this case) is used for BOLD time series with a TR 800 ms and 407 volumes. The design of the task in the AHRB sample (as well as MLS/ABCD) is presented in **Figure S1**. The models that are calculated include different 'anticipation' model versions observed in the literature over the years (also included the 10-feedback variation duration regressors [hit/miss for each of the five cue types]):

- Cue Model: Cue onset + Cue Duration (2sec)
- Ant Model: Cue onset + (Cue Duration [2sec] + Fixation Duration [variable, 1.5-4sec])
- Fix Model: Fixation onset + Fixation Duration (variable, 1.5-4sec)

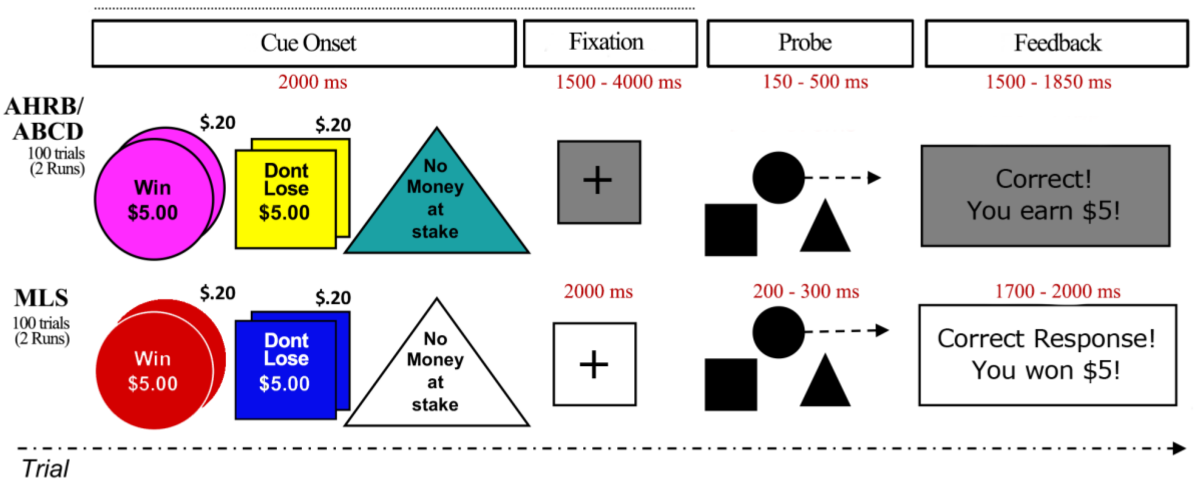


*Figure S1*. Task schematic for the AHRB, ABCD and MLS studies.

Schematic of the MID task design for the AHRB, ABCD and MLS samples. Both studies acquired 100 trials across two runs. Each task trial starts with a Cue indicating the trial time (Win [$5 or $0.20]; Lose [$5 or $0.20]; or Neutral). The cue lasts for 2000 ms. Following the cue is the Fixation cross. In the AHRB/ABCD samples, Fixation duration is variable (1500-4000 ms) but constant in MLS (2000 ms). The probe duration is a variable duration in all three samples. It is dependent on the participants performance. The probe window increases/decreases as the participants probe hit rate increases/decreases below a target of ~60%. The feedback phase of both the three studies is a variable duration and is adjusted based on the probe phase.

For the regressor estimates generated based on the provided behavioral data, efficiency can be calculated across model types. **Figure S2** displays the distribution and difference in estimated efficiency between the three model types across runs and the four contrasts for the Stage 1 Registered report (**NOTE**: in Stage 2 we learned of an error in *neuRosim* that impacted the interpretation of 'most efficient' model. See results in **Section 2.2** & **Figure S7**). These data suggest that across both runs the least efficient model is the Fixation Model (FixMod) and the most efficient model is the Cue Model (CueMod). While there is more similarity between the Anticipation Model (AntMod) and the Cue Model (CueMod), the latter in this is marginally better comparing vectors (via t-tests) as implemented in R using *ggsignif::geom_signif* (Ahlmann-Eltze & Patil, 2021). Efficiency is impacted by the modeled trial duration, number of trials, collinearity and other factors. The efficiency of a model's design matrix only reflects part of the first level model's variance, which is the product of the inverse of the efficiency and the residual variance. The most efficient design matrix may not fit the data well, increasing the residual variance and the overall variance of the estimated contrast. For example, consider CueMod and AntMod for the LGain v BL contrast. CueMod has higher efficiency due to lower overlap between the anticipation regressor (only modeled during Cue Onset + Cue Duration) and the Feedback regressor, but if the anticipation-based brain activation continues throughout the fixation period, CueMod will not capture this variability as well as AntMod. Whether CueMod outperforms AntMod for this contrast depends on whether the increased efficiency of CueMod is overshadowed by an increase in residual variance due to poor model fit.

The impact of model efficiency on reliability will be considered in parallel with how the residual variance estimate also varies. These modeling decisions may have an underlying impact on the underlying contrasts, as is shown in the figure below representing models across each run and contrast type. However, the impact on reliability estimates remains to be empirically tested across these different modeling approaches but one may hypothesize that the least efficient model (FixMod) and contrast (Small Gain v Neutral & Small Gain v Implicit Baseline) would have a lower reliability than the other models and contrasts.

- LGain: Large Gain > Neut
- SGain: Small Gain > Neut
- LGain v BL: Large Gain > Implicit Baseline
- SGain v BL: Small Gain > Implicit Baseline

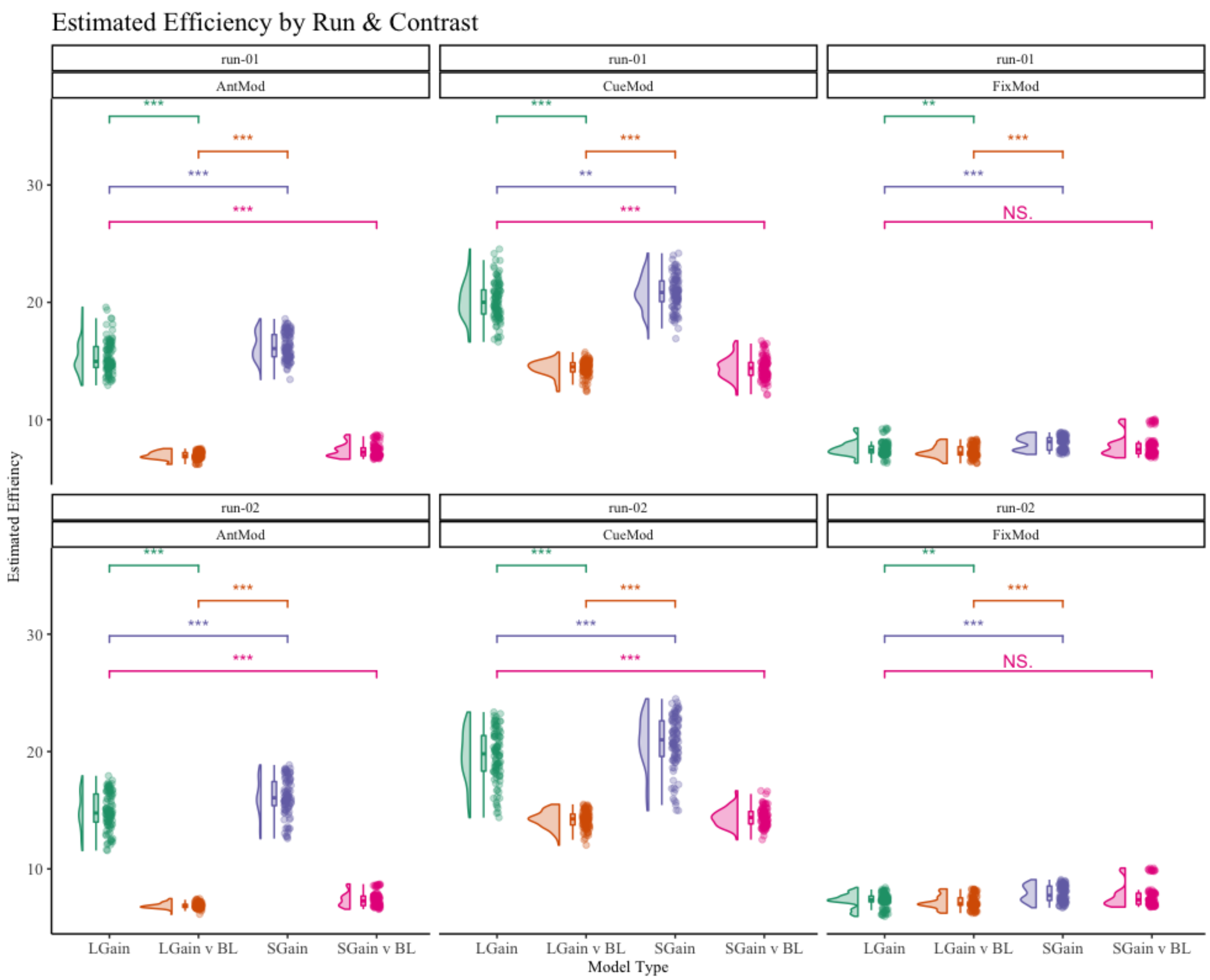


*Figure S2*: Modeling Efficiency Across Model, Run and Four MID Contrasts.

Comparing the model efficiencies between the four contrast types across the three model types. The Models are plotted for each run (run 01 and run 02) separately. LGain: Large Gain > Neut; SGain: Small Gain > Neut; LGain v BL: Large Gain > Implicit Baseline; SGain v BL: Small Gain > Implicit Baseline; CueMod: Cue onset + Cue Duration; AntMod: Cue onset + (Cue Duration + Fixation Duration; FixMod: Fixation onset + Fixation Duration. **Deprecated result:** We identified an error in neuRosim with how convolution is estimated. This does not impact other efficiency estimates as Nilearn is used in Stage 2 analyses.

### 1.2 Monetary Incentive Delay task description

As described elsewhere (Bjork, 2020; Demidenko et al., 2021; Knutson & Greer, 2008), the monetary incentive delay (MID) task measures reward anticipation. Apart from some minor differences, the MID task across the ABCD, AHRB and MLS samples are nearly identical. For example, during the MID task each trial starts with a cue type and consists of three phases: anticipation, probe and outcome (that is, feedback). The task regressors include different cue (five) and feedback types (ten), totaling 15-task regressors that are included in the GLM. **Table S1**, below, summarizes the trials, runs, cue types, timing and targeted accuracy information for the MID task across the three samples.

*Table S1*. Monetary Incentive Delay Task Details Across AHRB, ABCD and MLS samples.

| Sample | Trials | Runs | Cue Types (Trials) | Cue Duration (ms) | Fixation Duration (ms) | Probe Duration (ms) | Feedback Duration (ms) | Target Accuracy |
|---|---|---|---|---|---|---|---|---|
| AHRB | 50 | 2 | Win $5.00 (10), Win $0.20 (10), Neutral (10), Don’t Lose $5.00 (10), Don't Lose $0.20 (10) | 2000 | 1500 - 4000 | 150 - 500 | 1500 - 1850 | 60% |
| ABCD | 50 | 2 | Win $5.00 (10), Win $0.20 (10), Neutral (10), Don’t Lose $5.00 (10), Don't Lose $0.20 (10) | 2000 | 1500 - 4000 | 150 - 500 | 1500 - 1850 | 60% |
| MLS | 50 | 2 | Win $5.00 (10), Win $0.20 (10), Neutral (10), Don’t Lose $5.00 (10), Don't Lose $0.20 (10) | 2000 | 2000 | 300 - 500 | 1700 - 2000 | 60% |

### 1.3 FMRI Acquisition details

*Table S3*. Acquisition parameters for structural and functional data across *four* samples.

| | Scanner | Scan | TR (ms) | TE (ms) | Flip Angle | FOV (cm) | Voxel (mm) | Matrix |
|---|---|---|---|---|---|---|---|---|
| AHRB | GE MR750 | Structural | 7 | 2.9 | 8 | 25.6 | 1 | 256x256 |
| ABCD | GE MR750 | Structural | 2500 | 2 | 8 | 25.6 | 1 | 256x256 |
| | Philips | Structural | 6.31 | 2.9 | 8 | 25.6 | 1 | 256x256 |
| | Siemens | Structural | 2500 | 2.88 | 8 | 25.6 | 1 | 256x256 |
| MLS | GE Signa | Structural | 12 | 5.2 | 15 | 19.5 | 1.2 | 256x256 |

| | | | | | | | | |
|---|---|---|---|---|---|---|---|---|
| AHRB | GE MR750 | BOLD* | 800 | 30 | 52 | 21.6 | 2.4 | 90x90 |
| ABCD | GE MR750 | BOLD* | 800 | 30 | 52 | 21.6 | 2.4 | 90x90 |
| | Philips | BOLD* | 800 | 30 | 52 | 21.6 | 2.4 | 90x90 |
| | Siemens | BOLD* | 800 | 30 | 52 | 21.6 | 2.4 | 90x90 |
| MLS | GE Signa | BOLD | 2000 | 30 | 90 | 20 | 4 | 64x64 |

*BOLD runs are multiband 6 factor acquisition & Fieldmaps were collected. TR: Time Repetition; TE = Echo time; FOV: Field of view. ABCD & AHRB data are isotropic voxels (2.4 x2.4 x 2.4) and MLS data are anisotropic (3.125 x 3.125 x 4)

### 1.4. Preprocessing MRI & fMRI Data

Preprocessing of anatomical data. T1-weighted images are corrected for intensity non-uniformity (INU) with N4BiasFieldCorrection (Tustison et al., 2010), distributed with ANTs 2.3.3 (RRID:SCR_004757; Avants et al., 2008) and used as T1w-reference throughout the fMRIPrep workflow. The T1w-reference is then skull-stripped with a Nipype implementation of the antsBrainExtraction.sh workflow (from ANTs), using OASIS30ANTs as the target template. Brain tissue segmentation of cerebrospinal fluid (CSF), white-matter (WM) and gray-matter (GM) is performed on the brain-extracted T1w using fast (FSL 6.0.5.1:57b01774, RRID:SCR_002823; Zhang et al., 2001). Brain surfaces are reconstructed using recon-all (FreeSurfer 7.2.0, RRID:SCR_001847; Dale et al., 1999), and the brain mask estimated previously is refined with a custom variation of the method to reconcile ANTs-derived and FreeSurfer-derived segmentations of the cortical gray-matter of Mindboggle (RRID:SCR_002438; Klein et al., 2017). Volume-based spatial normalization to one standard space (MNI152NLin2009cAsym) is performed through nonlinear registration with antsRegistration (ANTs 2.3.3), using brain-extracted versions of both T1w reference and the T1w template. The following template are selected for spatial normalization: ICBM 152 Nonlinear Asymmetrical template version 2009c (RRID:SCR_008796; TemplateFlow ID: MNI152NLin2009cAsym; Fonov et al., 2009)

Preprocessing of functional data. For each of the 2 BOLD functional runs, the following preprocessing steps are performed. First, a reference volume and its skull-stripped version are generated using a custom methodology of fMRIPrep. The estimated fieldmap was then aligned with rigid-registration to the target EPI (echo-planar imaging) reference run. The field coefficients were mapped on to the reference EPI using the transform. The BOLD reference was then co-registered to the T1w reference using bbregister (FreeSurfer) which implements boundary-based registration (Greve & Fischl, 2009). Co-registration was configured with six degrees of freedom. The BOLD time-series were resampled into standard space, generating a preprocessed BOLD run in MNI152NLin2009cAsym space. Head-motion parameters with respect to the BOLD reference (transformation matrices, and six corresponding rotation and translation parameters) are estimated before any spatiotemporal filtering using mcflirt (FSL 6.0.5.1:57b01774; Jenkinson et al., 2002). The estimated fieldmap is then aligned with rigid-registration to the target EPI. Framewise displacement (FD) is calculated based on the preprocessed BOLD. Principal components are estimated after high-pass filtering the preprocessed BOLD time-series (using a discrete cosine filter with 128s cut-off) for anatomical (aCompCor). For the aCompCor decomposition, the k components with the largest singular values are retained, such that the retained components' time series are sufficient to explain 50 percent of variance across the nuisance mask (CSF, WM, combined, or temporal). The remaining components are dropped from consideration. The confounded time series derived from head motion estimates were expanded with the inclusion of temporal derivatives and quadratic terms for each (Satterthwaite et al., 2013). Frames that exceeded a threshold of 0.9 mm FD or 1.5 standardized DVARS were annotated as motion outliers.

### Section 2 – Results

The analytic code to recreate figures and estimates are available in the python notebooks and R markdown files shared in within the Stage 2 github repository. Specifically, the html reports include expanded information from the between-run and between-session HLM, emmeans, Specification Curves and other plots within the R html reports and may be recreated/reanalyzed using the share output files within the github Stage 2 repository.

### 2.1 Analytic modifications

For Aim 1b, instead of thresholding images by p < .001 (or *t*-stat 3.2) we converted the group *t*-stat to Cohen's *d* 3D effect size maps using the formula: $\frac{t\text{−}statistic}{\sqrt{N}}$. This is to avoid differences in *N*s between some models because of failures during preprocessing (e.g., *N* = 15 in ABCD failed aCompCor WM/GM/CSF masks).

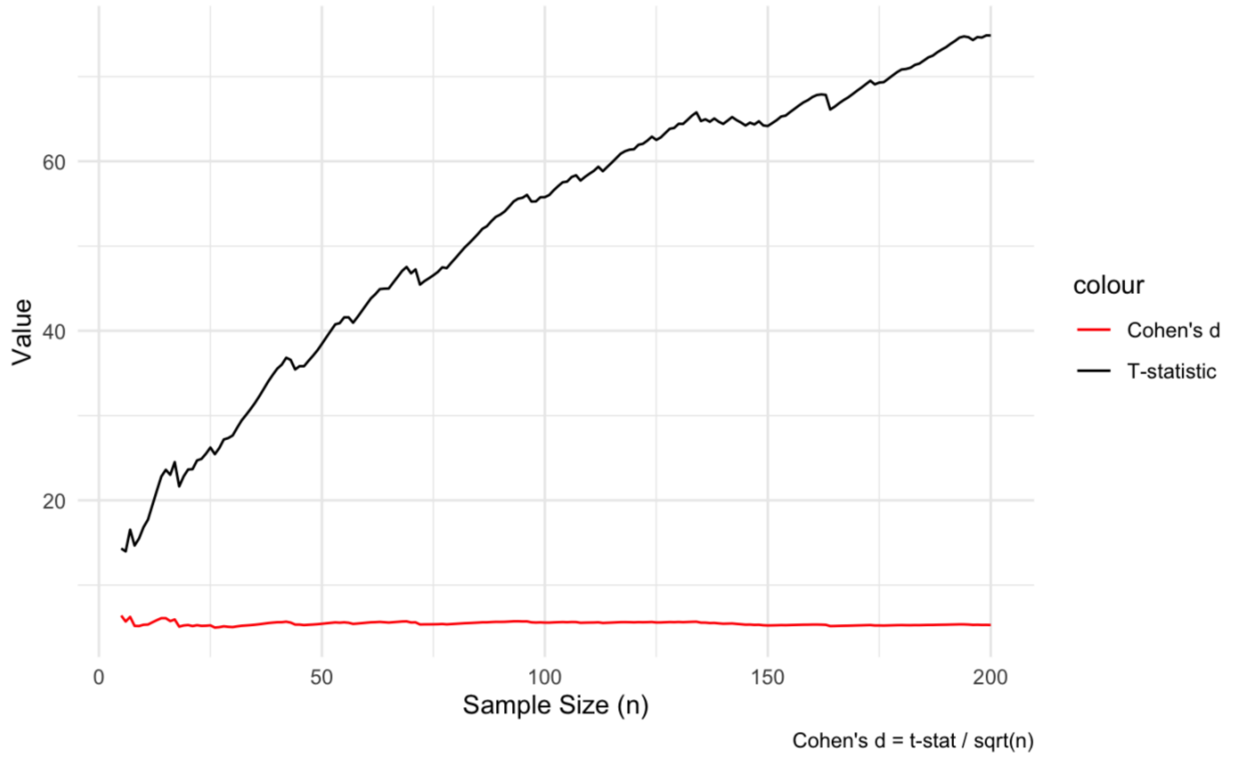


*Figure S3*: Change in *t*-statistic and Cohen's *d* across N = 0 to N = 200 in a randomly simulated data with $\mu$ = 5 and $\sigma$= 1. The population mean for *t*-test is assumed to be zero.

We ran the model permutations on the ABCD/AHRB (2.4mm data) and MLS (4mm) data with a weighted .50 FWHM smoothing parameter, we estimated the smoothness of the *group residual variance* maps for the data. Since the model permutations differed in several decisions, the smoothness is estimated across the 240 pipelines spanning four contrasts, four motion options, three model parameterizations and five smoothness parameters. The estimated *average* smoothness (Resel^[⅓]) for the ABCD 4.5 (SD = 1.4), AHRB 4.2 (SD = 1.3) and MLS 3.8 (SD

= 1.0). The distribution of estimated smoothness across group-level maps for the ABCD, AHRB and MLS data are reported in **Figure S4**.

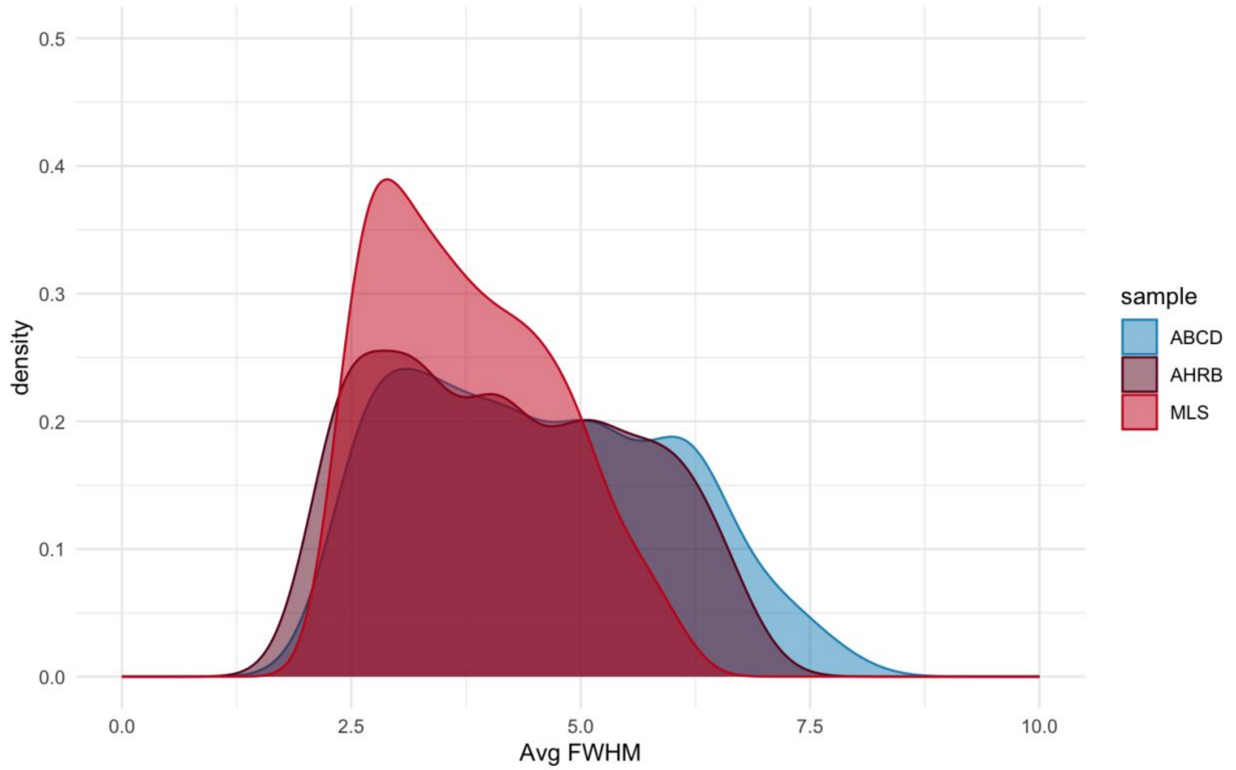


*Figure S4*: Estimates of smoothing of group level residual 3D volumes across 240 permutations for the Michigan Longitudinal (MLS), Adolescent Health Risk Behavior (AHRB) and Adolescent Brain Cognitive Development (ABCD) imaging data.

### 2.2 Descriptive Results

*Demographics Across Samples*: The demographic information is reported in **Table S4** and the days between sessions are visually represented in **Figure S5**.

*Table S4*. Age, Sex, Race/Ethnicity from Session 1 and Days Between Sessions Across ABCD, AHRB and MLS

| | **ABCD (N=119)** | **AHRB (N=60)** | **MLS (N=81)** |
|---|---|---|---|
| | | *Mean (SD)* | |
| **Age** | 9.8 (0.6) | 19.3 (1.3) | 20.7 (2.3) |
| **Days Btwn Session** | 747 (79.1) | 419 (80.1) | 1090 (624) |
| **Sex** | | *N* (%) | |
| Female | 58 (48.7%) | 35 (58.3%) | 31 (38.3%) |
| Male | 61 (51.3%) | 25 (41.7%) | 50 (61.7%) |
| **Race/Ethnicity** | | | |
| Asian | 4 (3.4%) | 0 (0%) | 0 (0%) |
| Black | 14 (11.8%) | 10 (16.7%) | 2 (2.5%) |
| Hispanic | 8 (6.7%) | 3 (5.0%) | 5 (6.2%) |
| Other | 15 (12.6%) | 5 (8.3%) | 1 (1.2%) |
| White | 78 (65.5%) | 42 (70.0%) | 73 (90.1%) |

Note: *MLS* participants reported on “caucasian”,”African American”, “Native American”, “Asian American”, “Filipino or Pacific Islander”, “Bi-Racial” and “Hispanic-caucasian race”, and *AHRB* “White Non-Hispanic”, “Black Non-Hispanic”, “Hispanic/Latinx”, and “american Indian/Alaska/Native Hawaiian”, “Other” for simplicity refactor to match ABCD “Race/Ethnicity” variable in *acspsw03*

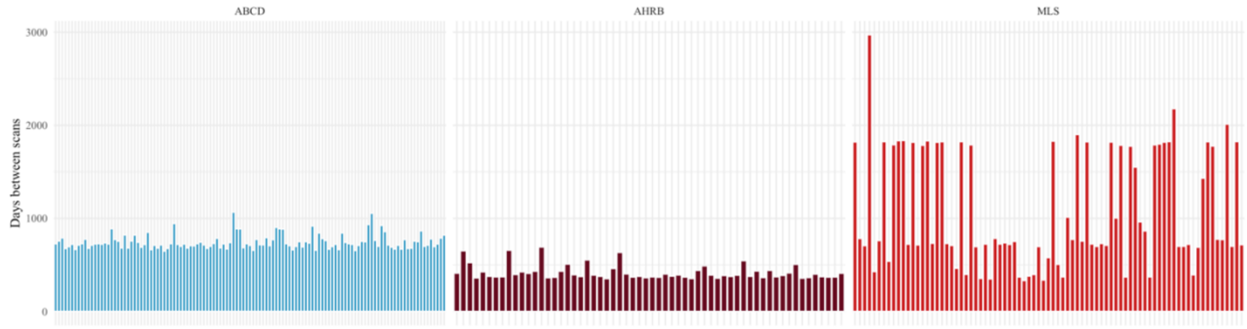


*Figure S5.* The number of days between sessions for subjects across ABCD, AHRB and MLS samples.

*Task Behavior Across Samples*: The Mean and Standard Deviation for the run average Mean Framewise Displacement, Average Probe Response Times and Average Probe Accuracies are reported in **Table S5** and **Figure S6**.

*Table S5*: The run average Mean FD, Average Probe Accuracy and Mean RT across samples and sessions.

| Sample | Session | Mean | SD | Min | Max |
|---|---|---|---|---|---|
| *Mean Framewise Displacement* | | | | | |
| ABCD | 1 | 0.25 | 0.15 | 0.06 | 0.86 |
| AHRB | 1 | 0.12 | 0.04 | 0.05 | 0.24 |
| MLS | 1 | 0.10 | 0.03 | 0.04 | 0.25 |
| ABCD | 2 | 0.25 | 0.23 | 0.05 | 1.29 |
| AHRB | 2 | 0.14 | 0.08 | 0.06 | 0.51 |
| MLS | 2 | 0.09 | 0.03 | 0.05 | 0.21 |
| *Average Probe Accuracy (%)* | | | | | |
| ABCD | 1 | 0.55 | 0.04 | 0.44 | 0.63 |
| AHRB | 1 | 0.57 | 0.04 | 0.48 | 0.66 |
| MLS | 1 | 0.72 | 0.13 | 0.40 | 0.94 |
| ABCD | 2 | 0.56 | 0.04 | 0.44 | 0.63 |
| AHRB | 2 | 0.58 | 0.03 | 0.51 | 0.65 |
| MLS | 2 | 0.67 | 0.12 | 0.37 | 0.94 |
| *Average probe MRT (ms)* | | | | | |
| ABCD | 1 | 306.8 | 34.5 | 233.5 | 406.2 |
| AHRB | 1 | 297.1 | 18.5 | 236.8 | 337.8 |
| MLS | 1 | 204.3 | 28.9 | 146.8 | 268.2 |

| | | | | | |
|---|---|---|---|---|---|
| ABCD | 2 | 274.3 | 34.1 | 214.3 | 439.3 |
| AHRB | 2 | 248.5 | 21.5 | 217.6 | 313.3 |
| MLS | 2 | 210.1 | 30.0 | 105.8 | 277.4 |

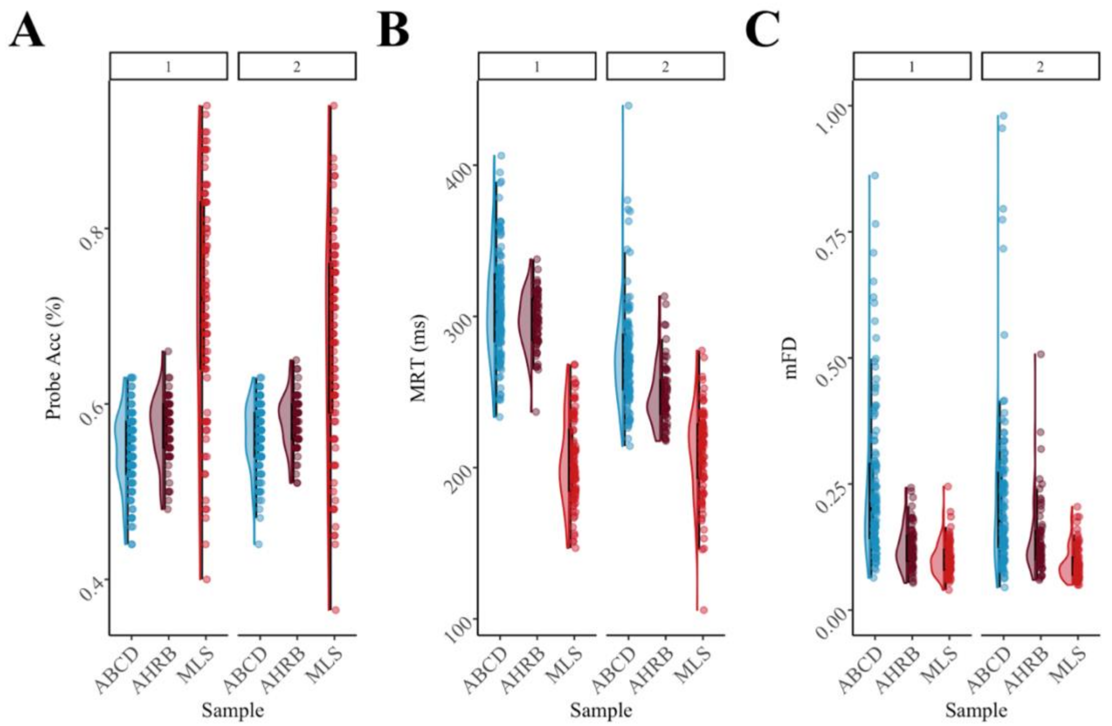


186 *Figure S6.* Distribution of (A) Mean Framewise Displacement, (B) Mean Probe RTs (ms) and
187 (C) Mean Probe Accuracy (%) across Sessions and ABCD, AHRB and MLS samples.

*Task Efficiency Across Samples*: The model efficiency was calculated as the inverse proportion of variance based on the design matrix. The design matrix varied only as a function of parameterization and motion regressors for the four contrasts. The formula used is: $\boldsymbol{Efficiency} = \frac{\mathbf{1}}{\boldsymbol{c}(\boldsymbol{X}'\boldsymbol{X})^{-\mathbf{1}}\boldsymbol{c}'}$. As is observed from **Figure S7**, contrary to the above/incorrect *neuRosim* **Figure S2**, the most efficient design (compared within a category) is the Anticipation Model ('AntMod'). Furthermore, consistent with our hypothesis, the most efficient contrast within a model is the *Large Gain* versus *Neutral* contrast.

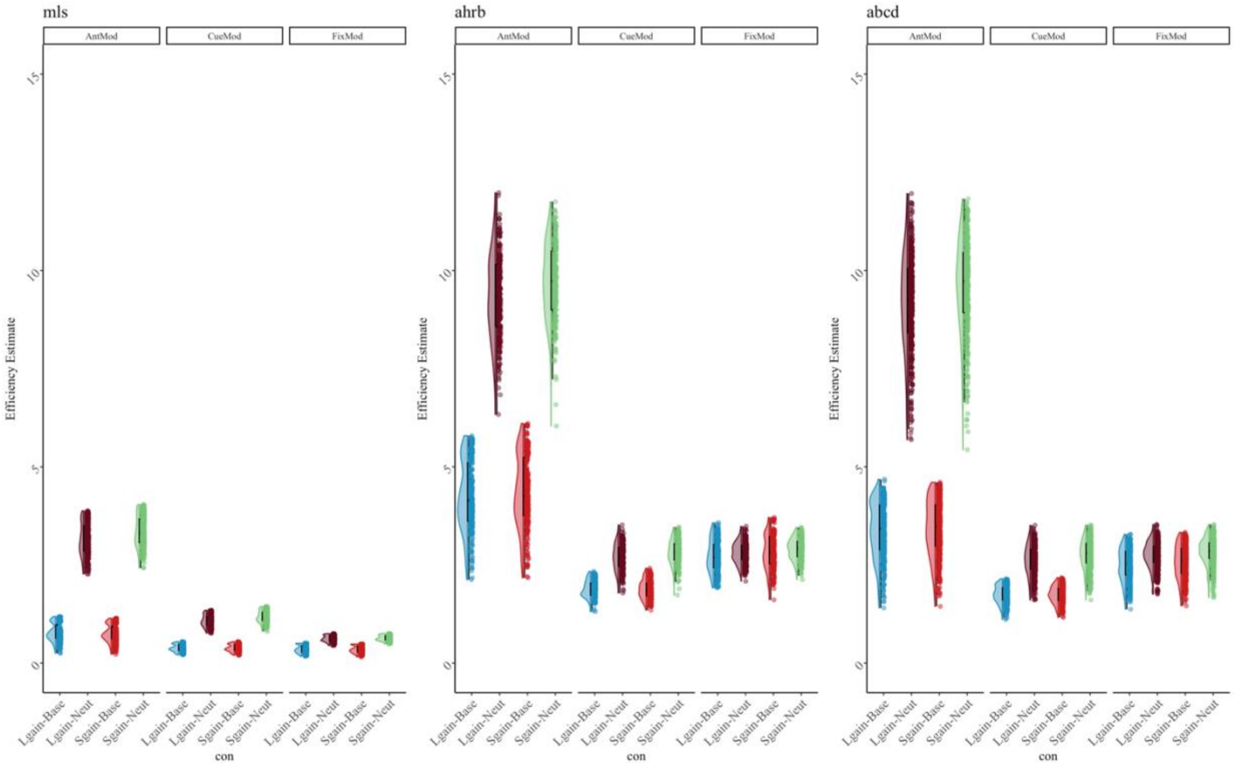


*Figure S7*. Distribution of estimated model efficiencies from design matrices for Model Parameterization and Contrast type across ABCD, AHRB and MLS samples.

*Between-run and Between-session similarity estimates:* Overall, the between-session ICC, Jaccard and Spearman Similarity estimates were higher than the Session 1 between-run estimates (**Table S5**).

*Table S5*. Session 1 Between-run and Between-session Median, Mean, Standard Deviation (SD), Minimum and Maximum of median Intraclass Correlation Coefficient (ICC) and Jaccard and Spearman Similarity and from 240 analytic models across ABCD, AHRB and MLS Samples.

| study | estimate | median | mean | sd | min | max |
|---|---|---|---|---|---|---|
| | | *Session 1: Between-runs* | | | | |
| ABCD | ICC* | .11 | .15 | .12 | -.07 | .43 |
| AHRB | ICC* | .18 | .20 | .13 | .00 | .52 |
| MLS | ICC* | .18 | .21 | .13 | .04 | .55 |
| ABCD | Jaccard | .09 | .11 | .09 | .01 | .45 |
| AHRB | Jaccard | .18 | .21 | .15 | .01 | .64 |
| MLS | Jaccard | .34 | .34 | .11 | .15 | .60 |
| ABCD | Spearman* | .68 | .68 | .14 | .35 | .89 |
| AHRB | Spearman* | .73 | .68 | .22 | .22 | .96 |
| MLS | Spearman* | .84 | .80 | .12 | .47 | .95 |
| | | *Between-sessions* | | | | |
| ABCD | ICC* | .15 | .16 | .07 | .03 | .34 |
| AHRB | ICC* | .21 | .23 | .13 | .04 | .53 |
| MLS | ICC* | .21 | .22 | .10 | .06 | .47 |
| ABCD | Jaccard | .25 | .26 | .13 | .02 | .61 |
| AHRB | Jaccard | .30 | .32 | .19 | .04 | .73 |
| MLS | Jaccard | .42 | .43 | .12 | .20 | .74 |
| ABCD | Spearman* | .80 | .76 | .13 | .40 | .94 |
| AHRB | Spearman* | .82 | .74 | .21 | .32 | .97 |
| MLS | Spearman* | .87 | .85 | .09 | .59 | .97 |

*Supra-threshold mask

### 2.3 Aim 1 results

*A. Between-Run Individual Reliability*:

The average and standard deviation across model permutations for each sample are reported in **Figure S8**. The distribution of median ICC estimates across [four] analytic options is reported in **Figure S9**. The complete supra-threshold specification curve for between-run median ICCs are reported in **Figure S9** and the sub-threshold in **Figure S11**.

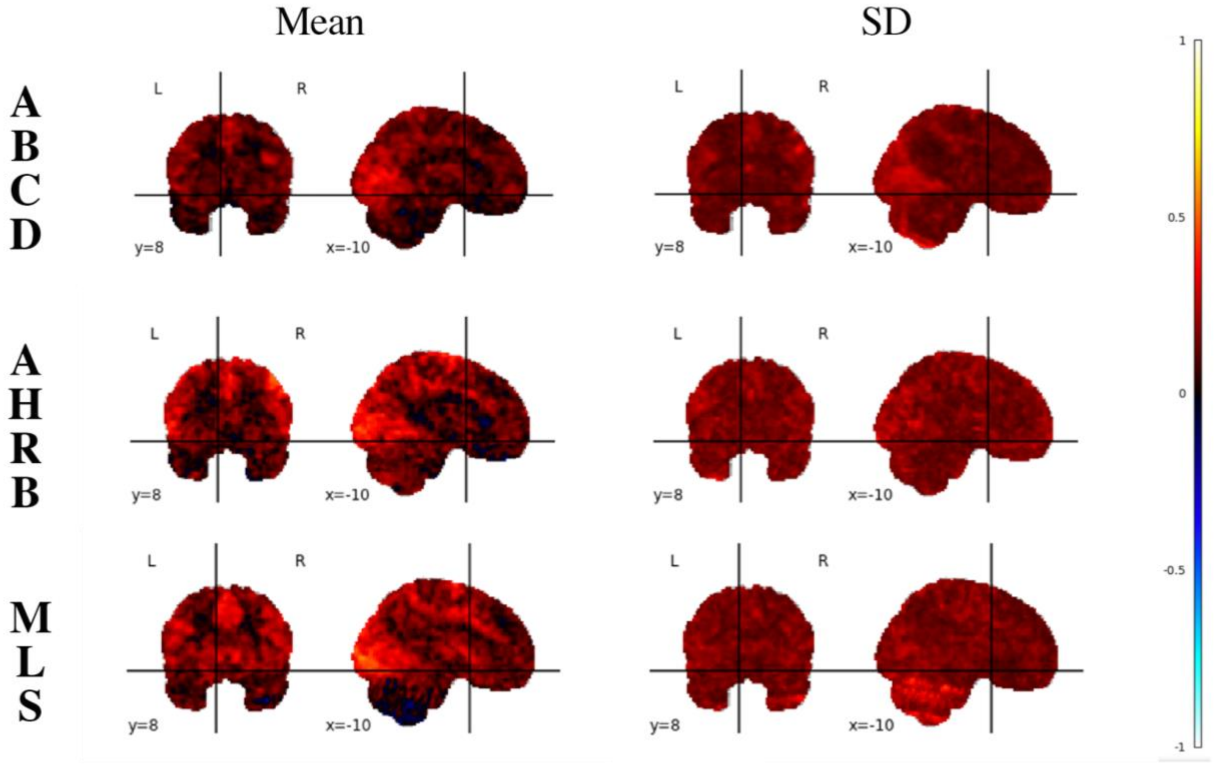


*Figure S8*: Mean and SD of *ICC estimates* across 240 permutations for the Adolescent Brain Cognitive Development (ABCD), Adolescent Health Risk Behavior (AHRB) and Michigan Longitudinal (MLS) 3D volumes.

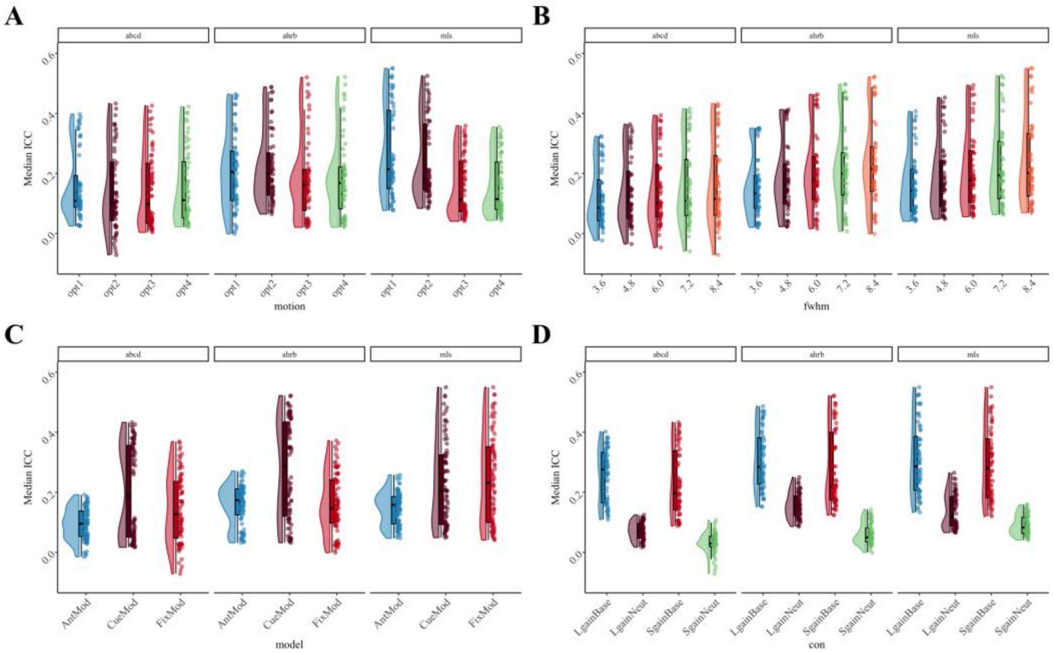


*Figure S9.* Supra-threshold Median ICC Session 1 between-run reliability estimates for (A) Motion, (B) FWHM, (C) Model Paramterization and (D) Contrasts analytic options across the ABCD, AHRB and MLS samples. Expanded version of in-text Figure 2.

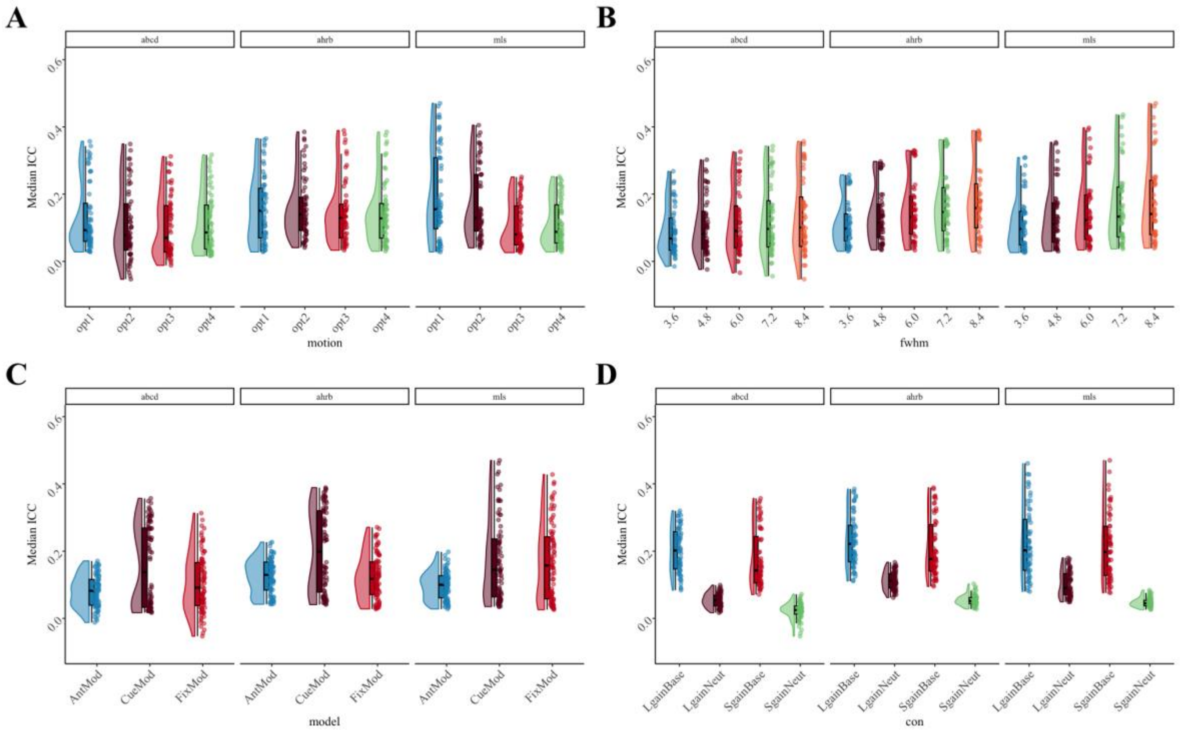


*Figure S10.* Sub-threshold Median ICC Session 1 between-run reliability estimates for Contrast (con) and Model Parameterization analytic options across the ABCD, AHRB and MLS samples.

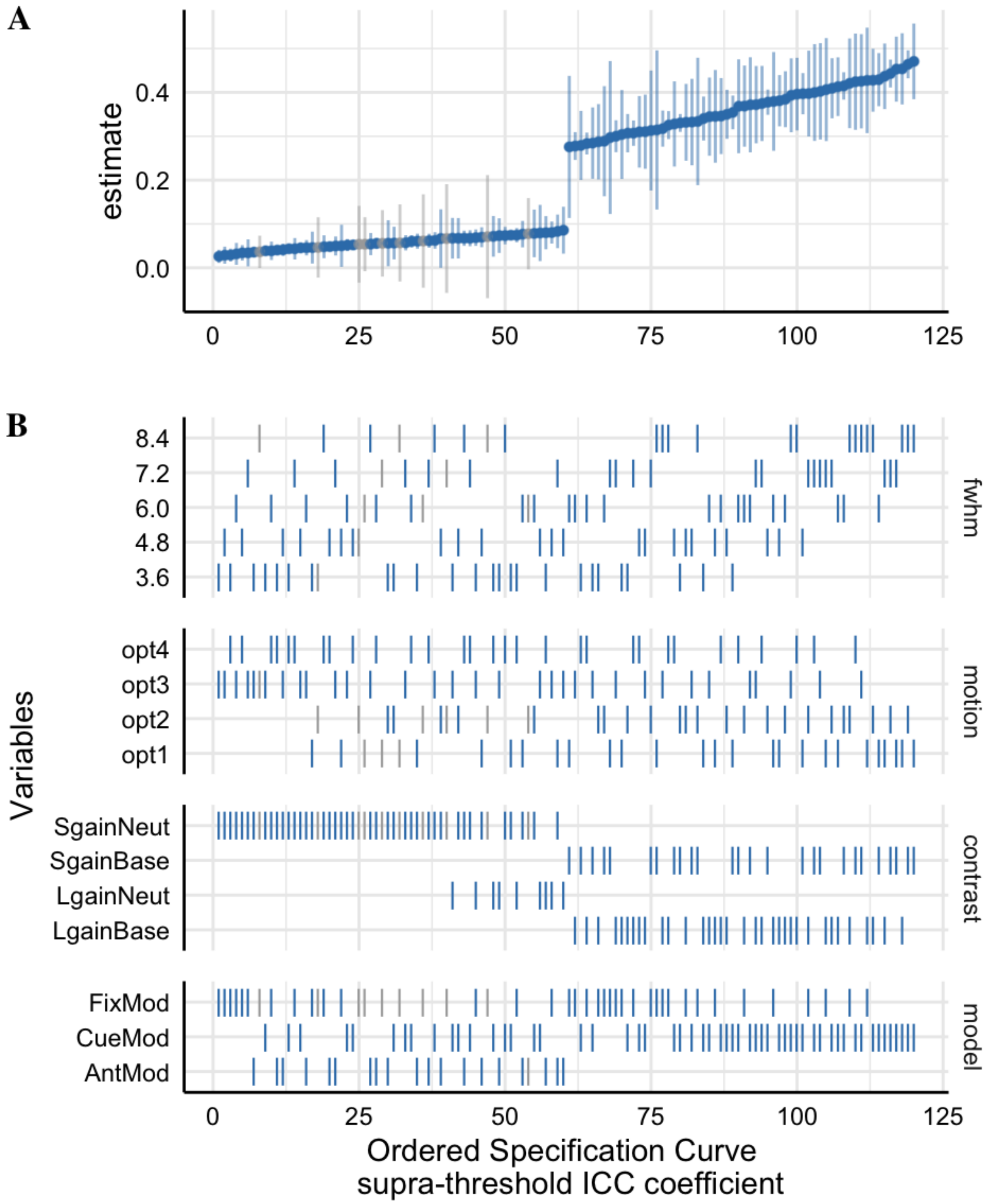


*Figure S11*: The 25th and 75th percentile supra-threshold Specification Curve of the Session 1 Between-run Median ICC estimates across 240 pipeline permutations for the ABCD, AHRB and MLS samples. Full length of estimates reported in **Figure 4.**
A. The distribution of the point estimate (average) and distribution (error bars) across the three samples. B. The model options (four) associated with each estimate.

*Table S6*: Tukey's HSB Estimate Means Differences for between-run Supra-threshold ICC Model Parameters in-text Table 3.

| Contrast | Est | SE | Low.CI | Up.CI | *p* |
|---|---|---|---|---|---|
| fwhm3.6 - fwhm4.8 | -.02 | .01 | -.04 | .00 | .023 |
| fwhm3.6 - fwhm6.0 | -.04 | .01 | -.06 | -.02 | .000 |
| fwhm3.6 - fwhm7.2 | -.06 | .01 | -.08 | -.04 | .000 |
| fwhm3.6 - fwhm8.4 | -.07 | .01 | -.09 | -.05 | .000 |
| fwhm4.8 - fwhm6.0 | -.02 | .01 | -.04 | .00 | .098 |
| fwhm4.8 - fwhm7.2 | -.03 | .01 | -.05 | -.01 | .000 |
| fwhm4.8 - fwhm8.4 | -.04 | .01 | -.06 | -.02 | .000 |
| fwhm6.0 - fwhm7.2 | -.01 | .01 | -.04 | .01 | .299 |
| fwhm6.0 - fwhm8.4 | -.03 | .01 | -.05 | -.01 | .006 |
| fwhm7.2 - fwhm8.4 | -.01 | .01 | -.03 | .01 | .575 |
| LgainBase - LgainNeut | .17 | .01 | .15 | .19 | .000 |
| LgainBase - SgainBase | .02 | .01 | .01 | .04 | .001 |
| LgainBase - SgainNeut | .23 | .01 | .21 | .25 | .000 |
| LgainNeut - SgainBase | -.14 | .01 | -.16 | -.13 | .000 |
| LgainNeut - SgainNeut | .06 | .01 | .04 | .08 | .000 |
| SgainBase - SgainNeut | .21 | .01 | .19 | .22 | .000 |
| opt1 - opt2 | .01 | .01 | -.01 | .03 | .283 |
| opt1 - opt3 | .05 | .01 | .03 | .07 | .000 |
| opt1 - opt4 | .05 | .01 | .03 | .06 | .000 |
| opt2 - opt3 | .04 | .01 | .02 | .06 | .000 |
| opt2 - opt4 | .03 | .01 | .02 | .05 | .000 |
| opt3 - opt4 | .00 | .01 | -.02 | .01 | .940 |
| AntMod - CueMod | -.10 | .01 | -.12 | -.09 | .000 |
| AntMod - FixMod | -.05 | .01 | -.06 | -.04 | .000 |
| CueMod - FixMod | .05 | .01 | .04 | .07 | .000 |

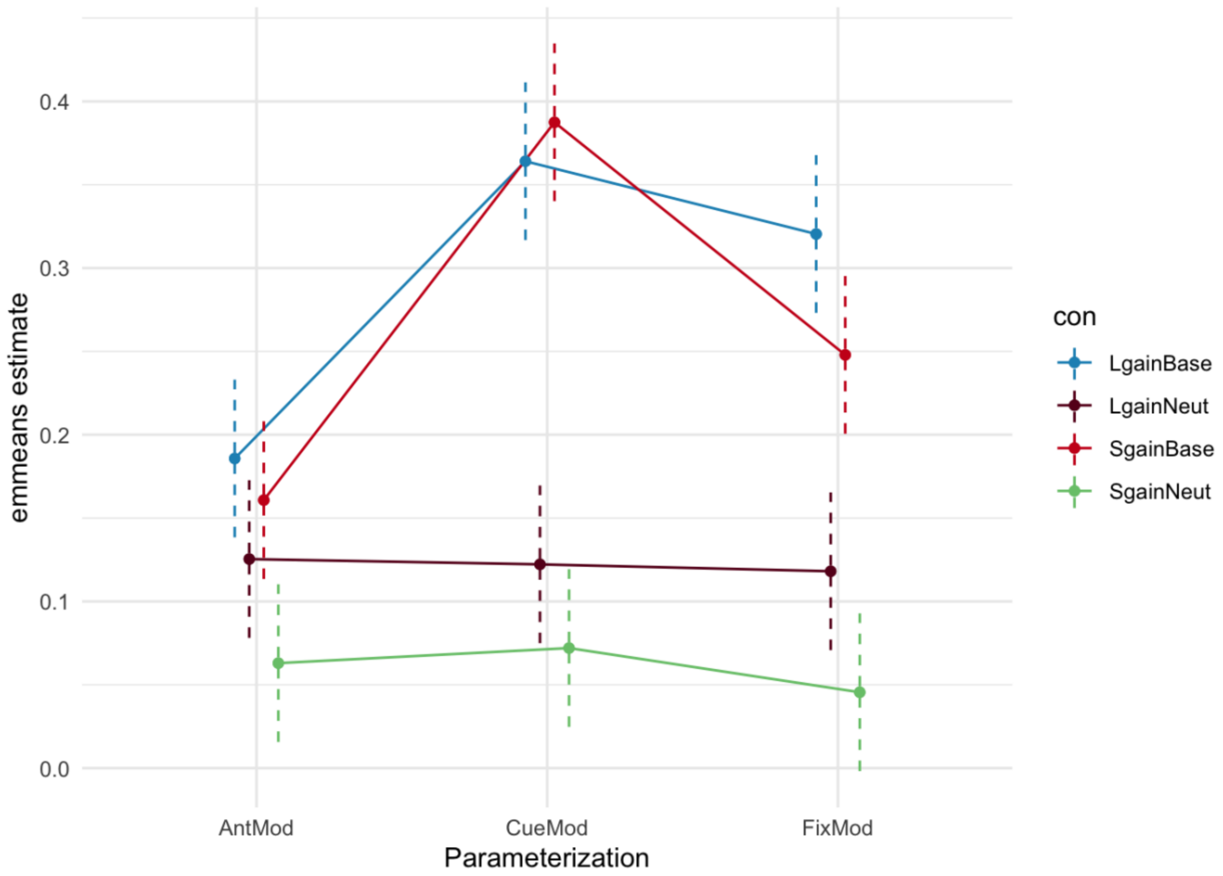


*Figure S12*: Median ICC estimate: Interaction plot of *emmeans* fitted model of Contrast-by-Model parameterization for Session 1 Between-run supra-threshold estimates using *emmip()*. Point estimate is a linear median ICC estimate from *emmeans* function. Dashed bars are estimated confidence intervals by *emmeans*.

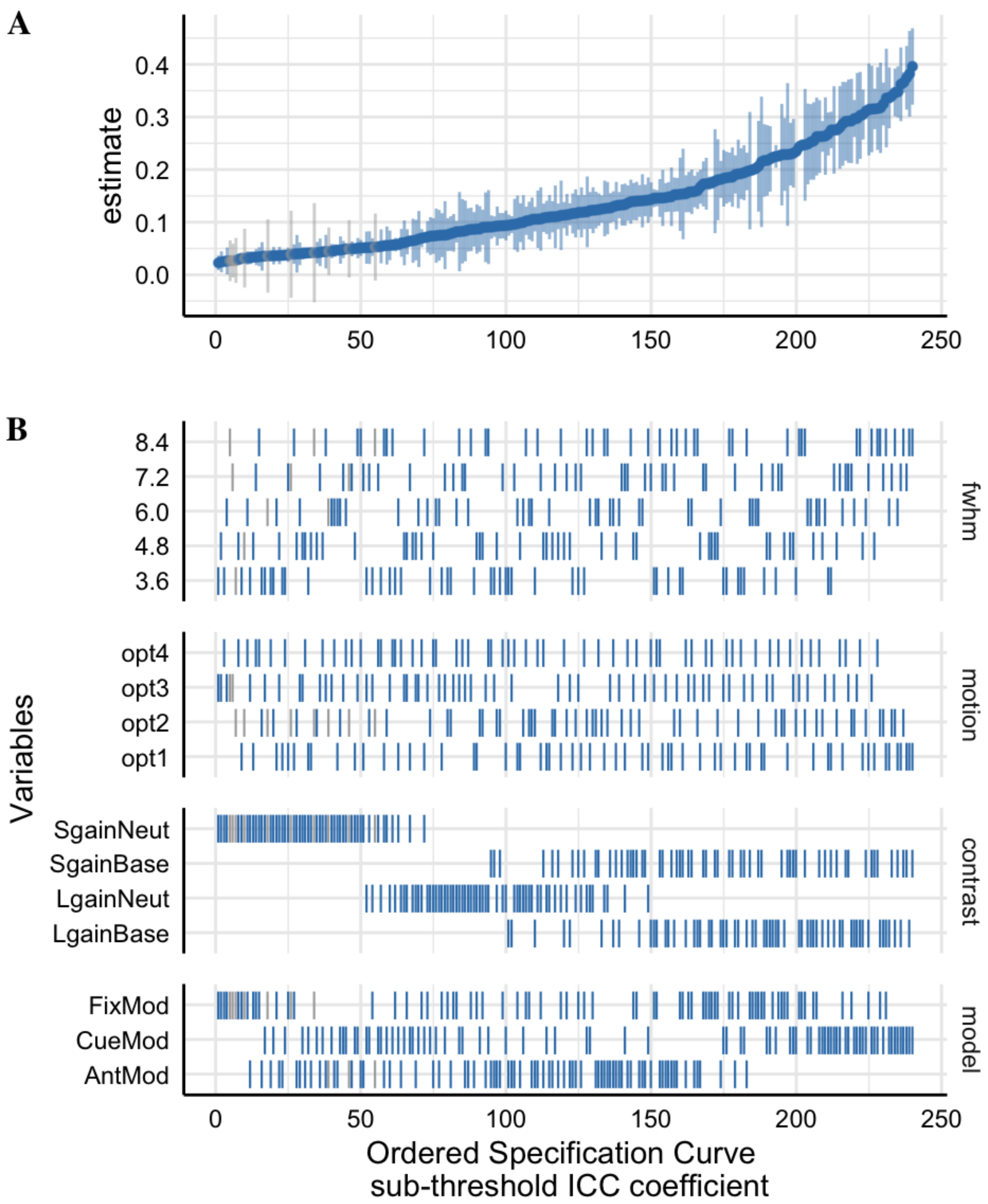


*Figure S13*: The sub-threshold Specification Curve of the Median Intraclass Correlation Coefficient (ICC[3,1]) estimates across 240 pipeline permutations for the ABCD, AHRB and MLS estimate.

A. The distribution of the point estimate (average) across the three studies and distribution across the three samples.

B. The model options (four) associated with each estimate.

*Table S7*: Hierarchical Linear Model: (A) Linear associations between the analytic decisions and the *Session 1 Between-run* median Intraclass Correlation Coefficient (ICC[3,1]), Between-subject (BS) and Within-subject variance (WS) from **sub-threshold mask** and (B) the impact of the analytic category on the marginal $R^2$.

**A. HLM Estimates for Sub-threshold Mask**

| | Median ICC | | | Median BS | | | Median WS | | |
|---|---|---|---|---|---|---|---|---|---|
| *Predictors* | *b* | *CI* | *p* | *b* | *CI* | *p* | *b* | *CI* | *p* |
| (Intercept) | .17 | .15 – .20 | <.001 | .31 | .21 – .41 | <.001 | 1.34 | 1.05 – 1.64 | <.001 |
| Reference [3.6] | | | | | | | | | |
| fwhm [4.8] | .02 | .01 – .03 | .001 | -.02 | -.06 – .01 | .18 | -.35 | -.42 – -.28 | <.001 |
| fwhm [6.0] | .03 | .02 – .05 | <.001 | -.04 | -.08 – -.01 | .02 | -.55 | -.62 – -.48 | <.001 |
| fwhm [7.2] | .05 | .04 – .06 | <.001 | -.06 | -.09 – -.02 | .002 | -.67 | -.74 – -.60 | <.001 |
| fwhm [8.4] | .06 | .05 – .07 | <.001 | -.07 | -.10 – -.03 | <.001 | -.75 | -.82 – -.68 | <.001 |
| Reference [opt1] | | | | | | | | | |
| motion [opt2] | -.02 | -.03 – -.01 | .003 | -.07 | -.10 – -.04 | <.001 | -.14 | -.21 – -.08 | <.001 |
| motion [opt3] | -.04 | -.05 – -.03 | <.001 | -.14 | -.17 – -.11 | <.001 | -.29 | -.35 – -.23 | <.001 |
| motion [opt4] | -.04 | -.05 – -.03 | <.001 | -.14 | -.17 – -.11 | <.001 | -.30 | -.36 – -.24 | <.001 |
| Reference [AntMod] | | | | | | | | | |
| model [CueMod] | .08 | .07 – .08 | <.001 | .18 | .15 – .20 | <.001 | .34 | .29 – .40 | <.001 |
| model [FixMod] | .03 | .02 – .04 | <.001 | .13 | .10 – .15 | <.001 | .38 | .33 – .44 | <.001 |
| Reference [LgainBase] | | | | | | | | | |
| con [LgainNeut] | -.13 | -.14 – -.12 | <.001 | -.25 | -.28 – -.22 | <.001 | -.46 | -.52 – -.40 | <.001 |
| con [SgainBase] | -.02 | -.03 – -.01 | <.001 | -.03 | -.06 – .01 | .12 | .01 | -.06 – .07 | .84 |
| con [SgainNeut] | -.18 | -.19 – -.17 | <.001 | -.27 | -.31 – -.24 | <.001 | -.49 | -.55 – -.43 | <.001 |

**B. Analytic Category Model Impact**

| **Comparison** | χ2 | Orig R2 | New R2 | ΔR2 | χ2 | Orig R2 | New R2 | ΔR2 | χ2 | Orig R2 | New R2 | ΔR2 |
|---|---|---|---|---|---|---|---|---|---|---|---|---|
| [Full] vs [New - fwhm] | 123 | .73 | .69 | .04 | 16 | .45 | .44 | .01 | 428 | .53 | .31 | .22 |
| [Full] vs [New - motion] | 84 | .73 | .71 | .02 | 94 | .45 | .39 | .06 | 115 | .53 | .49 | .04 |
| [Full] vs [New - model] | 252 | .73 | .63 | .10 | 147 | .45 | .36 | .09 | 209 | .53 | .44 | .09 |
| [Full] vs [New - con] | 867 | .73 | .17 | .56 | 362 | .45 | .17 | .28 | 360 | .53 | .36 | .17 |

*Table S8*: Tukey's HSB Estimate Means Differences for Sub-threshold Between-run ICC Model Parameters in Table S6.

| Contrast | Est | SE | Low.CI | Up.CI | *p* |
|---|---|---|---|---|---|
| fwhm3.6 - fwhm4.8 | -.02 | .01 | -.03 | .00 | .013 |
| fwhm3.6 - fwhm6.0 | -.03 | .01 | -.05 | -.02 | .000 |
| fwhm3.6 - fwhm7.2 | -.05 | .01 | -.06 | -.03 | .000 |
| fwhm3.6 - fwhm8.4 | -.06 | .01 | -.07 | -.04 | .000 |
| fwhm4.8 - fwhm6.0 | -.02 | .01 | -.03 | .00 | .044 |
| fwhm4.8 - fwhm7.2 | -.03 | .01 | -.04 | -.01 | .000 |
| fwhm4.8 - fwhm8.4 | -.04 | .01 | -.06 | -.02 | .000 |
| fwhm6.0 - fwhm7.2 | -.01 | .01 | -.03 | .00 | .134 |
| fwhm6.0 - fwhm8.4 | -.02 | .01 | -.04 | -.01 | .000 |
| fwhm7.2 - fwhm8.4 | -.01 | .01 | -.03 | .00 | .317 |
| LgainBase - LgainNeut | .13 | .01 | .12 | .14 | .000 |
| LgainBase - SgainBase | .02 | .01 | .01 | .03 | .000 |
| LgainBase - SgainNeut | .18 | .01 | .16 | .19 | .000 |
| LgainNeut - SgainBase | -.11 | .01 | -.12 | -.09 | .000 |
| LgainNeut - SgainNeut | .05 | .01 | .04 | .06 | .000 |
| SgainBase - SgainNeut | .16 | .01 | .14 | .17 | .000 |
| opt1 - opt2 | .02 | .01 | .00 | .03 | .018 |
| opt1 - opt3 | .04 | .01 | .03 | .05 | .000 |
| opt1 - opt4 | .04 | .01 | .02 | .05 | .000 |
| opt2 - opt3 | .03 | .01 | .01 | .04 | .000 |
| opt2 - opt4 | .02 | .01 | .01 | .04 | .000 |
| opt3 - opt4 | .00 | .01 | -.02 | .01 | .913 |
| AntMod - CueMod | -.08 | .00 | -.09 | -.07 | .000 |
| AntMod - FixMod | -.03 | .00 | -.04 | -.02 | .000 |
| CueMod - FixMod | .04 | .00 | .03 | .06 | .000 |

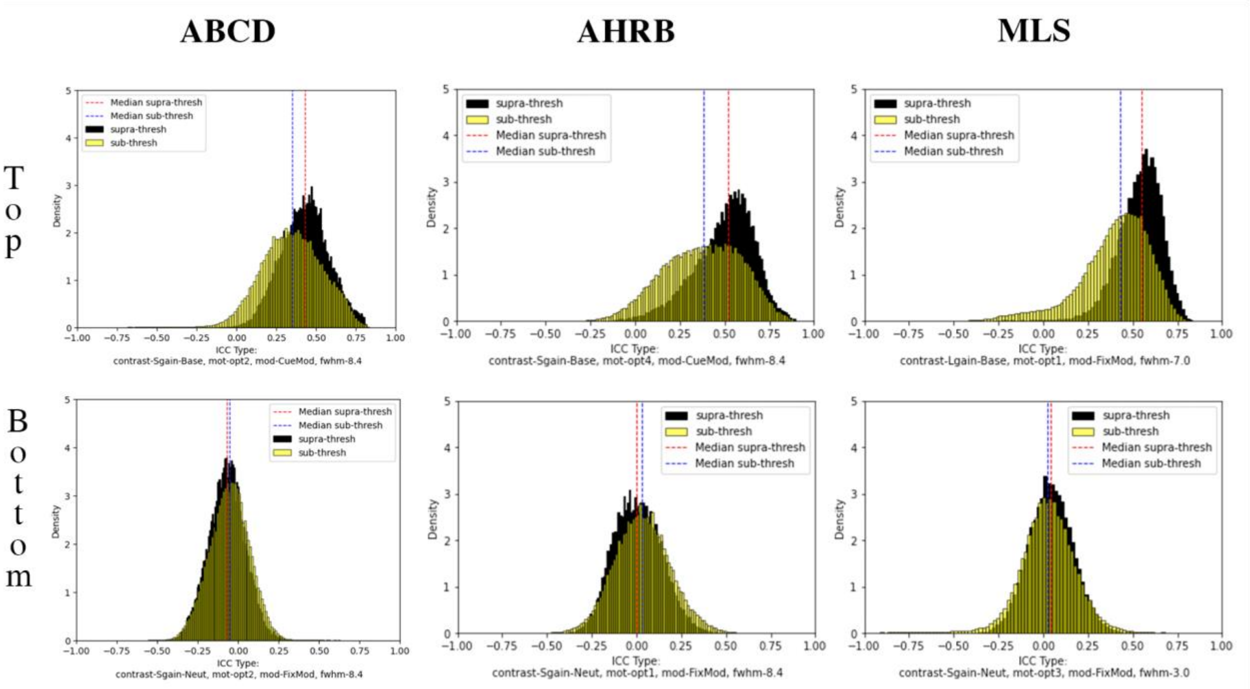


*Figure S14*: Voxelwise Distribution of ICCs for Supra- and Sub-threshold mask for highest (Top) and Lowest (Bottom) estimates from in-text Figure 3 and Figure S13 Across ABCD, AHRB and MLS samples.

*A. Between-Session Individual Reliability:*

The mean and standard deviation (**Figure S15**) of the 3D volumes across the 240 analytic decisions illustrate a consistent pattern, whereby the highest nose is within CSF and high noise regions across the three samples. Consistent with the Session 1 between-run median ICC estimates, variability in the median ICC estimate across 240 pipelines and three samples is best explained by contrast (marginal $\Delta R^2$ : .51) and model parameterization (marginal $\Delta R^2$ : .07), **see Table S9**. Compared to the between-run, the FWHM had a higher impact on the between-session model fit (marginal $\Delta R^2$: .06) but motion remained negligible (marginal $\Delta R^2$ : .02). Like the between-run estimates, the *Implicit Baseline* is the main contributor to the model parameterization differences (**Figure S17**).

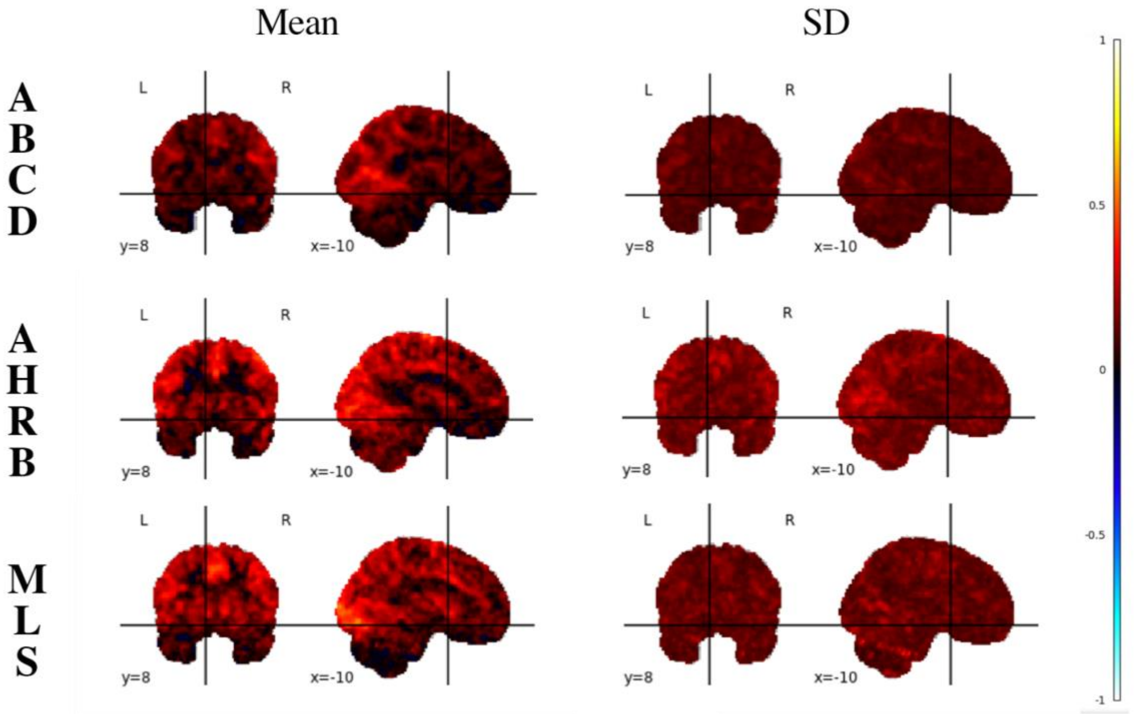


*Figure S15*: Mean and SD of ICC estimates across 240 permutations for the Adolescent Brain Cognitive Development (ABCD), Adolescent Health Risk Behavior (AHRB) and Michigan Longitudinal (MLS) 3D volumes.

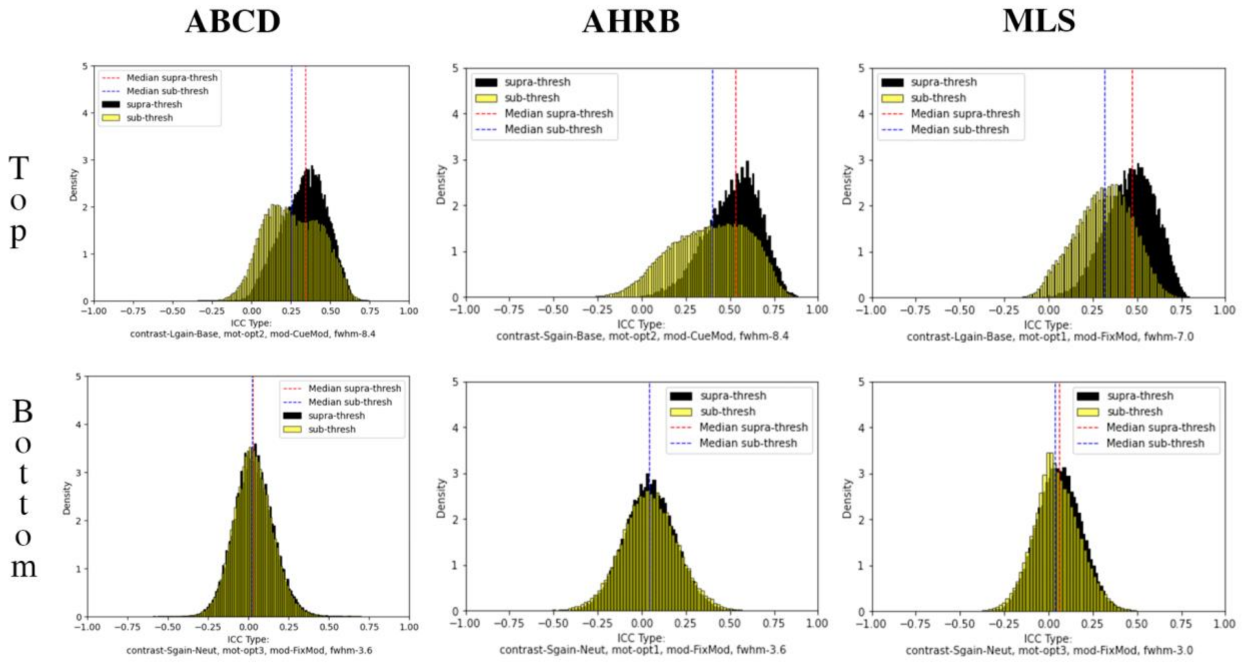


*Figure S16.* Voxelwise Distribution of ICCs for Supra- and Sub-threshold mask for highest (Top) and Lowest (Bottom) estimates from in-text Figure 3 and Figure S13 Across ABCD, AHRB and MLS samples.

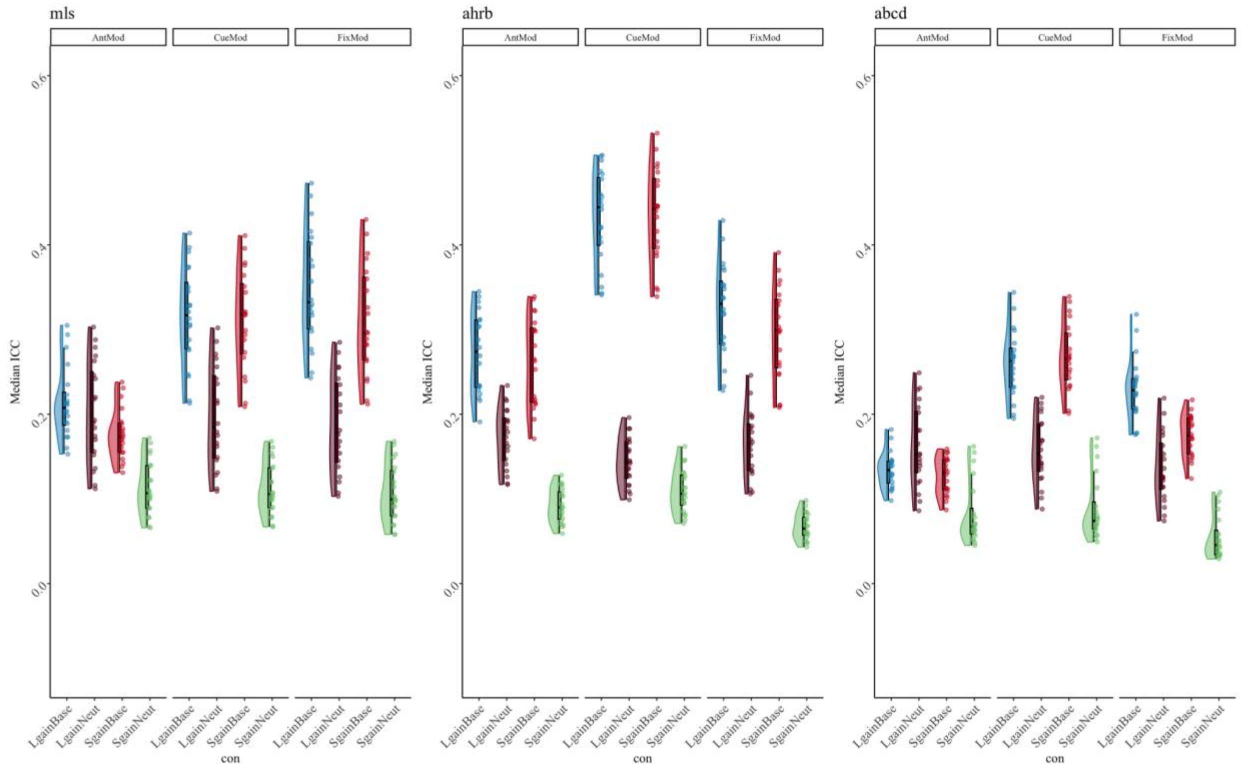


*Figure S17.* Supra-threshold Median ICC between-session reliability estimates for Contrast (con) and Model Parameterization analytic options across the ABCD, AHRB and MLS samples.

*Table S9*. Hierarchical Linear Model: (A) Linear associations between the analytic decisions and the *Between Session* median Intraclass Correlation Coefficient (ICC[3,1]), Between-subject (BS) and Within-subject variance (WS) from **supra-threshold mask** and (B) the impact of the analytic category on the marginal $R^2$.

**A. HLM Estimates for Supra-threshold Mask**

| | Median ICC | | | Median BS | | | Median WS | | |
|---|---|---|---|---|---|---|---|---|---|
| *Predictors* | *b* | *CI* | *p* | *b* | *CI* | *p* | *b* | *CI* | *p* |
| (Intercept) | .22 | .18 – .26 | <.001 | .15 | .11 – .20 | <.001 | .49 | .39 – .60 | <.001 |
| Reference [3.6] | | | | | | | | | |
| fwhm [4.8] | .03 | .01 – .04 | <.001 | -.01 | -.03 – .00 | .11 | -.11 | -.14 – -.09 | <.001 |
| fwhm [6.0] | .05 | .03 – .06 | <.001 | -.02 | -.04 – -.01 | .01 | -.18 | -.20 – -.15 | <.001 |
| fwhm [7.2] | .06 | .05 – .07 | <.001 | -.03 | -.05 – -.01 | <.001 | -.22 | -.24 – -.19 | <.001 |
| fwhm [8.4] | .07 | .06 – .09 | <.001 | -.04 | -.05 – -.02 | <.001 | -.25 | -.27 – -.22 | <.001 |
| Reference [opt1] | | | | | | | | | |
| motion [opt2] | .00 | -.01 – .01 | .50 | -.02 | -.03 – -.00 | .01 | -.05 | -.08 – -.03 | <.001 |
| motion [opt3] | -.03 | -.04 – -.02 | <.001 | -.06 | -.07 – -.04 | <.001 | -.12 | -.14 – -.09 | <.001 |
| motion [opt4] | -.03 | -.04 – -.02 | <.001 | -.06 | -.07 – -.04 | <.001 | -.12 | -.14 – -.10 | <.001 |
| Reference [AntMod] | | | | | | | | | |
| model [CueMod] | .07 | .06 – .08 | <.001 | .08 | .07 – .10 | <.001 | .17 | .15 – .19 | <.001 |
| model [FixMod] | .03 | .02 – .04 | <.001 | .07 | .06 – .08 | <.001 | .16 | .14 – .18 | <.001 |
| Reference [LgainBase] | | | | | | | | | |
| con [LgainNeut] | -.11 | -.12 – -.10 | <.001 | -.12 | -.13 – -.11 | <.001 | -.23 | -.25 – -.20 | <.001 |
| con [SgainBase] | -.02 | -.03 – -.01 | .00 | -.02 | -.03 – -.00 | .03 | -.01 | -.03 – .02 | .52 |
| con [SgainNeut] | -.19 | -.20 – -.18 | <.001 | -.14 | -.15 – -.12 | <.001 | -.24 | -.26 – -.22 | <.001 |

**B. Analytic Category Model Impact**

| Comparison | $\chi2$ | Orig R2 | New R2 | ΔR2 | $\chi2$ | Orig R2 | New R2 | ΔR2 | $\chi2$ | Orig R2 | New R2 | ΔR2 |
|---|---|---|---|---|---|---|---|---|---|---|---|---|
| [Full] vs [New - fwhm] | 159 | .66 | .60 | .06 | 25 | .49 | .48 | .01 | 336 | .59 | .43 | .16 |
| [Full] vs [New - motion] | 65 | .66 | .64 | .02 | 94 | .49 | .44 | .05 | 126 | .59 | .54 | .05 |
| [Full] vs [New - model] | 174 | .66 | .59 | .07 | 185 | .49 | .38 | .11 | 275 | .59 | .47 | .12 |

| | | | | | | | | | | | | |
|---|---|---|---|---|---|---|---|---|---|---|---|---|
| [Full] vs [New - con] | 800 | .66 | .15 | .51 | 421 | .49 | .18 | .31 | 507 | .59 | .32 | .27 |

*Table S10*: Tukey's HSB Estimate Means Differences for Supra-threshold Between-session ICC Model Parameters in Table S9.

| Contrast | Est | SE | Low.CI | Up.CI | *p* |
|---|---|---|---|---|---|
| fwhm3.6 - fwhm4.8 | -.03 | .01 | -.04 | -.01 | .001 |
| fwhm3.6 - fwhm6.0 | -.05 | .01 | -.06 | -.03 | .000 |
| fwhm3.6 - fwhm7.2 | -.06 | .01 | -.08 | -.05 | .000 |
| fwhm3.6 - fwhm8.4 | -.07 | .01 | -.09 | -.06 | .000 |
| fwhm4.8 - fwhm6.0 | -.02 | .01 | -.04 | .00 | .009 |
| fwhm4.8 - fwhm7.2 | -.04 | .01 | -.05 | -.02 | .000 |
| fwhm4.8 - fwhm8.4 | -.05 | .01 | -.07 | -.03 | .000 |
| fwhm6.0 - fwhm7.2 | -.02 | .01 | -.03 | .00 | .089 |
| fwhm6.0 - fwhm8.4 | -.03 | .01 | -.04 | -.01 | .000 |
| fwhm7.2 - fwhm8.4 | -.01 | .01 | -.03 | .01 | .342 |
| LgainBase - LgainNeut | .11 | .01 | .10 | .13 | .000 |
| LgainBase - SgainBase | .02 | .01 | .00 | .03 | .005 |
| LgainBase - SgainNeut | .19 | .01 | .18 | .20 | .000 |
| LgainNeut - SgainBase | -.09 | .01 | -.11 | -.08 | .000 |
| LgainNeut - SgainNeut | .08 | .01 | .06 | .09 | .000 |
| SgainBase - SgainNeut | .17 | .01 | .16 | .19 | .000 |
| opt1 - opt2 | .00 | .01 | -.02 | .01 | .906 |
| opt1 - opt3 | .03 | .01 | .01 | .04 | .000 |
| opt1 - opt4 | .03 | .01 | .02 | .05 | .000 |
| opt2 - opt3 | .03 | .01 | .02 | .05 | .000 |
| opt2 - opt4 | .04 | .01 | .02 | .05 | .000 |
| opt3 - opt4 | .00 | .01 | -.01 | .02 | .922 |
| AntMod - CueMod | -.07 | .00 | -.08 | -.06 | .000 |
| AntMod - FixMod | -.03 | .00 | -.04 | -.02 | .000 |
| CueMod - FixMod | .04 | .00 | .02 | .05 | .000 |

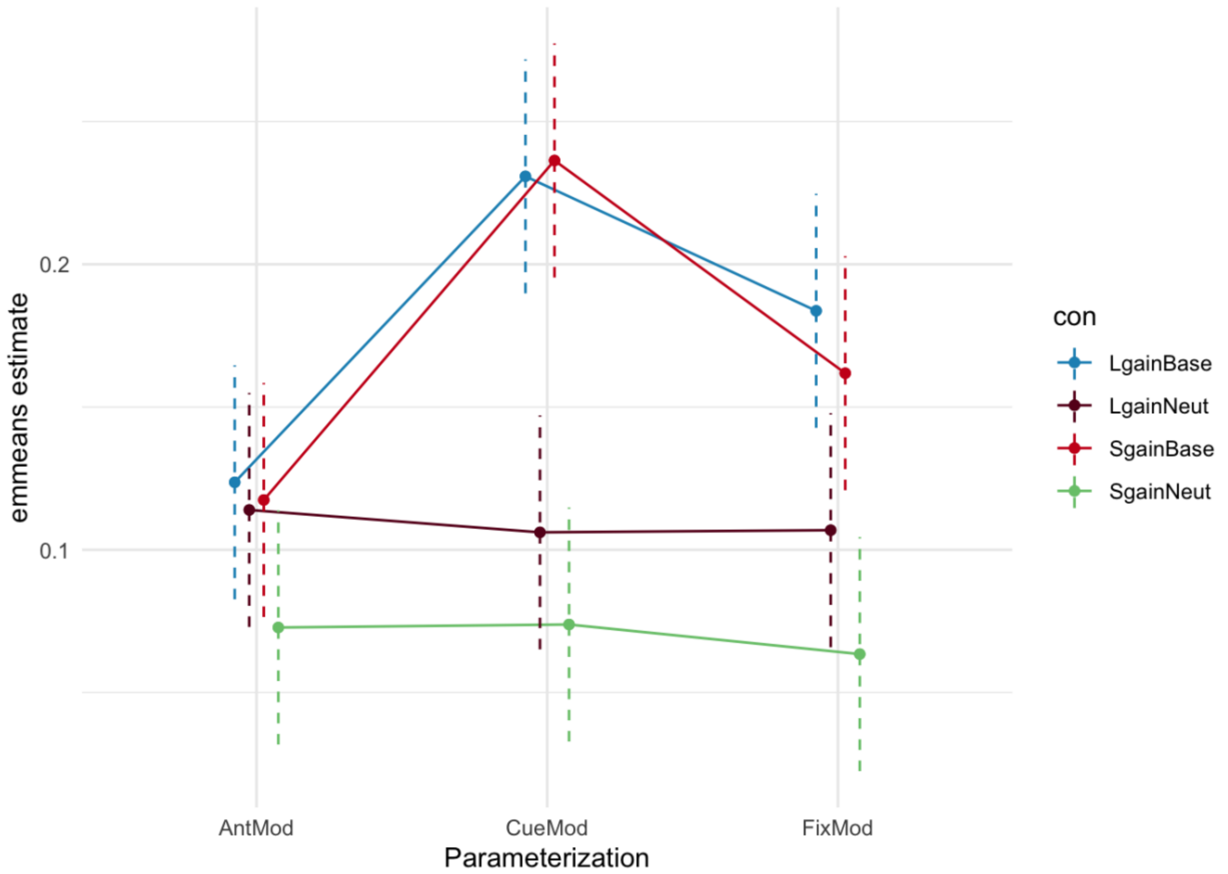


*Figure S18*: Interaction plot of *emmeans* fitted model of Contrast-by-Model parameterization for Between-session supra-threshold median ICC estimates using *emmip()*. Point estimate is a linear estimate from *emmeans* function. Dashed bars are estimated confidence intervals by *emmeans*.

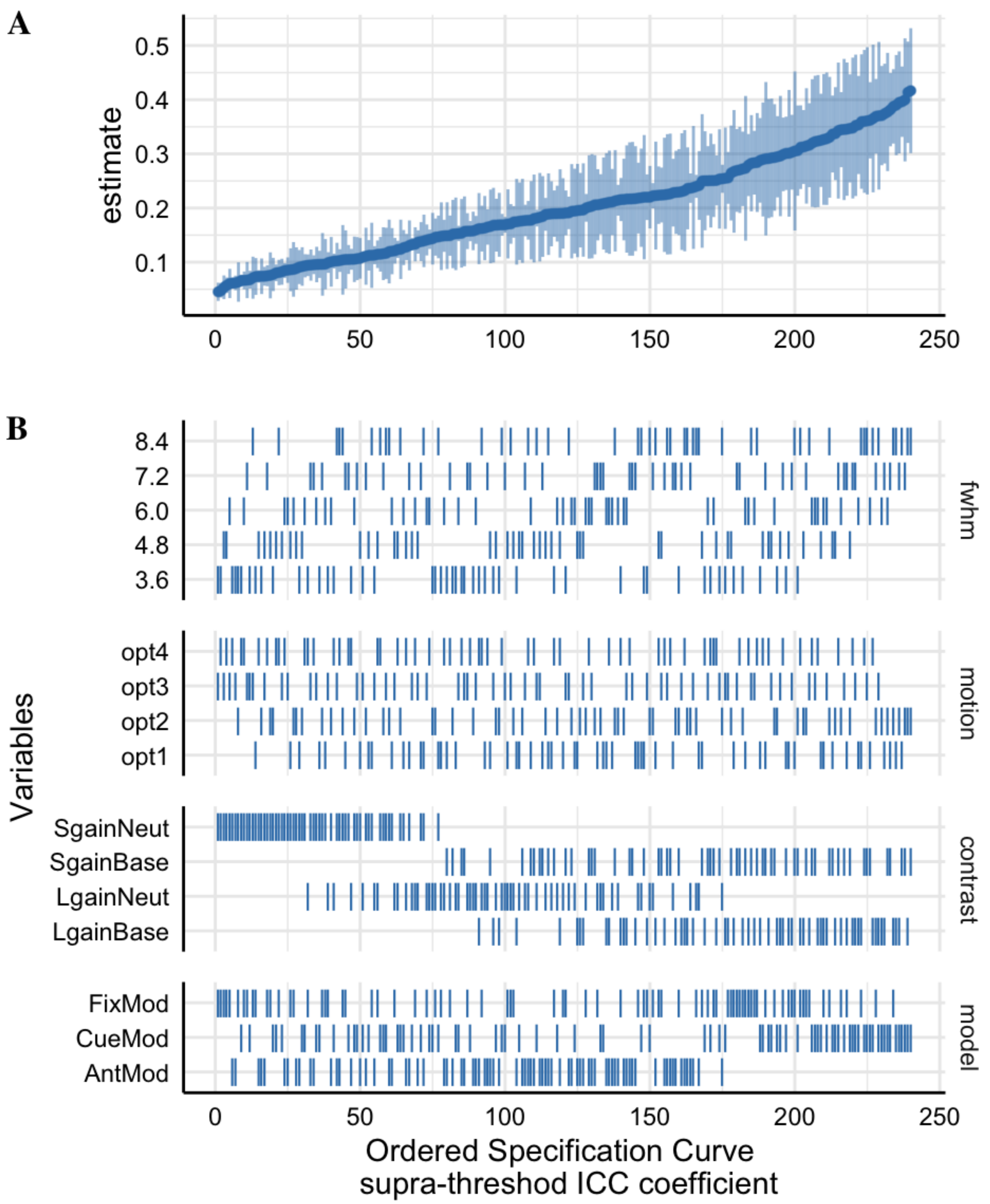


*Figure S19*: The supra-threshold Specification Curve of the Between-Session Median ICC estimates across 240 pipeline permutations for the ABCD, AHRB and MLS estimate.
A. The distribution of the point estimate (average) across the three studies and distribution across the three samples.
B. The model options (four) associated with each estimate.

*Table S11*. Hierarchical Linear Model: (A) Linear associations between the analytic decisions and the *Between Session* median Intraclass Correlation Coefficient (ICC[3,1]), Between-subject (BS) and Within-subject variance (WS) from **sub-threshold mask** and (B) the impact of the analytic category on the marginal $R^2$.

**A. HLM Estimates for Sub-threshold Mask**

| | Median ICC | | | Median BS | | | Median WS | | |
|---|---|---|---|---|---|---|---|---|---|
| *Predictors* | *b* | *CI* | *p* | *b* | *CI* | *p* | *b* | *CI* | *p* |
| (Intercept) | .13 | .10 – .16 | <.001 | .14 | .10 – .19 | <.001 | .84 | .67 – 1.02 | <.001 |
| Reference [3.6] | | | | | | | | | |
| fwhm [4.8] | .02 | .01 – .03 | <.001 | -.01 | -.02 – .01 | .24 | -.19 | -.23 – -.15 | <.001 |
| fwhm [6.0] | .04 | .03 – .05 | <.001 | -.02 | -.03 – -.00 | .04 | -.30 | -.34 – -.26 | <.001 |
| fwhm [7.2] | .05 | .04 – .06 | <.001 | -.02 | -.04 – -.01 | .01 | -.37 | -.41 – -.33 | <.001 |
| fwhm [8.4] | .07 | .06 – .08 | <.001 | -.03 | -.04 – -.01 | .00 | -.41 | -.46 – -.37 | <.001 |
| Reference [opt1] | | | | | | | | | |
| motion [opt2] | .00 | -.01 – .01 | .87 | -.02 | -.04 – -.01 | .00 | -.11 | -.15 – -.07 | <.001 |
| motion [opt3] | -.03 | -.03 – -.02 | <.001 | -.07 | -.08 – -.05 | <.001 | -.22 | -.26 – -.18 | <.001 |
| motion [opt4] | -.03 | -.03 – -.02 | <.001 | -.07 | -.08 – -.05 | <.001 | -.22 | -.26 – -.18 | <.001 |
| Reference [AntMod] | | | | | | | | | |
| model [CueMod] | .05 | .05 – .06 | <.001 | .09 | .08 – .11 | <.001 | .26 | .22 – .29 | <.001 |
| model [FixMod] | .02 | .01 – .03 | <.001 | .06 | .05 – .07 | <.001 | .25 | .21 – .28 | <.001 |
| Reference [LgainBase] | | | | | | | | | |
| con [LgainNeut] | -.07 | -.08 – -.06 | <.001 | -.12 | -.13 – -.10 | <.001 | -.37 | -.41 – -.33 | <.001 |
| con [SgainBase] | -.01 | -.02 – .00 | .07 | -.01 | -.02 – .00 | .11 | -.02 | -.06 – .02 | .41 |
| con [SgainNeut] | -.11 | -.12 – -.10 | <.001 | -.13 | -.14 – -.12 | <.001 | -.39 | -.43 – -.35 | <.001 |

**B. Analytic Category Model Impact**

| **Comparison** | χ2 | Orig R2 | New R2 | ΔR2 | χ2 | Orig R2 | New R2 | ΔR2 | χ2 | Orig R2 | New R2 | ΔR2 |
|---|---|---|---|---|---|---|---|---|---|---|---|---|
| [Full] vs [New - fwhm] | 225 | .62 | .51 | .11 | 14 | .51 | .50 | .01 | 343 | .58 | .42 | .16 |
| [Full] vs [New - motion] | 79 | .62 | .59 | .03 | 122 | .51 | .44 | .07 | 153 | .58 | .52 | .06 |
| [Full] vs [New - model] | 205 | .62 | .52 | .10 | 216 | .51 | .38 | .13 | 236 | .58 | .48 | .10 |

| | | | | | | | | | | | | |
|---|---|---|---|---|---|---|---|---|---|---|---|---|
| [Full] vs [New - con] | 609 | .62 | .24 | .38 | 424 | .51 | .21 | .30 | 484 | .58 | .33 | .25 |

*Table S12*: Tukey's HSB Estimate Mean Differences for Sub-threshold Between-session ICC Model Parameters in Table S11.

| Contrast | Est | SE | Low.CI | Up.CI | *p* |
|---|---|---|---|---|---|
| fwhm3.6 - fwhm4.8 | -.02 | .00 | -.03 | -.01 | .000 |
| fwhm3.6 - fwhm6.0 | -.04 | .00 | -.05 | -.03 | .000 |
| fwhm3.6 - fwhm7.2 | -.05 | .00 | -.07 | -.04 | .000 |
| fwhm3.6 - fwhm8.4 | -.07 | .00 | -.08 | -.05 | .000 |
| fwhm4.8 - fwhm6.0 | -.02 | .00 | -.03 | -.01 | .001 |
| fwhm4.8 - fwhm7.2 | -.03 | .00 | -.05 | -.02 | .000 |
| fwhm4.8 - fwhm8.4 | -.05 | .00 | -.06 | -.03 | .000 |
| fwhm6.0 - fwhm7.2 | -.02 | .00 | -.03 | .00 | .008 |
| fwhm6.0 - fwhm8.4 | -.03 | .00 | -.04 | -.02 | .000 |
| fwhm7.2 - fwhm8.4 | -.01 | .00 | -.03 | .00 | .042 |
| LgainBase - LgainNeut | .07 | .00 | .06 | .08 | .000 |
| LgainBase - SgainBase | .01 | .00 | .00 | .02 | .274 |
| LgainBase - SgainNeut | .11 | .00 | .10 | .12 | .000 |
| LgainNeut - SgainBase | -.06 | .00 | -.07 | -.05 | .000 |
| LgainNeut - SgainNeut | .04 | .00 | .03 | .05 | .000 |
| SgainBase - SgainNeut | .10 | .00 | .09 | .11 | .000 |
| opt1 - opt2 | .00 | .00 | -.01 | .01 | .999 |
| opt1 - opt3 | .03 | .00 | .02 | .04 | .000 |
| opt1 - opt4 | .03 | .00 | .02 | .04 | .000 |
| opt2 - opt3 | .03 | .00 | .02 | .04 | .000 |
| opt2 - opt4 | .03 | .00 | .02 | .04 | .000 |
| opt3 - opt4 | .00 | .00 | -.01 | .01 | 1.000 |
| AntMod - CueMod | -.05 | .00 | -.06 | -.05 | .000 |
| AntMod - FixMod | -.02 | .00 | -.03 | -.01 | .000 |
| CueMod - FixMod | .03 | .00 | .02 | .04 | .000 |

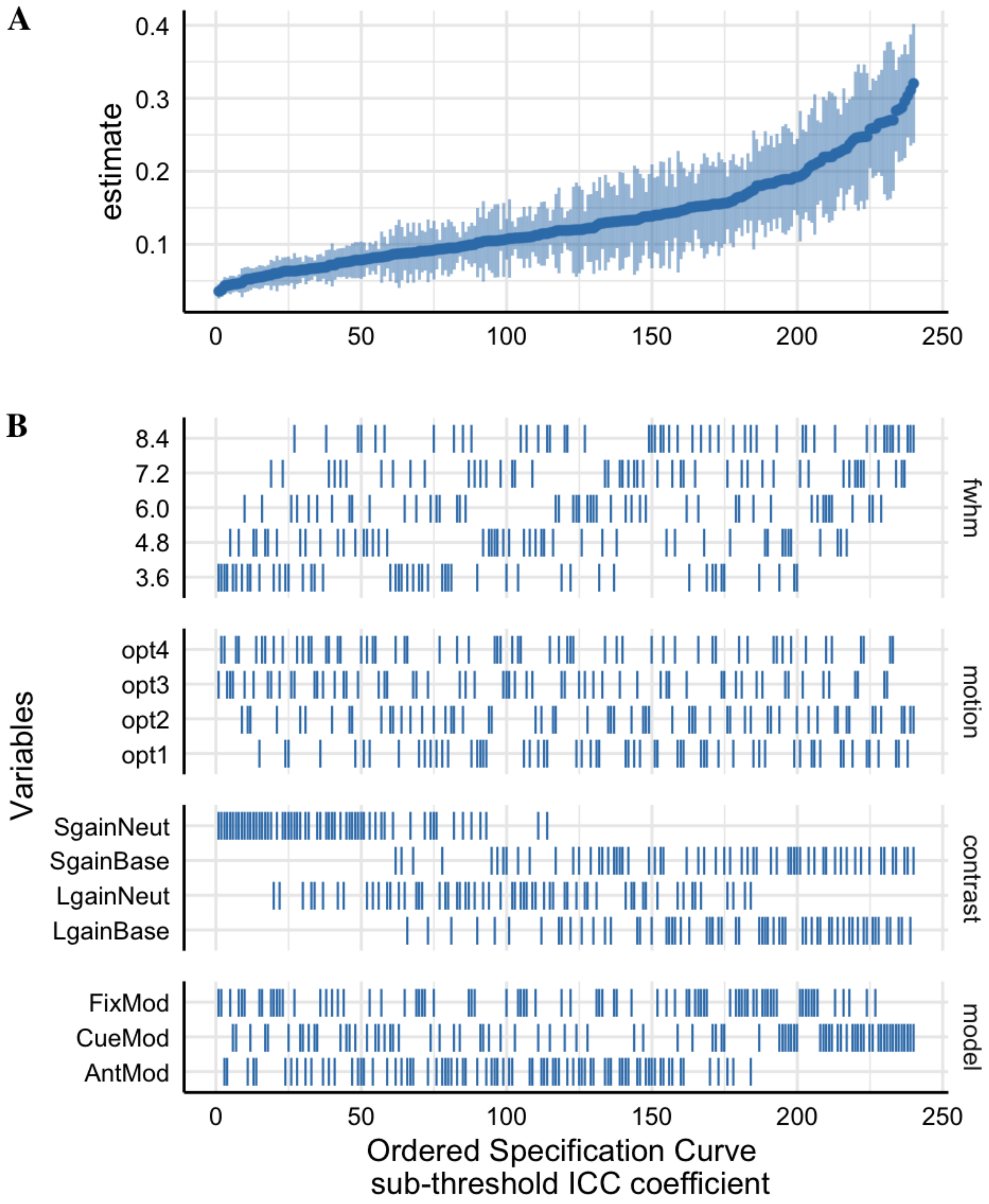


*Figure S20*: The sub-threshold Specification Curve of the Between-Session Median ICC estimates across 240 pipeline permutations for the ABCD, AHRB and MLS estimate.
A. The distribution of the point estimate (average) across the three studies and distribution across the three samples.
B. The model options (four) associated with each estimate.

*B. Between-Run Group Reliability:*

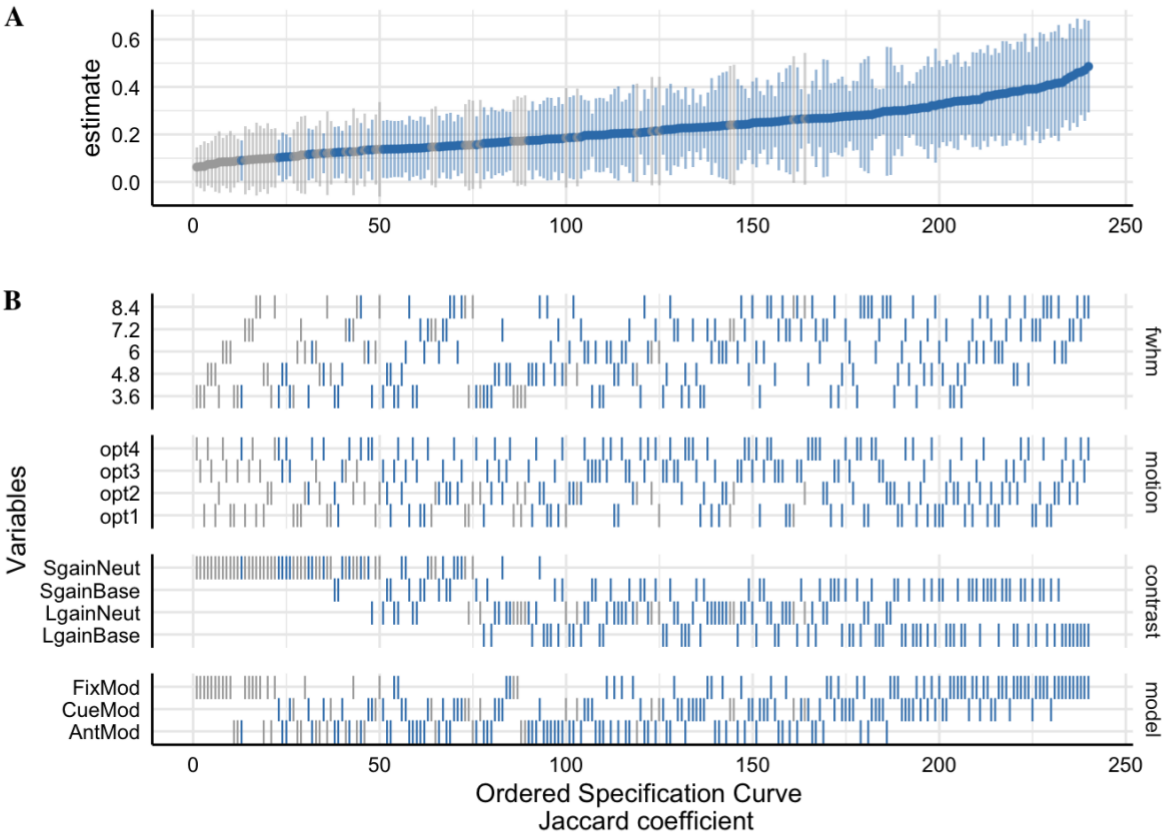


*Figure S21*: The Specification Curve of the Session 1 Between-run Jaccard Similarity estimates across 240 pipeline permutations for the ABCD, AHRB and MLS samples.
A. The distribution of the point estimate (average) across the three studies and distribution across the three samples.
B. The model options (four) associated with each estimate.

*Table S13*: Tukey's HSB Estimate Means Differences for (A) Jaccard and (B) Spearman Model Parameters in-text Table 4.

| Contrast | Est | SE | Low.CI | Up.CI | *p* |
|---|---|---|---|---|---|
| A. Jaccard Similarity | | | | | |
| fwhm3.6 - fwhm4.8 | -.03 | .01 | -.06 | .00 | .037 |
| fwhm3.6 - fwhm6 | -.05 | .01 | -.08 | -.02 | .000 |
| fwhm3.6 - fwhm7.2 | -.07 | .01 | -.10 | -.04 | .000 |
| fwhm3.6 - fwhm8.4 | -.08 | .01 | -.11 | -.06 | .000 |
| fwhm4.8 - fwhm6 | -.02 | .01 | -.05 | .01 | .171 |
| fwhm4.8 - fwhm7.2 | -.04 | .01 | -.07 | -.01 | .001 |
| fwhm4.8 - fwhm8.4 | -.05 | .01 | -.08 | -.03 | .000 |
| fwhm6 - fwhm7.2 | -.02 | .01 | -.05 | .01 | .448 |
| fwhm6 - fwhm8.4 | -.03 | .01 | -.06 | .00 | .031 |
| fwhm7.2 - fwhm8.4 | -.01 | .01 | -.04 | .02 | .737 |
| LgainBase - LgainNeut | .09 | .01 | .06 | .11 | .000 |
| LgainBase - SgainBase | .03 | .01 | .01 | .05 | .008 |
| LgainBase - SgainNeut | .18 | .01 | .16 | .21 | .000 |
| LgainNeut - SgainBase | -.05 | .01 | -.08 | -.03 | .000 |
| LgainNeut - SgainNeut | .10 | .01 | .07 | .12 | .000 |
| SgainBase - SgainNeut | .15 | .01 | .13 | .18 | .000 |
| opt1 - opt2 | -.01 | .01 | -.04 | .01 | .437 |
| opt1 - opt3 | .00 | .01 | -.02 | .03 | .998 |
| opt1 - opt4 | .00 | .01 | -.03 | .02 | .979 |
| opt2 - opt3 | .02 | .01 | -.01 | .04 | .332 |
| opt2 - opt4 | .01 | .01 | -.01 | .03 | .687 |
| opt3 - opt4 | -.01 | .01 | -.03 | .02 | .938 |
| AntMod - CueMod | -.05 | .01 | -.07 | -.03 | .000 |
| AntMod - FixMod | -.08 | .01 | -.10 | -.07 | .000 |
| CueMod - FixMod | -.03 | .01 | -.05 | -.01 | .000 |
| B. Spearman Supra-threshold Similarity | | | | | |
| fwhm3.6 - fwhm4.8 | -.05 | .01 | -.07 | -.03 | .000 |
| fwhm3.6 - fwhm6 | -.09 | .01 | -.11 | -.07 | .000 |
| fwhm3.6 - fwhm7.2 | -.11 | .01 | -.14 | -.09 | .000 |
| fwhm3.6 - fwhm8.4 | -.13 | .01 | -.16 | -.11 | .000 |
| fwhm4.8 - fwhm6 | -.04 | .01 | -.06 | -.02 | .000 |
| fwhm4.8 - fwhm7.2 | -.06 | .01 | -.09 | -.04 | .000 |

| | | | | | |
|---|---|---|---|---|---|
| fwhm4.8 - fwhm8.4 | -.08 | .01 | -.11 | -.06 | .000 |
| fwhm6 - fwhm7.2 | -.03 | .01 | -.05 | .00 | .008 |
| fwhm6 - fwhm8.4 | -.05 | .01 | -.07 | -.02 | .000 |
| fwhm7.2 - fwhm8.4 | -.02 | .01 | -.04 | .00 | .107 |
| LgainBase - LgainNeut | .20 | .01 | .18 | .22 | .000 |
| LgainBase - SgainBase | .01 | .01 | -.01 | .03 | .531 |
| LgainBase - SgainNeut | .34 | .01 | .32 | .36 | .000 |
| LgainNeut - SgainBase | -.19 | .01 | -.21 | -.17 | .000 |
| LgainNeut - SgainNeut | .14 | .01 | .12 | .16 | .000 |
| SgainBase - SgainNeut | .33 | .01 | .31 | .35 | .000 |
| opt1 - opt2 | -.01 | .01 | -.03 | .00 | .217 |
| opt1 - opt3 | -.01 | .01 | -.03 | .01 | .578 |
| opt1 - opt4 | -.01 | .01 | -.03 | .01 | .305 |
| opt2 - opt3 | .00 | .01 | -.01 | .02 | .915 |
| opt2 - opt4 | .00 | .01 | -.02 | .02 | .998 |
| opt3 - opt4 | .00 | .01 | -.02 | .02 | .967 |
| AntMod - CueMod | -.02 | .01 | -.04 | -.01 | .001 |
| AntMod - FixMod | -.01 | .01 | -.02 | .01 | .384 |
| CueMod - FixMod | .01 | .01 | .00 | .03 | .054 |

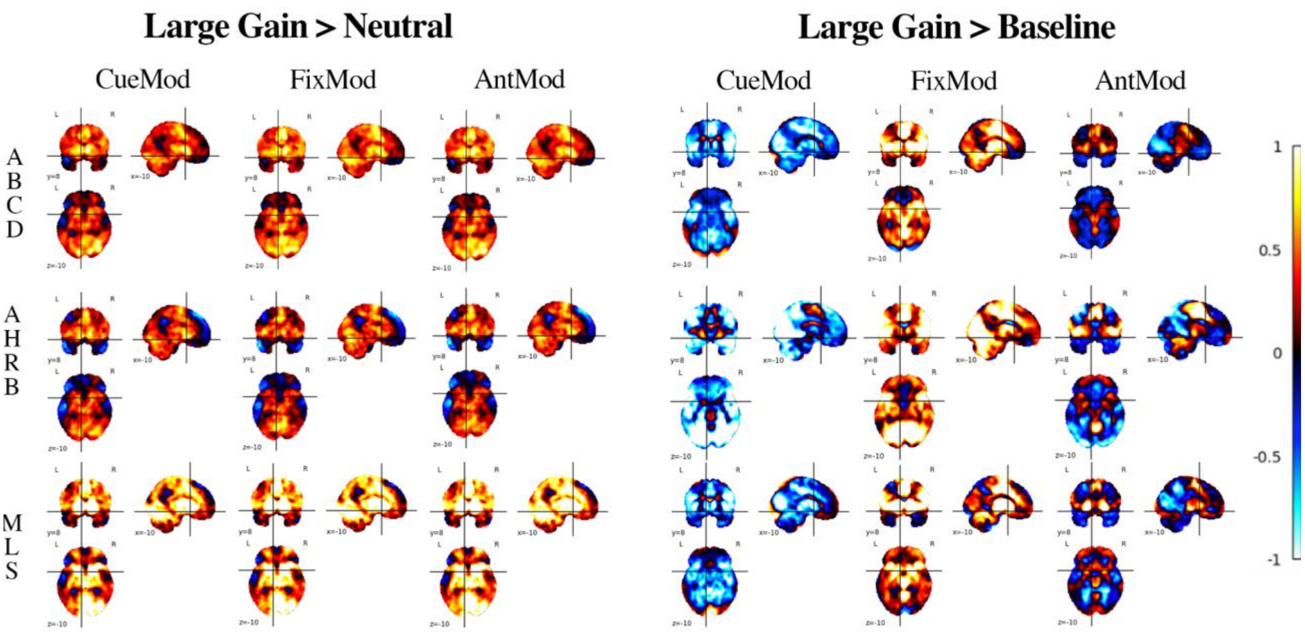


*Figure S22*: Comparing Lgain-Neut & Lgain-Base contrasts for Session 1 run average group activity for Cue, Fixation and Anticipation Parameterization for Motion opt2 and FWHM 8.4 (MLS 7.0) across ABCD, AHRB and MLS samples.

Note: For quick access on NeuroVault, example image search: "*_type-session_contrast-Lgain-Base_mask-mni152_mot-opt2_mod-CueMod_fwhm-8.4_stat-cohensd.nii.gz*"

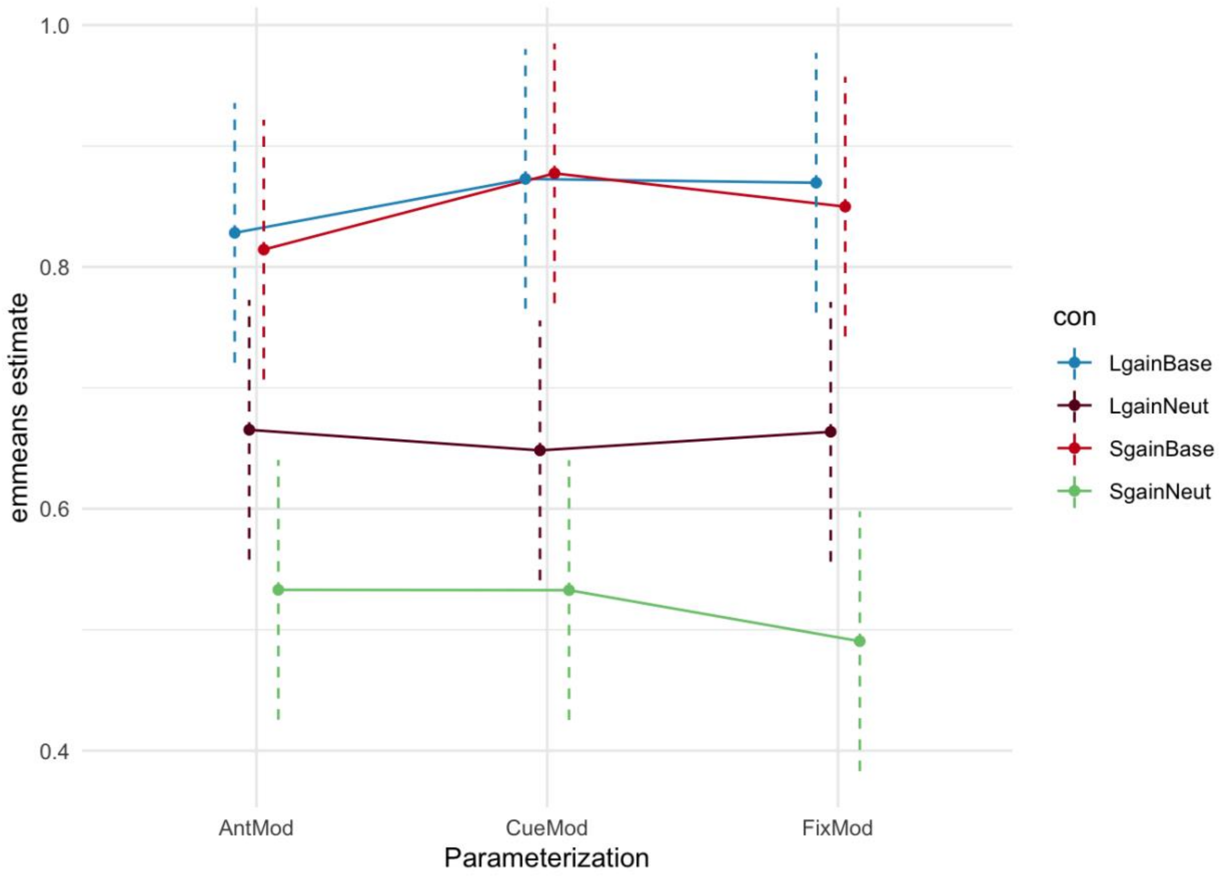


*Figure S23*: Spearman *rho*: Interaction plot of *emmeans* fitted model of Contrast-by-Model parameterization for Between-run supra-threshold Spearman Similarity estimates using *emmip()*. Point estimate is a linear spearman *rho* estimate from *emmeans* function. Dashed bars are estimated confidence intervals by *emmeans*.

*B. Between-Session Group Reliability:*

*Table S14.* Hierarchical Linear Model: (A) Linear associations between the analytic decisions and the Jaccard and Spearman supra-threshold mask between-session similarity and (B) the impact of the analytic category on the marginal $R^2$.

**A. HLM Group-map Estimates**

| | Jaccard | | | Spearman | | |
|---|---|---|---|---|---|---|
| *Predictors* | *b* | *CI* | *p* | *b* | *CI* | *p* |
| (Intercept) | .29 | .20 – .38 | <.001 | .82 | .76 – .87 | <.001 |
| Reference [3.6] | | | | | | |
| fwhm [4.8] | .04 | .02 – .06 | <.001 | .04 | .03 – .06 | <.001 |
| fwhm [6.0] | .07 | .05 – .10 | <.001 | .07 | .05 – .08 | <.001 |
| fwhm [7.2] | .10 | .08 – .12 | <.001 | .09 | .07 – .10 | <.001 |
| fwhm [8.4] | .12 | .10 – .14 | <.001 | .10 | .08 – .12 | <.001 |
| Reference [opt1] | | | | | | |
| motion [opt2] | .04 | .02 – .06 | <.001 | .03 | .02 – .04 | <.001 |
| motion [opt3] | .03 | .01 – .05 | .00 | .05 | .03 – .06 | <.001 |
| motion [opt4] | .04 | .02 – .06 | <.001 | .05 | .04 – .06 | <.001 |
| Reference [AntMod] | | | | | | |
| model [CueMod] | .00 | -.01 – .02 | .64 | -.01 | -.02 – .00 | .12 |
| model [FixMod] | .10 | .08 – .12 | <.001 | -.01 | -.02 – .01 | .31 |
| Reference [LgainBase] | | | | | | |
| con [LgainNeut] | -.06 | -.08 – -.04 | <.001 | -.15 | -.16 – -.14 | <.001 |
| con [SgainBase] | -.04 | -.06 – -.02 | <.001 | -.01 | -.03 – -.00 | .05 |
| con [SgainNeut] | -.24 | -.26 – -.22 | <.001 | -.32 | -.34 – -.31 | <.001 |

**B. Analytic Category Model Impact**

| **Comparison** | $\chi 2$ | Orig R2 | New R2 | ΔR2 | $\chi 2$ | Orig R2 | New R2 | ΔR2 |
|---|---|---|---|---|---|---|---|---|
| [Full] vs [New - fwhm] | 124 | .47 | .40 | .07 | 184 | .74 | .69 | .05 |
| [Full] vs [New - motion] | 22 | .47 | .45 | .02 | 61 | .74 | .73 | .01 |
| [Full] vs [New - model] | 149 | .47 | .39 | .08 | 3 | .74 | .74 | .00 |
| [Full] vs [New - con] | 468 | .47 | .15 | .32 | 1141 | .74 | .07 | .67 |

*Table S15*: Tukey's HSB Estimate Means Differences for (A) Jaccard and (B) Spearman Model Parameters in-text Table S14.

| Contrast | Est | SE | Low.CI | Up.CI | *p* |
|---|---|---|---|---|---|
| A. Jaccard Similarity | | | | | |
| fwhm3.6 - fwhm4.8 | -.04 | .01 | -.07 | -.01 | .003 |
| fwhm3.6 - fwhm6 | -.07 | .01 | -.11 | -.04 | .000 |
| fwhm3.6 - fwhm7.2 | -.10 | .01 | -.13 | -.07 | .000 |
| fwhm3.6 - fwhm8.4 | -.12 | .01 | -.15 | -.09 | .000 |
| fwhm4.8 - fwhm6 | -.03 | .01 | -.06 | .00 | .040 |
| fwhm4.8 - fwhm7.2 | -.06 | .01 | -.09 | -.03 | .000 |
| fwhm4.8 - fwhm8.4 | -.08 | .01 | -.11 | -.04 | .000 |
| fwhm6 - fwhm7.2 | -.02 | .01 | -.06 | .01 | .209 |
| fwhm6 - fwhm8.4 | -.04 | .01 | -.08 | -.01 | .002 |
| fwhm7.2 - fwhm8.4 | -.02 | .01 | -.05 | .01 | .455 |
| LgainBase - LgainNeut | .06 | .01 | .03 | .08 | .000 |
| LgainBase - SgainBase | .04 | .01 | .01 | .06 | .001 |
| LgainBase - SgainNeut | .24 | .01 | .21 | .27 | .000 |
| LgainNeut - SgainBase | -.02 | .01 | -.04 | .01 | .338 |
| LgainNeut - SgainNeut | .18 | .01 | .16 | .21 | .000 |
| SgainBase - SgainNeut | .20 | .01 | .18 | .23 | .000 |
| opt1 - opt2 | -.04 | .01 | -.07 | -.02 | .000 |
| opt1 - opt3 | -.03 | .01 | -.06 | .00 | .013 |
| opt1 - opt4 | -.04 | .01 | -.07 | -.01 | .001 |
| opt2 - opt3 | .01 | .01 | -.01 | .04 | .654 |
| opt2 - opt4 | .00 | .01 | -.02 | .03 | .976 |
| opt3 - opt4 | -.01 | .01 | -.03 | .02 | .880 |
| AntMod - CueMod | .00 | .01 | -.03 | .02 | .886 |
| AntMod - FixMod | -.10 | .01 | -.12 | -.08 | .000 |
| CueMod - FixMod | -.10 | .01 | -.12 | -.08 | .000 |
| B. Spearman Supra-threshold Similarity | | | | | |
| fwhm3.6 - fwhm4.8 | -.04 | .01 | -.06 | -.02 | .000 |
| fwhm3.6 - fwhm6 | -.07 | .01 | -.09 | -.05 | .000 |
| fwhm3.6 - fwhm7.2 | -.09 | .01 | -.11 | -.07 | .000 |
| fwhm3.6 - fwhm8.4 | -.10 | .01 | -.12 | -.08 | .000 |
| fwhm4.8 - fwhm6 | -.03 | .01 | -.05 | -.01 | .004 |
| fwhm4.8 - fwhm7.2 | -.05 | .01 | -.07 | -.02 | .000 |
| fwhm4.8 - fwhm8.4 | -.06 | .01 | -.08 | -.04 | .000 |

| | | | | | |
|---|---|---|---|---|---|
| fwhm6 - fwhm7.2 | -.02 | .01 | -.04 | .00 | .119 |
| fwhm6 - fwhm8.4 | -.03 | .01 | -.05 | -.01 | .001 |
| fwhm7.2 - fwhm8.4 | -.01 | .01 | -.03 | .01 | .463 |
| LgainBase - LgainNeut | .15 | .01 | .13 | .17 | .000 |
| LgainBase - SgainBase | .01 | .01 | .00 | .03 | .196 |
| LgainBase - SgainNeut | .32 | .01 | .31 | .34 | .000 |
| LgainNeut - SgainBase | -.14 | .01 | -.16 | -.12 | .000 |
| LgainNeut - SgainNeut | .17 | .01 | .15 | .19 | .000 |
| SgainBase - SgainNeut | .31 | .01 | .29 | .33 | .000 |
| opt1 - opt2 | -.03 | .01 | -.05 | -.01 | .000 |
| opt1 - opt3 | -.05 | .01 | -.06 | -.03 | .000 |
| opt1 - opt4 | -.05 | .01 | -.07 | -.03 | .000 |
| opt2 - opt3 | -.02 | .01 | -.03 | .00 | .106 |
| opt2 - opt4 | -.02 | .01 | -.04 | .00 | .024 |
| opt3 - opt4 | .00 | .01 | -.02 | .01 | .943 |
| AntMod - CueMod | .01 | .01 | .00 | .02 | .265 |
| AntMod - FixMod | .01 | .01 | -.01 | .02 | .568 |
| CueMod - FixMod | .00 | .01 | -.02 | .01 | .850 |

**2.4 Aim 2 results**

*Between-Run Reliability:*

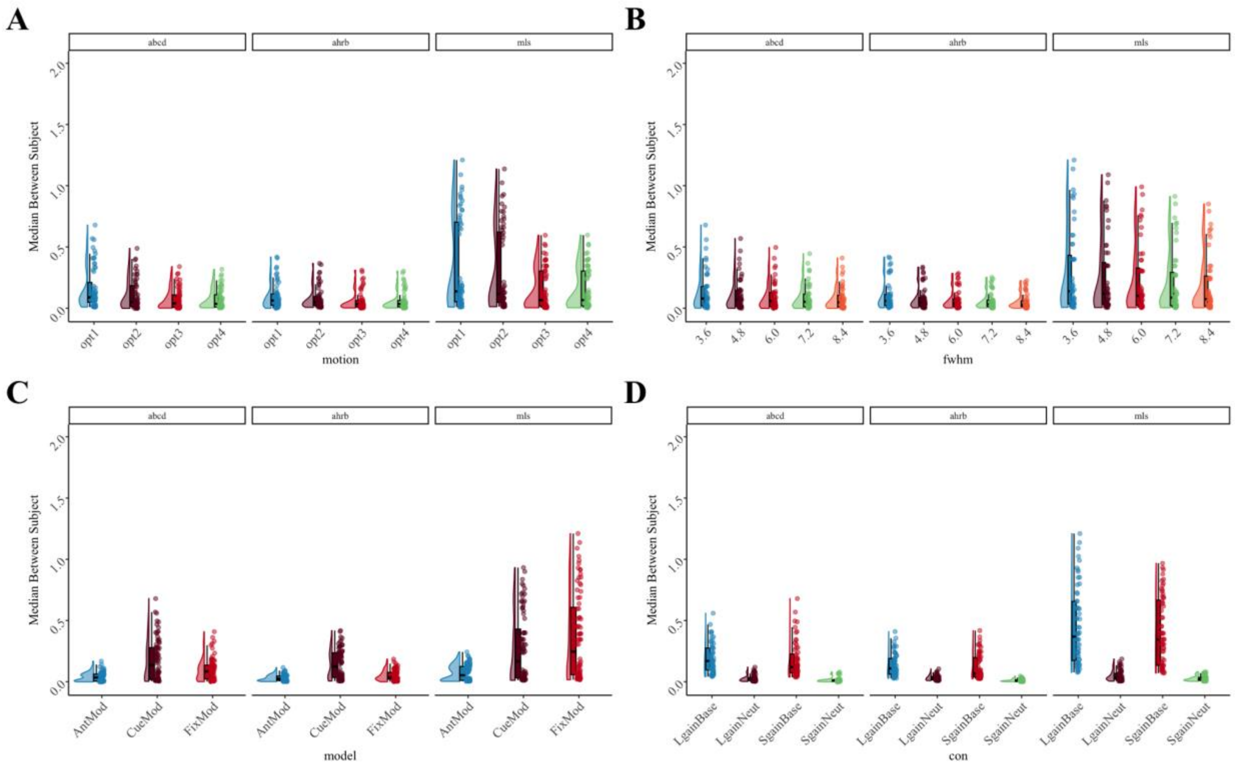


*Figure S24.* Session 1 Between-run: Supra-threshold Median **Between-subject variance** estimates across (A) Motion, (B) FWHM, (C) Model Parameterization and (D) Contrast analytic options for between-run reliability across the ABCD, AHRB and MLS samples.

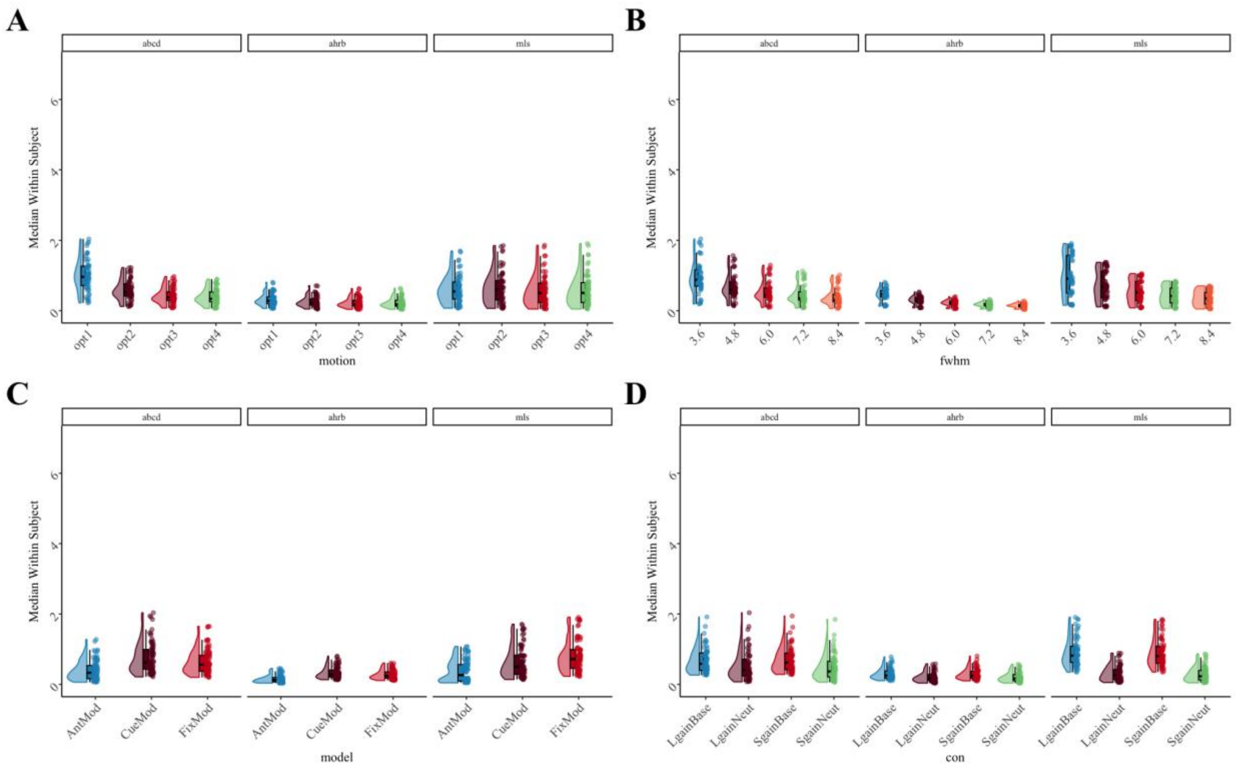


*Figure S25.* Session 1 Between-run: Supra-threshold Median **Within-subject variance** estimates across (A) Motion, (B) FWHM, (C) Model Parameterization and (D) Contrast analytic options for between-run reliability across the ABCD, AHRB and MLS samples.

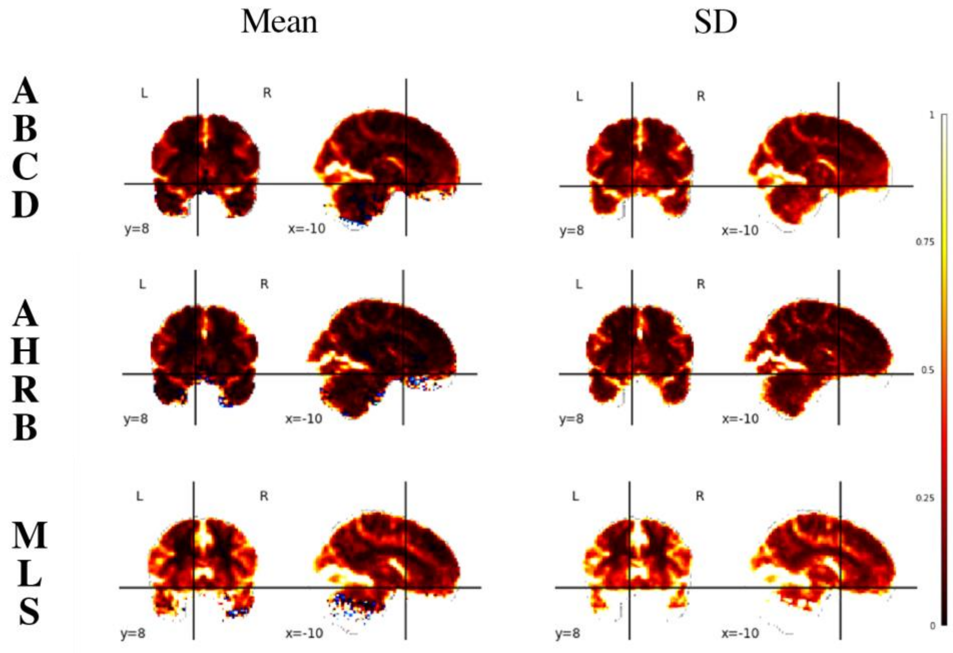


*Figure S26*: Mean and SD of Between-subject variance $(\sigma_r^2)$ estimates across 240 permutations for the Adolescent Brain Cognitive Development (ABCD), Adolescent Health Risk Behavior (AHRB) and Michigan Longitudinal (MLS) 3D volumes.

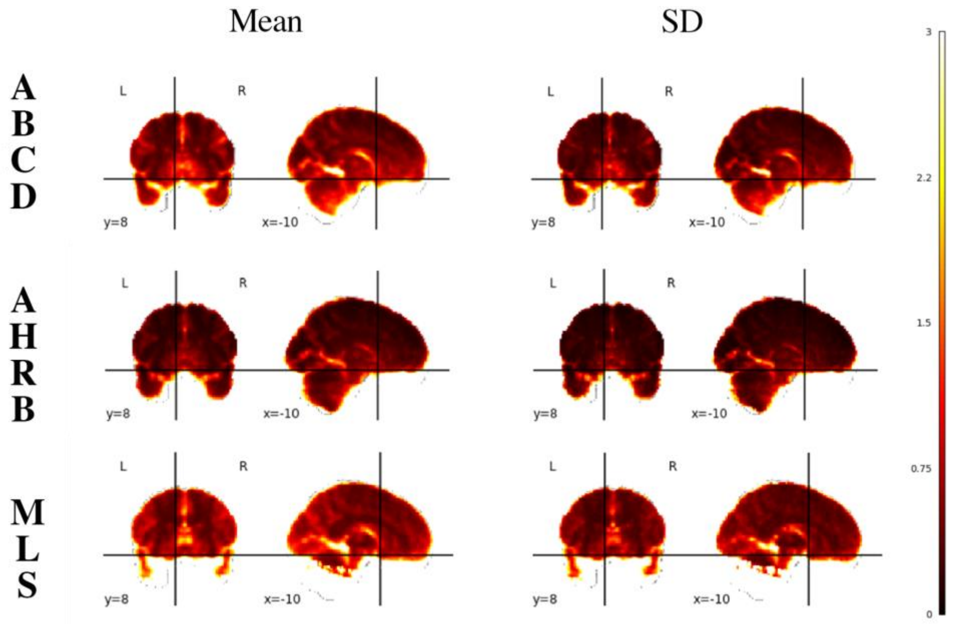


*Figure S27*: Mean and SD of Within-subject variance estimates $(\sigma_v^2)$ across 240 permutations for the Adolescent Brain Cognitive Development (ABCD), Adolescent Health Risk Behavior (AHRB) and Michigan Longitudinal (MLS) 3D volumes.

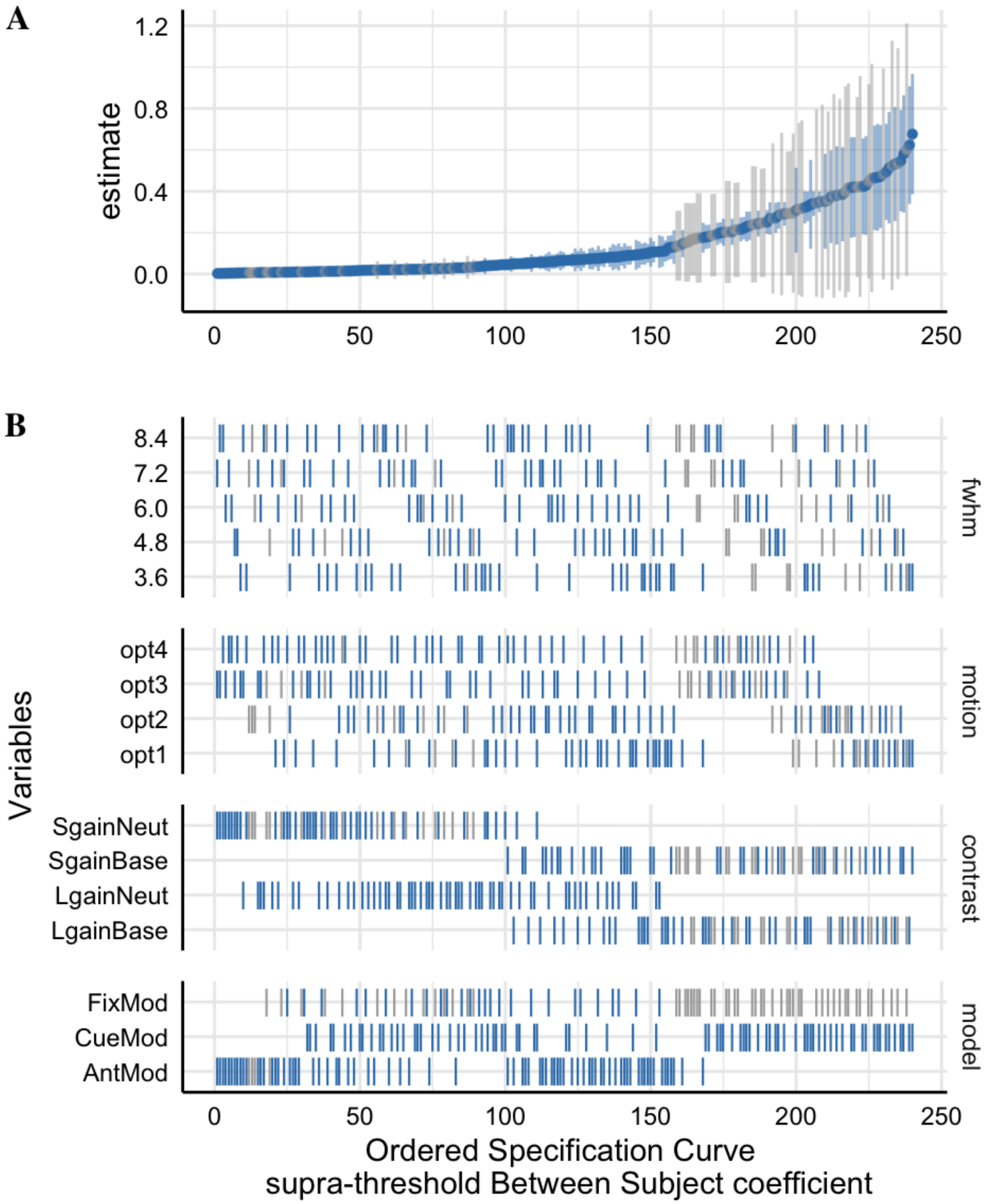


382
383 *Figure S28*: Session 1 Between-run: The supra-threshold Specification Curve of the Median
384 Between-subject variance ($\sigma_r^2$) estimates across 240 pipeline permutations for the ABCD,
385 AHRB and MLS estimate.
386 A. The distribution of the point estimate (average) across the three studies and distribution across the three samples.
387 B. The model options (four) associated with each estimate.
388

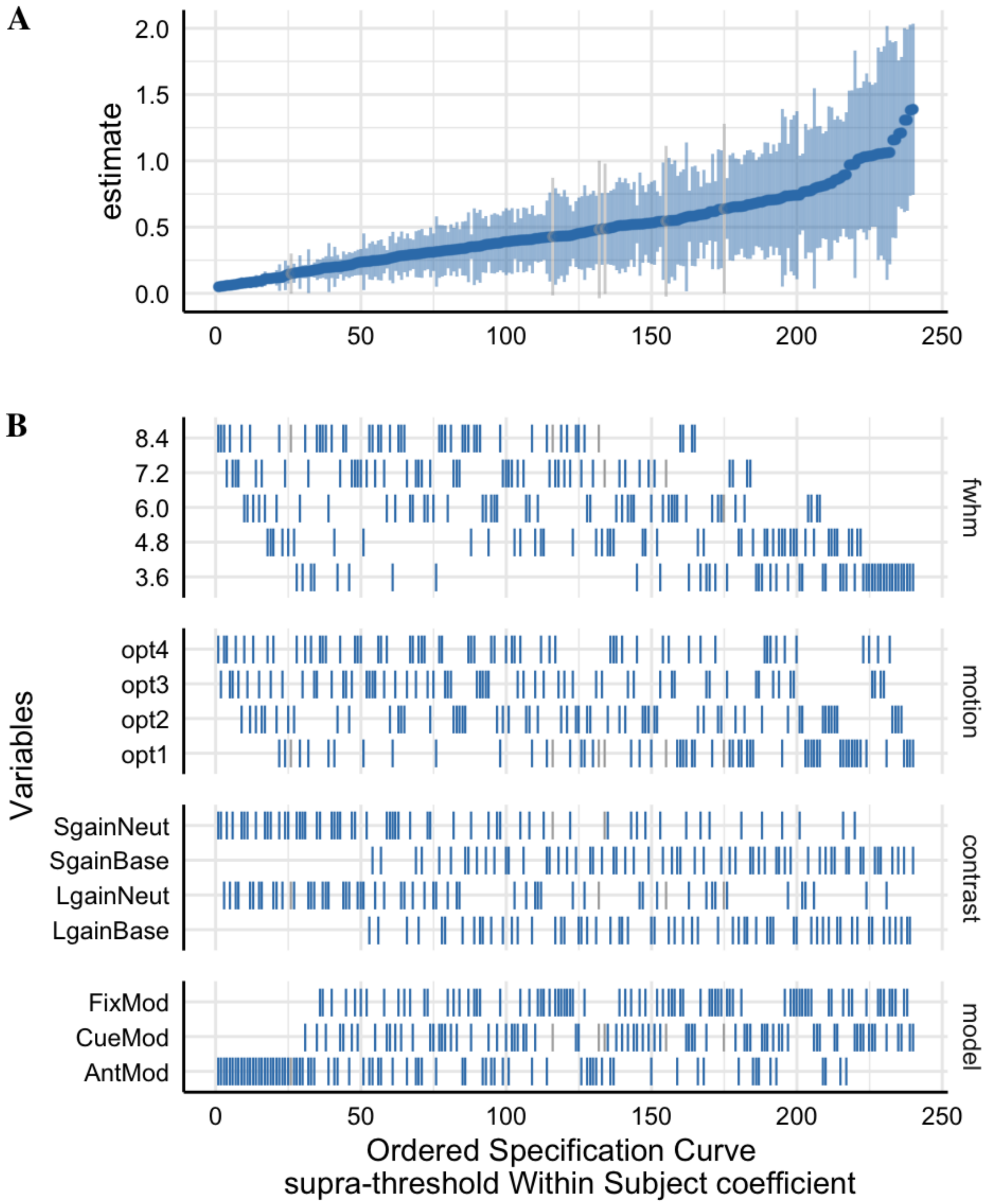


*Figure S29*: Session 1 Between-run: The supra-threshold Specification Curve of the Median Within-subject variance ($\sigma_v^2$) estimates across 240 pipeline permutations for the ABCD, AHRB and MLS estimate.

A. The distribution of the point estimate (average) across the three studies and distribution across the three samples.

B. The model options (four) associated with each estimate.

*Between-Session Reliability:*

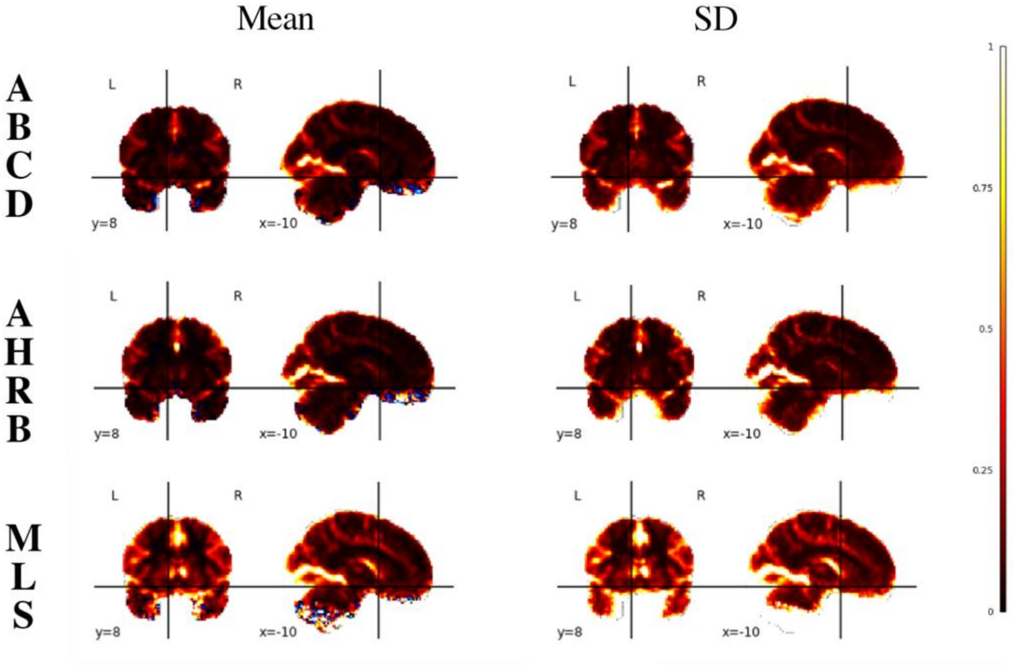


*Figure S30*: Between-session Mean and SD of Between-subject variance $(\sigma_r^2)$ estimates across 240 permutations for the Adolescent Brain Cognitive Development (ABCD), Adolescent Health Risk Behavior (AHRB) and Michigan Longitudinal (MLS) 3D volumes.

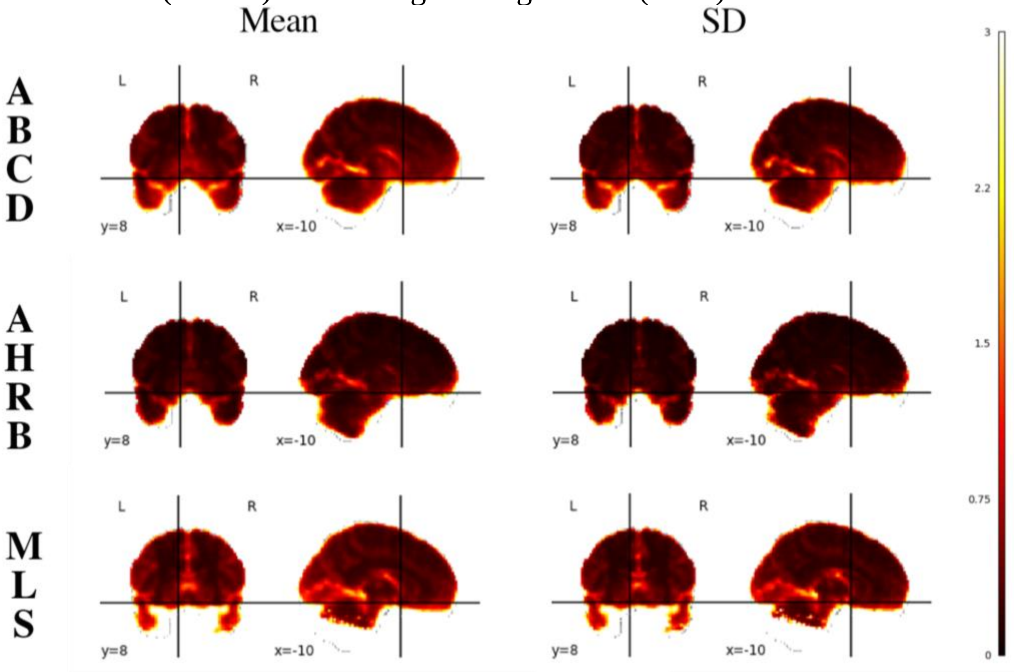


*Figure S31*: Between-session Mean and SD of Within-subject variance $(\sigma_v^2)$ estimates across 240 permutations for the Adolescent Brain Cognitive Development (ABCD), Adolescent Health Risk Behavior (AHRB) and Michigan Longitudinal (MLS) 3D volumes.

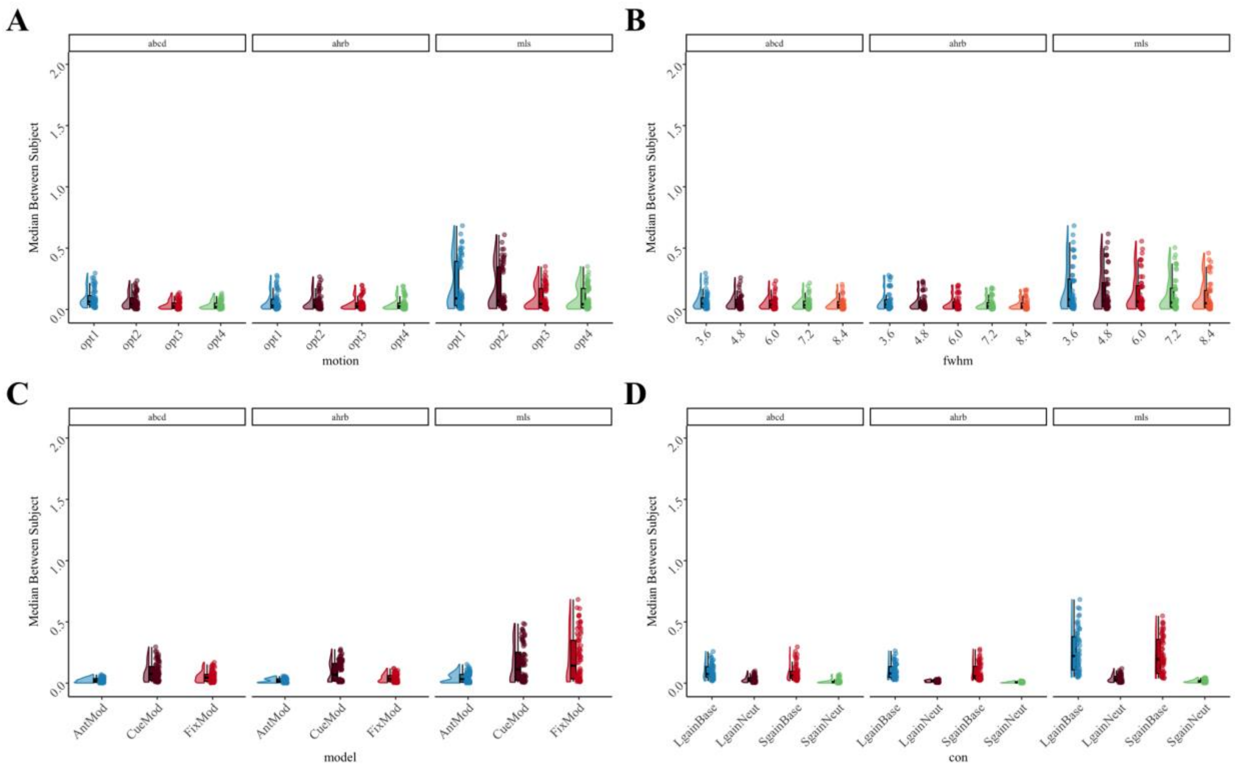


*Figure S32*. Between-session: Supra-threshold Median Between-subject variance ($\sigma_r^2$) estimates across (A) Motion, (B) FWHM, (C) Model Parameterization and (D) Contrast analytic options for between-run reliability across the ABCD, AHRB and MLS samples.

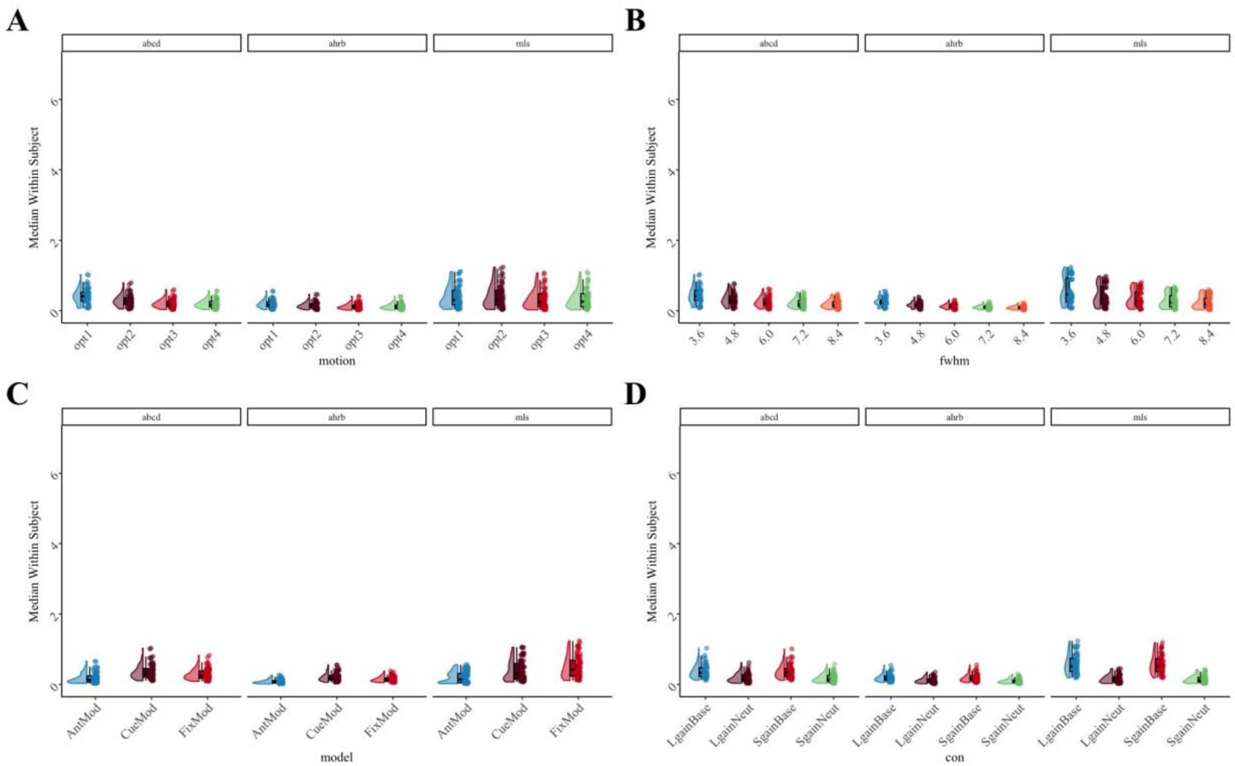


*Figure S33.* Between-session: Supra-threshold Median Within-subject variance ($\sigma_v^2$) estimates across (A) Motion, (B) FWHM, (C) Model Parameterization and (D) Contrast analytic options for between-run reliability across the ABCD, AHRB and MLS samples.

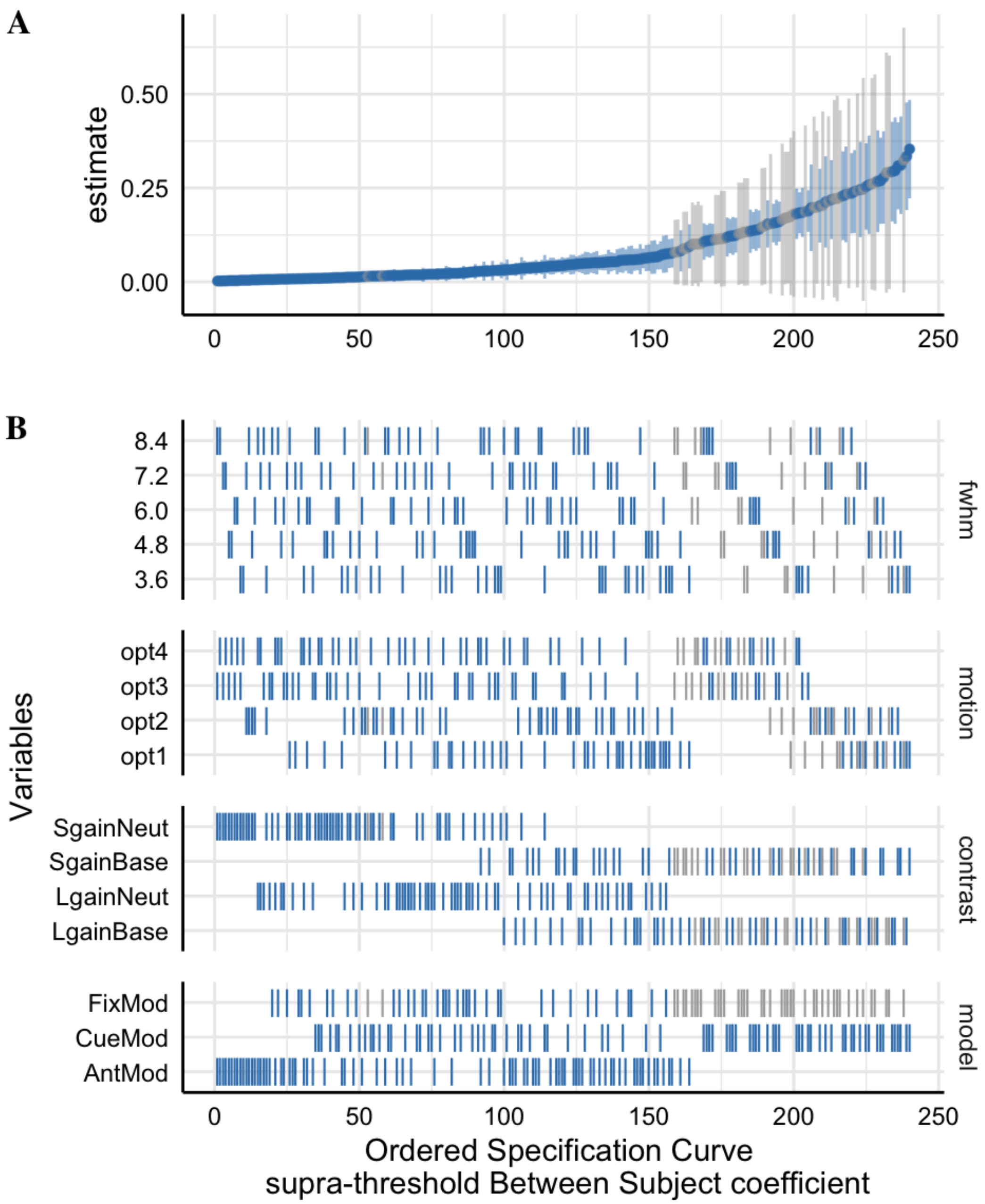


*Figure S34*: Between-session: The supra-threshold Specification Curve of the Median Between-subject variance ($\sigma_r^2$) estimates across 240 pipeline permutations for the ABCD, AHRB and MLS estimate.
A. The distribution of the point estimate (average) across the three studies and distribution across the three samples.
B. The model options (four) associated with each estimate.

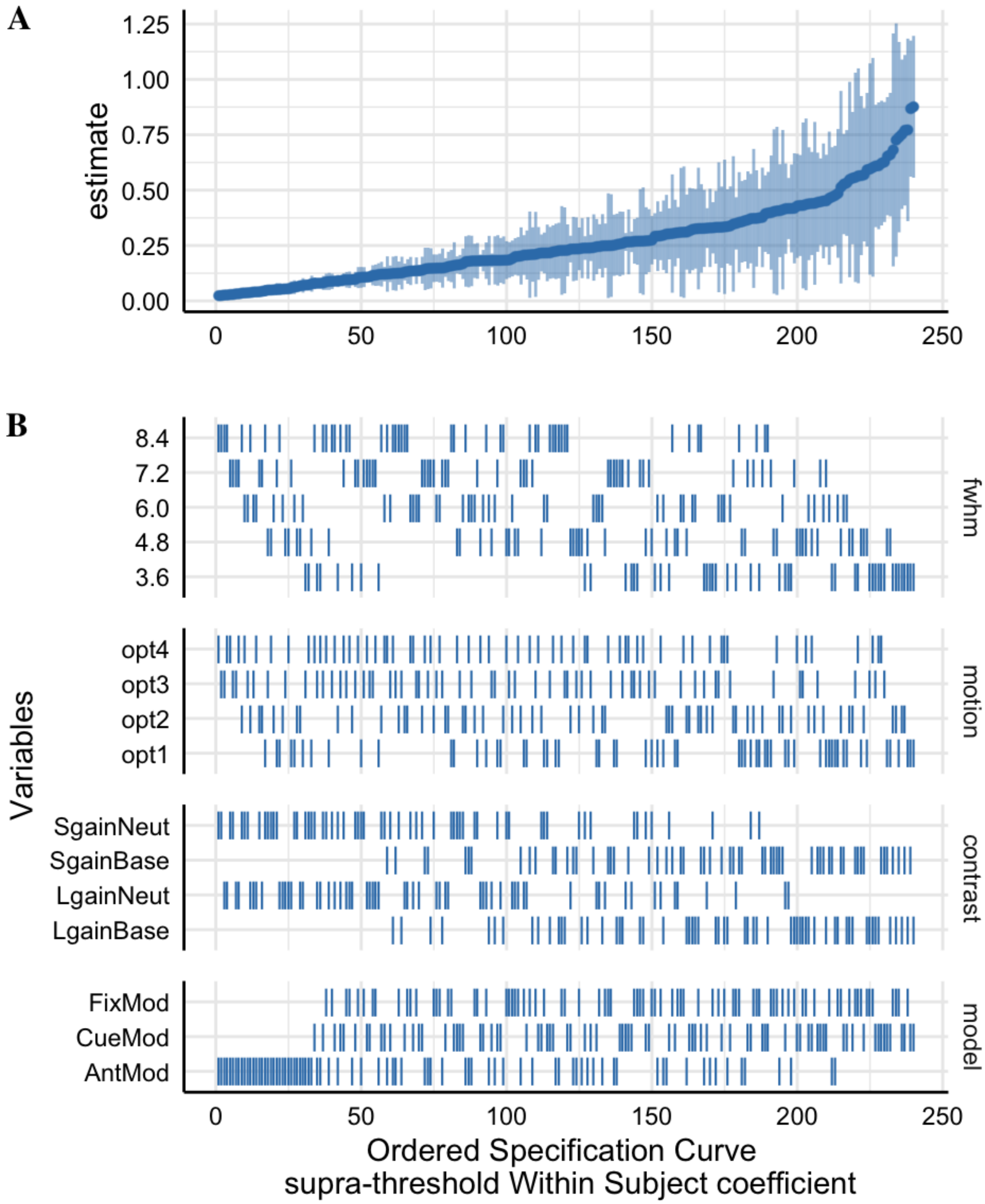


*Figure S35*: Between-session: The supra-threshold Specification Curve of the Median Within-subject variance ($\sigma_v^2$) estimates across 240 pipeline permutations for the ABCD, AHRB and MLS estimate.

A. The distribution of the point estimate (average) across the three studies and distribution across the three samples.

B. The model options (four) associated with each estimate.

### 2.5 Aim 3 results

*Between-Run Stability Effect Size*:

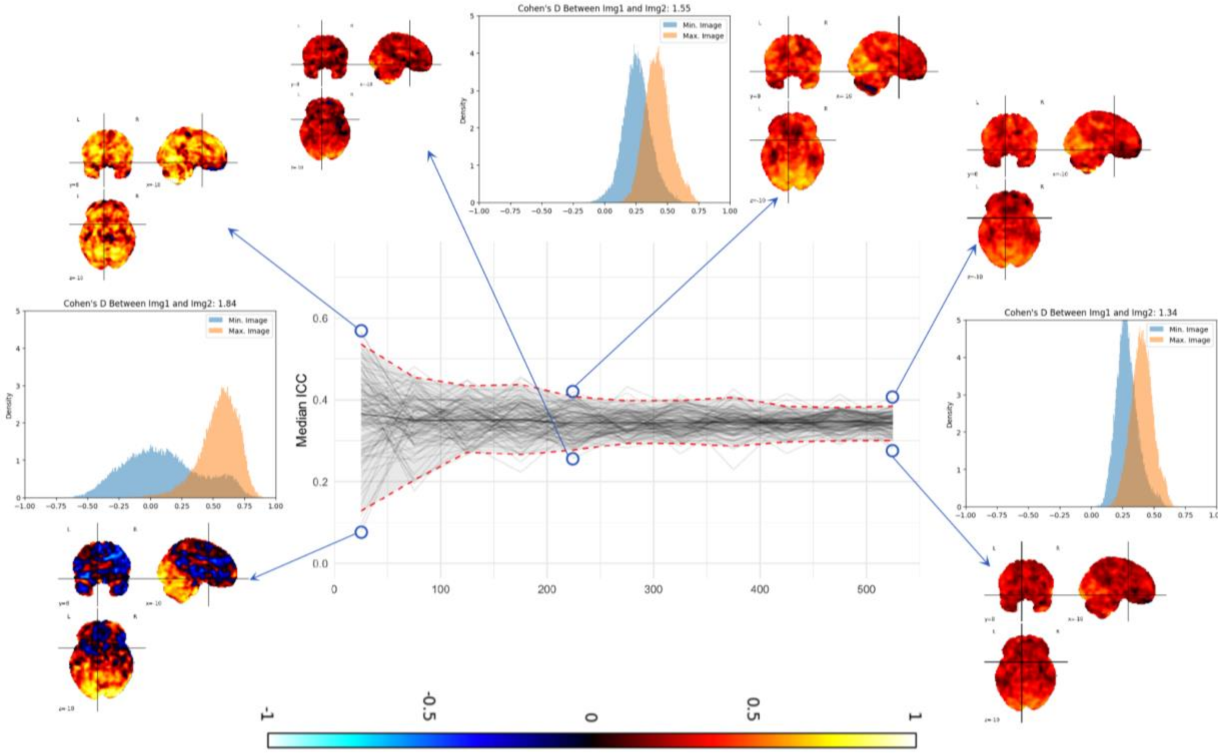


*Figure S36*: Changes in the Median ICC (Supra-threshold mask) estimate in the ABCD sample from *N* 25 to 525 with 100 bootstraps at each *N* for Top Model in Figure 2: *Small Gain* versus *Baseline* Contrast, Cue Model, Motion option 1 and FWHM 8.4. The associated 3D volumes are plotted for the maximum and minimum median ICC value at N 25, 225 and 525 (circled) and associated voxelwise distribution of maps and Cohen's *d* between maps are provided.
*Note:* Upper and Lower dashed red lines: +/- 95% Confidence Intervals for the median estimates; black solid line is the average of the median estimates; light gray lines are individual subsamples, N 25 to N 525, for each bootstrap.

### 2.6 Post Hoc Analyses

*Modeling impacts on Left/Right NAcc*:

#### Effect of analytic decisions on ICC estimate for Left and Right Nucleus Accumbens

For the MID task, researchers are often interested in the activation of the bilateral nucleus accumbens (NAc). The strength of the median ICC estimate from 3D volumes is that it is agnostic to small, anatomical biases and captures the central tendency of ICC estimates across the brain. However, a weakness is that it lacks specificity that is often of interest to brain-behavior researchers. A *post hoc* analysis of the Left and Right NAc was performed using the NAc region of interest from the Harvard-Oxford subcortical atlas (procedure described in Demidenko et al., 2023) for the Session 1 between-run data.

The specification curve and the HLM results are reported for the Left and Right NAc in supplemental **Figure S37** and **Table S16**, respectively. The average ICC estimate across the 240 pipelines varied across the three samples for the *Left NAc* (ABCD = .09 [Min: -0.06, Max: .32]; AHRB = .11 [Min: -.23, Max: .46]; MLS = .17 [Min: .03, Max: .44]) and *Right NAc* (ABCD = .08 [Min: -0.04, Max: .32]; AHRB = .03 [Min: -.25, Max: .42]; MLS = .11 [Min: -.07, Max: .40]). In general, model parameterization had a near zero impact on the ICC estimates for the Left ($\Delta R^2$:  00) and Right NAc ($\Delta R^2$: .01). The analytic decision that explained the largest amount of variance in the ICC estimates is contrast selection for the Left ($\Delta R^2$: .27) and Right Nac ($\Delta R^2$: .24). For example, the change from the contrast of *Large Gain* versus *Implicit Baseline* to *Large Gain* versus *Neutral* results in a $b$ = .01 decrease in the ICC estimate for the Left NAc and $b$ = -02 decrease for the Right NAc. The largest effect on the ICC estimates is the change from the contrast of *Large Gain* versus *Implicit Baseline* to *Small Gain* versus *Neutral* which results in a $b$ = .13 decrease for the Left NAc and $b$ = .10 decrease for the Right NAc estimate. Consistent with the Aim 1a results, for Left NAc and Right NAc, the highest average ICC estimate across the three studies is for the *Small Gain* versus *Implicit Baseline* contrast for the Cue Model with no motion correction and 8.4mm FWHM.

*Table S16*: Hierarchical Linear Model: (A) Linear associations between the analytic decisions and the ICC estimate for Left and Right NAc and (B) the impact of the analytic category on the marginal $R^2$.

**A. HLM Nucleuss Accumbens (NAc) Estimates**

| | Left Nac | | | Right Nac | | |
|---|---|---|---|---|---|---|
| *Predictors* | *b* | *CI* | *p* | *b* | *CI* | *p* |
| (Intercept) | .16 | .11 – .20 | <.001 | .11 | .07 – .14 | <.001 |
| Reference [3.6] | | | | | | |
| fwhm [4.8] | .02 | .00 – .04 | .02 | .01 | -.01 – .03 | .23 |
| fwhm [6.0] | .04 | .02 – .06 | <.001 | .02 | .01 – .04 | .01 |
| fwhm [7.2] | .05 | .03 – .07 | <.001 | .04 | .02 – .05 | <.001 |
| fwhm [8.4] | .06 | .04 – .08 | <.001 | .05 | .04 – .07 | <.001 |
| Reference [opt1] | | | | | | |
| motion [opt2] | -.05 | -.06 – -.03 | <.001 | -.04 | -.06 – -.02 | <.001 |
| motion [opt3] | -.06 | -.08 – -.05 | <.001 | -.06 | -.08 – -.05 | <.001 |
| motion [opt4] | -.07 | -.09 – -.06 | <.001 | -.06 | -.08 – -.05 | <.001 |
| Reference [AntMod] | | | | | | |
| model [CueMod] | .02 | .00 – .03 | .01 | .01 | -.01 – .02 | .27 |
| model [FixMod] | .01 | -.00 – .03 | .05 | .03 | .01 – .04 | <.001 |
| Reference [LgainBase] | | | | | | |
| con [LgainNeut] | -.01 | -.02 – .01 | .28 | -.02 | -.04 – -.01 | .01 |
| con [SgainBase] | .00 | -.02 – .02 | .98 | .03 | .01 – .04 | <.001 |
| con [SgainNeut] | -.13 | -.14 – -.11 | <.001 | -.10 | -.12 – -.09 | <.001 |

**B. Analytic Category Model Impact**

| **Comparison** | **χ2** | **Orig R2** | **New R2** | **ΔR2** | **χ2** | **Orig R2** | **New R2** | **ΔR2** |
|---|---|---|---|---|---|---|---|---|
| [Full] vs [New - fwhm] | 57 | .38 | .34 | .04 | 48 | .36 | .33 | .03 |
| [Full] vs [New - motion] | 91 | .38 | .31 | .07 | 83 | .36 | .30 | .06 |
| [Full] vs [New - model] | 7 | .38 | .38 | .00 | 16 | .36 | .35 | .01 |
| [Full] vs [New - con] | 305 | .38 | .11 | .27 | 260 | .36 | .12 | .24 |

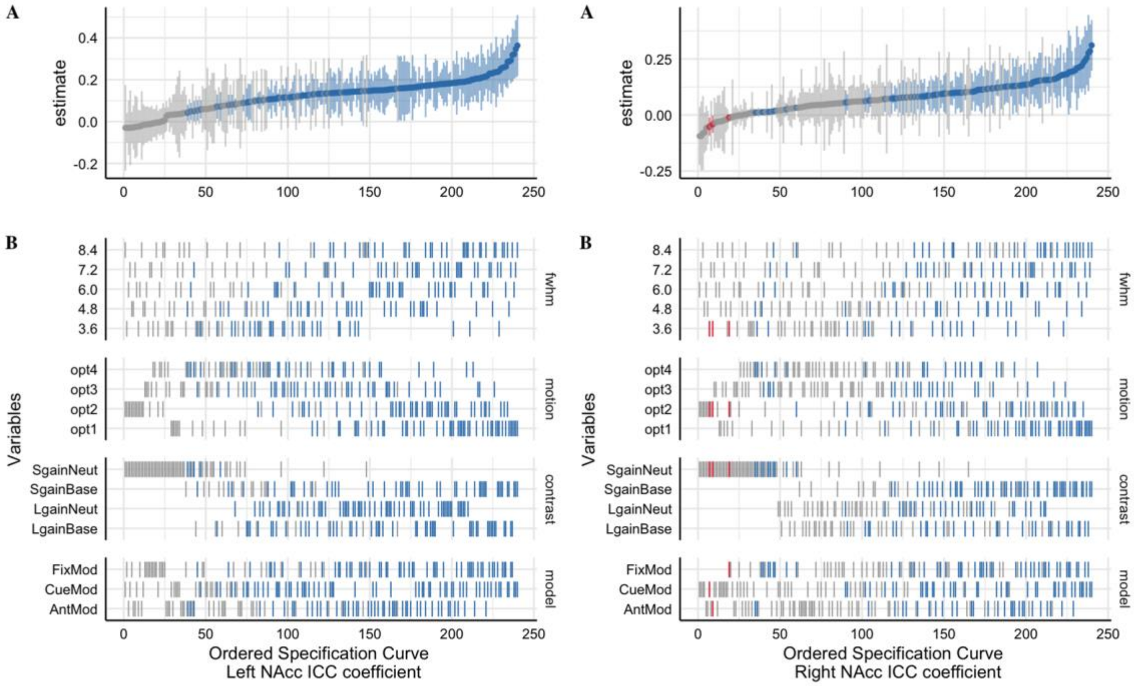


*Figure S37*: The Specification Curve of the ICC estimates for **left** and **right** NAcc across 240 pipeline permutations for the ABCD, AHRB and MLS samples.
A. The distribution of the point estimate (average) across the three studies and distribution across the three samples.
B. The model options (four) associated with each estimate.

### Group-level Cohen's *d* association with estimated ICC

Given the potential association between estimated ICCs and group-level activations magnitudes, the correlation between run and session maps was evaluated for the supra-threshold mask using Spearman *rho*. Across the 240 pipeline permutations, the *rho* coefficient between Session 1 group-level Cohen's *d* maps and Session 1 between-run ICC maps are low on average but vary widely for *Run 1* (ABCD = -.05 [Min: -.43; Max: .22]; AHRB = .09 [Min: -.41; Max: .50]; MLS = .08 [Min: -.35, Max: .43) and *Run 2* (ABCD = -.04 [Min: -.47; Max: .26]; AHRB = .10 [Min: -.40; Max: .51]; MLS = .08 [Min: -.38, Max: .46). This pattern is consistent for the session-level estimates, whereby the associations between the session group-level maps and the between-session ICC maps are low on average but vary widely for *Session 1* (ABCD = .01 [Min: -.40; Max: .29]; AHRB = .11 [Min: -.45; Max: .53]; MLS = .12 [Min: -.28, Max: .43) and *Session 2* (ABCD = -.01 [Min: -.46; Max: .30]; AHRB = .12 [Min: -.43; Max: .53]; MLS = .11 [Min: -.31, Max: .39]).
